# Rigid Control of Motor Unit Firing Rates in the Human Tibialis Anterior Muscle Persists during Neurofeedback

**DOI:** 10.1101/2025.02.10.637074

**Authors:** Meng-Jung Lee, Patrick Ofner, Hsien-Yung Huang, Joris Mulder, Jaime Ibanez, Dario Farina, Carsten Mehring

## Abstract

The conventional framework of motor-unit (MU) control assumes that MUs in a MU pool are constrained by a fixed recruitment order and a common input. This rigid-control framework has been challenged by recent findings suggesting that MU activity could be flexibly modulated, potentially mediated by descending cortical inputs. In this study, rather than evaluating flexibility from the perspective of recruitment thresholds, we investigated control flexibility by assessing if human participants can voluntarily modulate MU firing rates beyond rigid-control constraints. Specifically, we examined whether participants could voluntarily modulate the firing rates of a pair of MUs from the tibialis anterior muscle during real-time feedback. Two tasks involving target-reach with different visual feedback derived from the MUs firing rates were conducted. In both tasks, there was no evidence that participants were able to change MU firing rates in a way that would violate rigid control robustly. Our findings demonstrate limited flexibility in MU control in human tibialis anterior muscle within single-session training, even when real-time MU activity feedback was provided. The results suggest that MU flexibility is not inherently present in the human lower limb.

## Introduction

Motor units (MUs) are fundamental functional units through which the nervous system controls movement [1]. Flexible MU control could potentially provide a control signal for neural interface applications with more than a single degree of freedom per muscle [2]. A non-invasive approach has been developed to decompose high-density surface electromyography (HD-sEMG) signals, enabling the real-time acquisition of multiple MU activities simultaneously [3-6]. Despite the potential numerous degrees of freedom (DoF) available with MUs, studies have shown that the central nervous system employs various strategies to reduce its computational load [7-9], effectively limiting the DoF. MUs are orderly recruited based on their intrinsic neural properties according to the Henneman’s size principle [10-12], which has been observed robustly in various muscles [13-17]. Additionally, activities of MUs within the same functional group [18-20] are highly correlated due to the large portion of common input they receive [21-24]. Such a rigid control scheme proposes that the activity of each MU changes in a predictable way dependent on the common input. In this framework one would expect that the firing rates of any two MUs in a single-compartment muscle within the same pool are confined to a one-dimensional curve (a one-dimensional manifold) in the two-dimensional space spanned by their firing rates (state space) [25], imposing constraints on the voluntary control of the individual MUs [26, 27].

Though the recruitment order of MUs appears to be fixed, reversals in recruitment order can be observed in certain cases such as during fast force changes [25, 28, 29]. Selective activation of single MUs at different recruitment thresholds within the same muscle has also been reported with biofeedback training [30-32]. Furthermore, in macaque monkeys, cortical micro-stimulation and MU activity during dynamic force profiles revealed deviations from activity patterns typically observed during slow force ramp tasks, consistent with the presence of multiple descending neural pathways enabling selective cortical control of MUs [25]. However, a recent study in humans showed that participants could only achieve selective control of MUs within a muscle by leveraging the recruitment-derecruitment hysteresis, without providing any evidence for voluntary selective control of the input to MUs [26]. Given that recruitment order can be influenced by factors such as contraction speed [25], muscle length [33] and posture [34], a change of the recruitment order do not necessarily indicate flexible control of individual MUs [1, 28]. Therefore, a more robust approach, such as assessing firing rate modulation while excluding recruitment and derecruitment considerations, is desirable to determine whether MUs can be flexibly controlled. However, as of yet it remains unclear whether MU firing rates can be voluntarily and flexibly modulated while they are steadily active, i.e., recruited.

In this study, we, therefore, investigated the flexibility in controlling MU firing rates in the human tibialis anterior muscle. Unlike previous studies that examined changes in recruitment order, we focused exclusively on continuously recruited MUs to avoid the effect of MU recruitment and derecruitment. If MUs can be controlled independently through descending neural signals, their firing rates should not be restricted monotonically to the common drive during activation regardless of their recruitment thresholds. To assess flexible MU control, we borrowed the concept of ‘displacement zone’ introduced by Marshall et al. [25] (Fig. 2a). The term ‘displacement zone’ refers to the deviation of the firing rates of two MUs from a one-dimensional manifold within the two-dimensional state space. Firing rates of MUs can only fall within the displacement zone if the firing rates of the two MUs change simultaneously in opposite directions. Such behavior cannot be achieved through one source of common input and thus would indicate flexible MU control.

Two single-session target-reaching tasks, namely displacement control and difference control, were conducted. Real-time visual feedback of the MU firing rates was provided via a cursor on a computer screen (Fig. 1), with each task employing a distinct mapping of firing rates to cursor movement. In the displacement control task, the cursor position along the X/Y-axes was controlled by the firing rates of the MU pair. In the difference control task, participants were provided with a less restrictive control condition with an alternative mapping of MU firing rates to the cursor position. No evidence was found to support that participants could consistently modulate MU firing rates into the displacement zones in either task.

**Figure 1.**
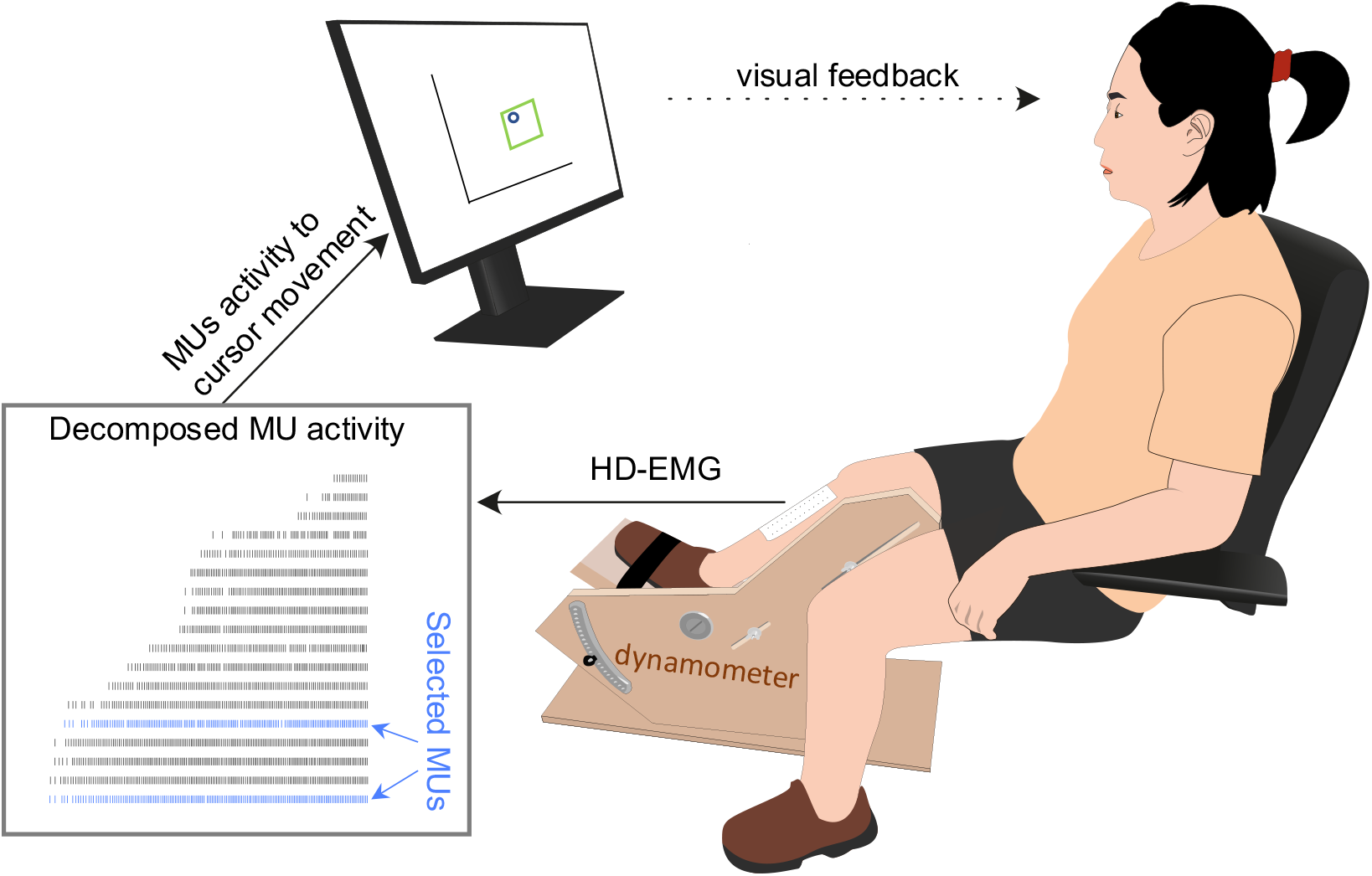
Schematic overview of the experimental setup. The participant’s foot from the dominant leg was secured to the dynamometer with Velcro. A one-dimensional force sensor on the dynamometer measured the force exerted by the dorsiflexion movement. High-density surface electromyographic signals from the tibialis anterior muscle were obtained and decomposed into MU activities in real-time. A pair of MUs were selected as controlled MUs. The activities of the controlled MUs were translated into cursor movement that was presented to the participant on a screen.

## Results

### Real time displacement control with motor unit activities

Under the hypothesis of rigid control, the firing rates of any two recruited MUs would either increase or decrease simultaneously. Fig. 2a illustrates the firing rates (FRs) of an MU pair under rigid control in the state space. The firing rates follow a one-dimensional manifold (black curve) as from r_t_ to r_t1_. The green zones represent the possible firing rate combinations passing through r_t_ which are consistent with the rigid control hypothesis. Conversely, under the flexible control hypothesis, the firing rates of MUs can be any combination and are not restricted to the green zones. In the state space, firing rates that pass through r_t_ but violate the rigid control hypothesis are referred to as ‘displacement zones’, represented by the red-colored areas in Fig. 2a., such as in r_t2_ [25]. If participants could volitionally and consistently modulate firing rates of two MUs used for feedback to fall within the displacement zones, it would suggest that they were able to independently control the two units and that those MUs were receiving inputs changing in opposite directions.

**Figure 2.**
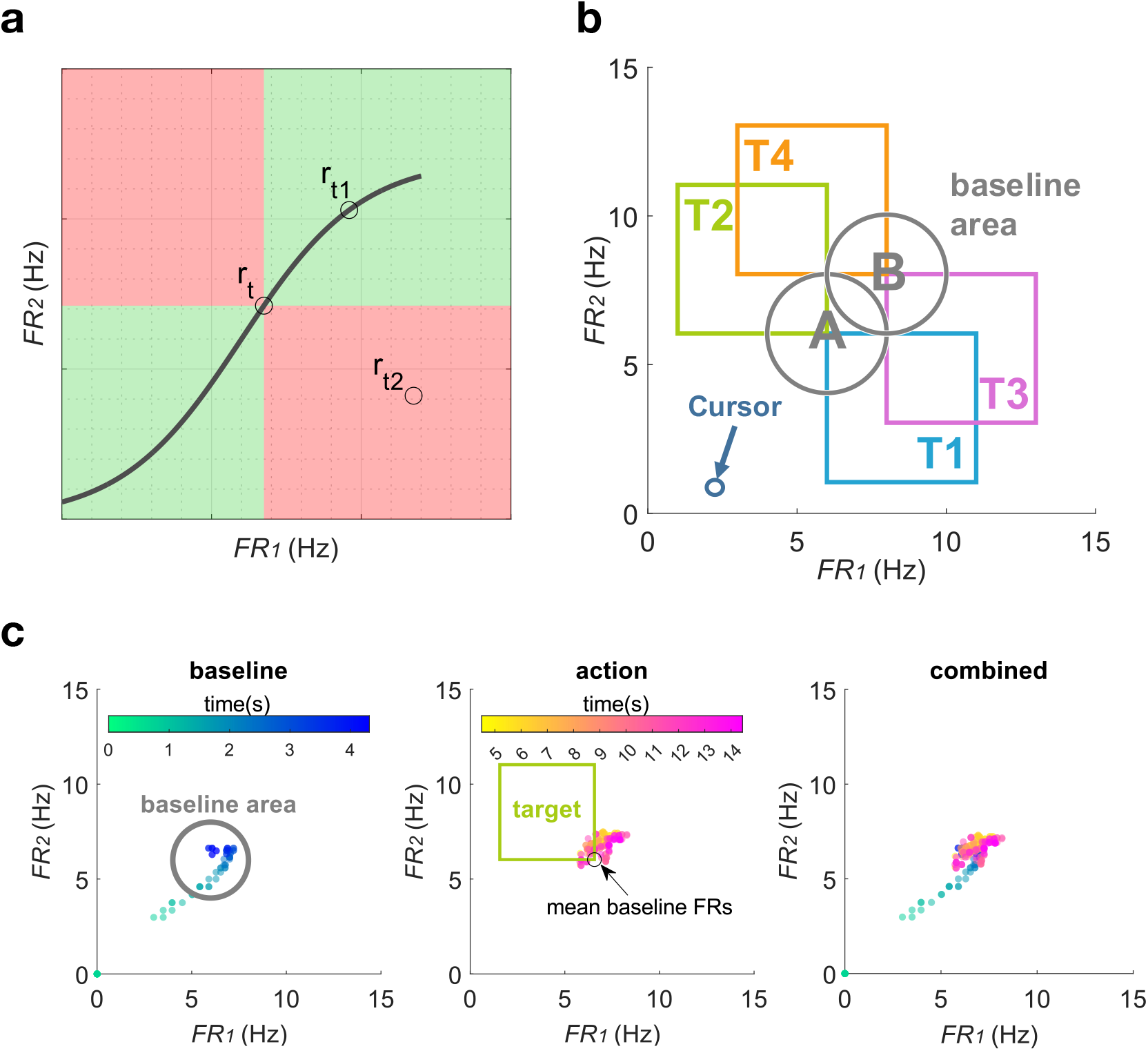
(a) Schematic illustration of displacement zones (red) and a one-dimensional manifold (black line) for two MUs under the rigid control hypothesis. Figure adapted from Fig.3a of Marshall et al. (2022). (b) Illustration of the displacement control task. T1-T4 refer to each individual targets. (c) Cursor trajectory from an example trial. Left: Cursor trajectory from the beginning of the trial to the end of the 3 s holding period over the baseline area (grey). Middle: Cursor trajectory during the 10 s action phase. The target position was calibrated with the averaged FRs from the two MUs during the baseline phase. The color depicts the time passed since the beginning of the trial. Right: Cursor trajectory during the entire trial. The cursor position was updated every 62.5 ms. FR_1_ and FR_2_ correspond to the firing rates of MU_1_ and MU_2_, respectively. In the displacement control task, firing rates of MU_1_ was mapped to cursor movement along the x-axis, while MU_2_ was mapped to the movement along the y-axis.

In the displacement control task, targets were positioned within the displacement zones. The single-session experiment included 5 blocks, with each block comprising 40 trials, for a total of 200 trials. Each trial consisted of two phases: the baseline phase and the action phase. A trial began with the baseline phase, during which the cursor was held over one of the two baseline areas (Fig. 2b, grey circles) for 3 consecutive seconds. The two baseline areas A and B corresponded to two distinct force levels. The baseline phase was followed by the action phase, where participants were required to navigate the cursor to one of the target areas T1 to T4. T1 or T2 were presented after baseline area A, and T3 or T4 were presented after baseline area B. In each trial, only one baseline area and one target were displayed. Importantly, the target position was placed within a displacement zone relative to the average firing rates of the two MUs during the baseline phase, e.g., r_t_ in Fig. 2a. Fig. 2c shows the cursor trajectory of an example trial.

The mean recruitment and derecruitment thresholds for the two controlled MUs were 2.1±1.2 % MVC (mean ± std. dev.) and 2.7±1.4 % MVC for *MU*_1_, and 2.6±1.7 % MVC and 2.6±2.0 % MVC for *MU*_2_, respectively (Fig. 3a, Bayesian Wilcoxon signed-rank test BF_10_ =1.7 for recruitment and BF_10_ = 0.37 for derecruitment thresholds). We preferentially selected MUs with low recruitment thresholds to reduce the risk of fatigue. The dorsiflexion forces exerted by the participants during the action phase remained stable across the experimental blocks (Fig. 3b, trend-BF_10_ = 0.4, see Methods for details on trend-BF_10_), and there was evidence for the force being dependent on the target (Fig. 3c, Bayesian repeated measures ANOVA, BF_inclusion_ > 100, BF_inclusion_ =4 and BF_inclusion_ =0.4 for target, target – phase interaction and phase, respectively). Average force levels for T3 and T4 were slightly higher than for T1 and T2, regardless of the trial phase (see Supplementary Table 12 for results of post hoc tests).

**Figure 3.**
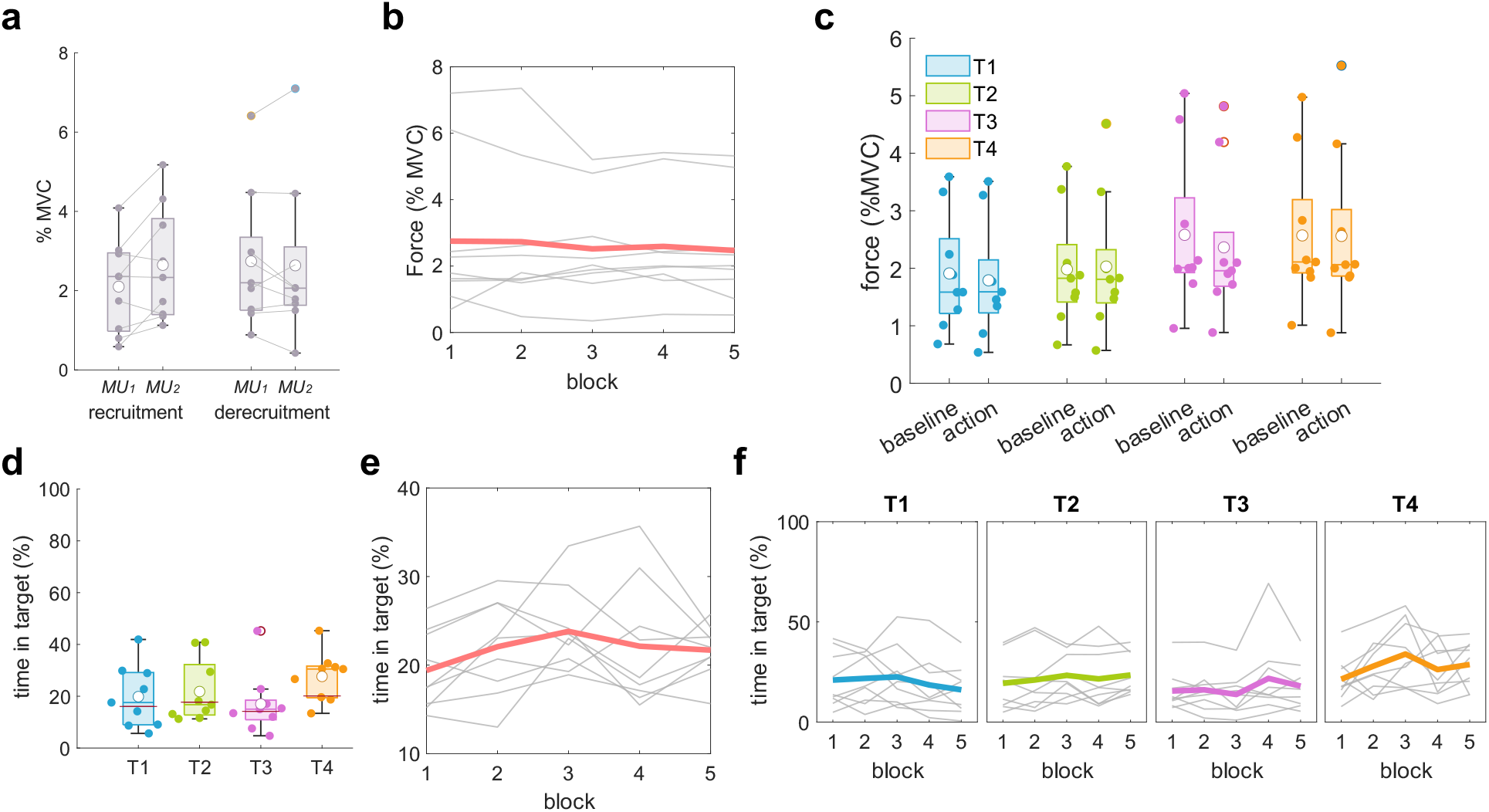
(a) Recruitment and derecruitment thresholds of the two MUs used to control the cursor. (b) Force produced across the five blocks during the action phase (individual participants in gray; mean across participants in red). (c) Force produced during baseline and action phases. (d) Percentage of time the cursor was held inside targets separately for each target. Horizontal dark red bars depict the chance level for each target. Chance levels were determined by randomly reassigning T1 and T2, as well as randomly assigning T3 and T4 for 1000 repetitions. In (c) and (d): each colored dot depicts one participant; white dots depict the mean; horizontal lines the median. (e) Percentage of time the cursor was held inside the targets across blocks for individual participants (gray) and mean across participants (red). (f) Percentage of time the cursor was held inside targets across blocks for each target. Individual participants in gray; means across participants in color.

The performance was evaluated by the percentage of time the cursor was held within the target during each trial. Overall, performance was low, with the average varying between 17% and 28% (Fig. 3d) across targets and was only slightly higher than chance level (Fig. 3d, dark red bars). There was no evidence for a target-dependent performance (Fig. 3d, Bayesian repeated measures ANOVA BF_inclusion_ for target =0.5). Furthermore, trend analysis on the slope of the performance across blocks showed no tendency of improvement in performance across blocks (Fig. 3e, trend-BF_10_ =0.6), or a target-specific learning effect (Fig. 3f, Bayesian repeated measures ANOVA of trends, trend-BF_inclusion_ =0.9 for target).

### Restricted modulation of motor unit firing rates in the displacement control task

We compared the firing rates of both controlled MUs between baseline and action phases in the displacement control task to detect deviations from the one-dimensional manifold. Here, *ΔFR*_1_ and *ΔFR*_2_ are defined as

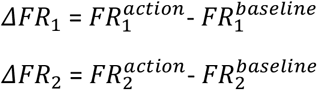

where 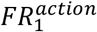 refers to the mean firing rate during the action phase for *MU*_1_, and 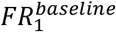 refers to the mean firing rate during the baseline phase for *MU*_1_. The same applies to *MU*_2_. The following three hypotheses were evaluated using Bayesian hypothesis testing (see Methods for details):

*H*_0_: No change in firing rates.

*H*_1_: Successful target reach.

*H*_2_: Firing rate changes not in line with the target area but at least one of the MUs changed its firing rate. *H*_2_ is the complement of *H*_0_ and *H*_1_.

Pooled data from all blocks and participants provided strong evidence against participants being able to modulate the FRs of the controlled MUs towards successful target reach for T1, T2 and T3 and moderate evidence against target reach for T4 (Fig 4. top-right panel and Table 1). Participants did change the firing rates during action phase as compared to baseline phase (BF_20_>10 for all targets), but not towards the target area (BF_21_ > 10 for T1-3 and BF_21_ = 7.2 for T4). For the analysis of individual blocks, anecdotal evidence for successful target reach was observed only in block 3 for T4 (Supplementary Table 1; Fig. 4). No other blocks and targets showed evidence for successful target reach. These findings obtained from the MU firing rates were consistent with the low performance (Fig. 3d) and the lack of improvement in performance across blocks (Fig. 3e-f).

**Table 1:**
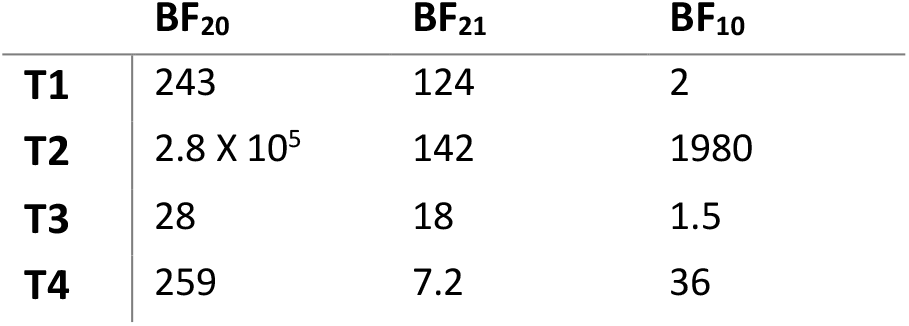
Bayes factors (BFs) quantifying relative strength of evidence for H_0_, H_1_ and H_2_ separately for targets T1 to T4 (see Methods).

|  | **BF<sub>20</sub>** | **BF<sub>21</sub>** | **BF<sub>10</sub>** |
| --- | --- | --- | --- |
| **T1** | 243 | 124 | 2 |
| **T2** | $2.8 \times 10^5$ | 142 | 1980 |
| **T3** | 28 | 18 | 1.5 |
| **T4** | 259 | 7.2 | 36 |

**Table 2:** Bayes factors (BFs) quantifying relative strength of evidence for H_0_, H_1_ and H_2_ for targets from opposing quadrants.

|  | **BF<sub>20</sub>** | **BF<sub>21</sub>** | **BF<sub>10</sub>** |
| --- | --- | --- | --- |
| **T1 & T2** | $5.9 \times 10^5$ | $1.9 \times 10^3$ | 304 |
| **T3 & T4** | $1.3 \times 10^5$ | 63 | $2 \times 10^3$ |

**Figure 4.**
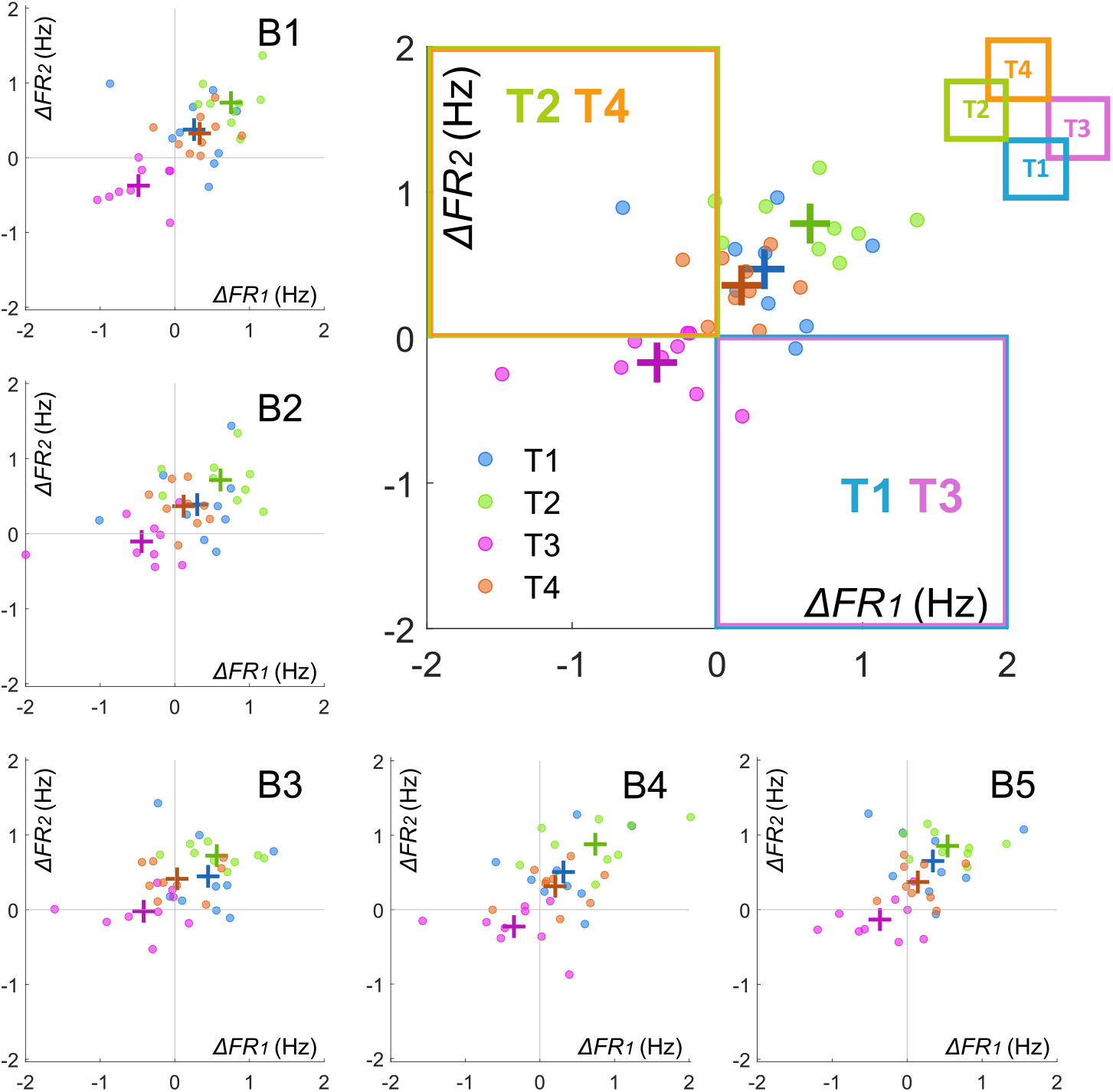
Averaged ΔFR_1_ and ΔFR_2_ across the whole experiment (top-right subfigure), as well as separately for block 1 to block 5, indicated as B1 to B5 (dots: individual participants with color indicating the target; +: target-specific mean across participants). Colored rectangular boxes label the target areas.

We assessed the variability of firing rate differences between baseline and action phases for each individual participant (Fig. 5a, b). Firing rate variability was quantified using the standard deviation of *ΔFRdiff* and *ΔFRsum*, defined as follows:

**Figure 5.**
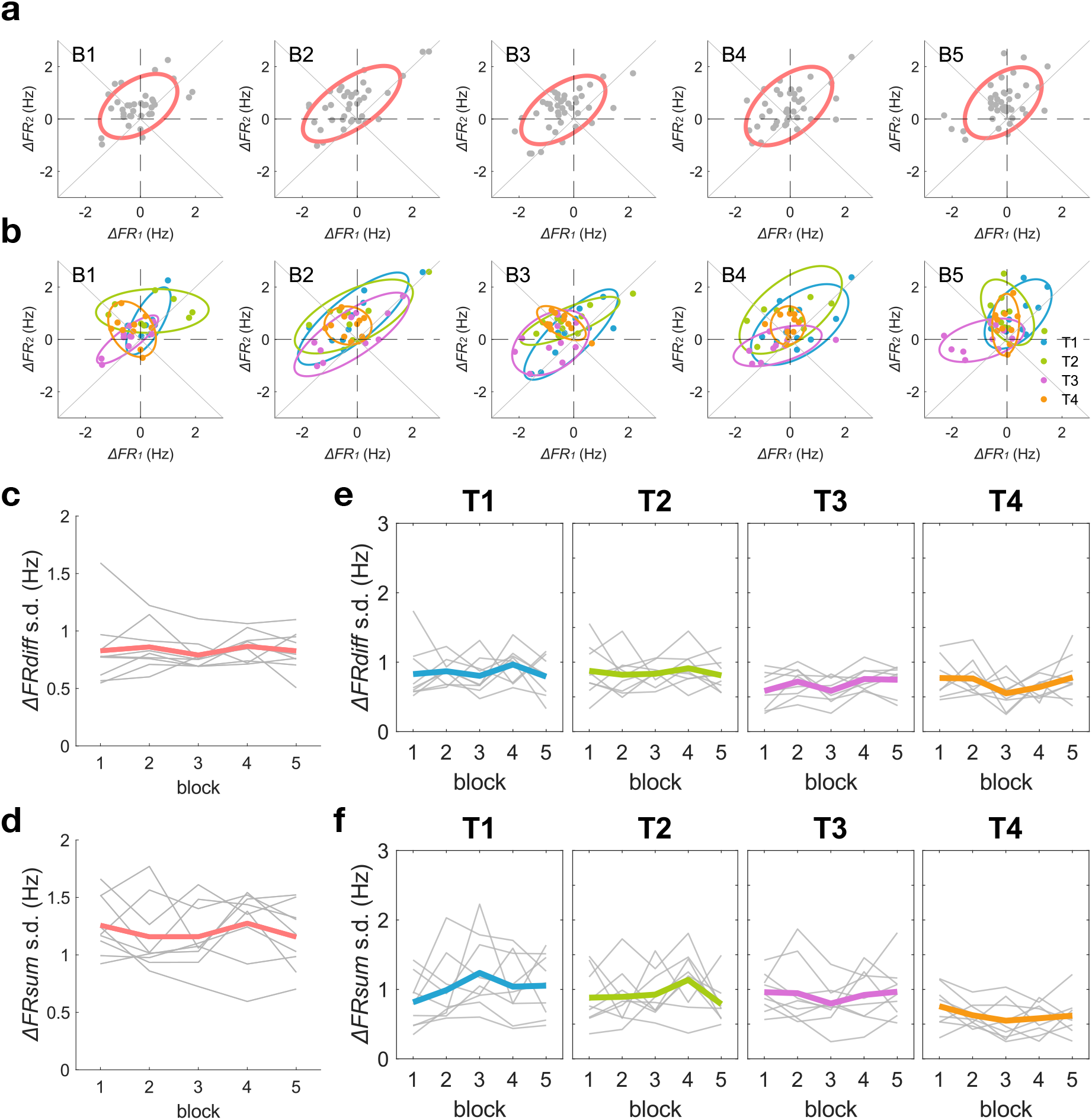
(a-b) Distribution of ΔFR_1_ and ΔFR_2_ from a single participant across blocks. Data pooled across all targets (a, gray dots: single trials) and separately for each target across blocks (b). Ellipses visualize bivariate Gaussian distributions fitted to firing rate differences, with contour lines containing 80% of the data probability. B1 to B5 indicate block 1 to block 5. (c-d) Standard deviation of the difference (c) and the sum (d) of ΔFR_1_ and ΔFR_2_ (Individual participants in gray; mean across participants in red). (e-f) Same as (c) and (d) but separately for targets T1 – T4.

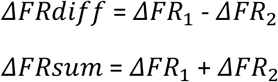

The standard deviation of *ΔFRsum* captures the variability in the firing rates of the two MUs along the diagonal from the origin towards the upper right in the state space (Fig. 5a) and is expected to correlate with the variability of the exerted force, while the standard deviation of *ΔFRdiff* quantifies variability in firing rates perpendicular to this diagonal. There was no indication that either of these variabilities changed across blocks (Fig. 5c,d, trend-BF_10_ = 0.4 and 0.3 for c and d, respectively), and also not in a target-specific way (Fig. 5e,f, Bayesian repeated measures ANOVA of trends, trend-BF_inclusion_ for target equal to 0.6 and 0.7 for e and f, respectively).

We further analyzed target reach in individual participants in the displacement control task. Four out of ten participants (including the one participant excluded from the preceding population analysis, P28) showed moderate to strong evidence for target reach for one or two targets (Fig. 6a). Figure 6b shows the *ΔFR*_1_ and *ΔFR*_2_ for each individual trial across the four targets for a single participant (P23), who demonstrated evidence for target reach for T2 and T4 but not for T1 and T3. Note that T1 and T3 belong to one quadrant, while T2 and T4 belong to a different, but also shared, quadrant. Such single-quadrant target reaching was observed in all four participants who successfully reached individual targets. Although four participants demonstrated successful reach of individual targets, none were able to reach opposing targets corresponding to the same baseline area, such as T1 and T2, or T3 and T4. This result suggests that successful target reach was unlikely achieved through flexible MU control, as participants would likely have been able to navigate the cursor also to targets in the opposite quadrant if such control had been present.

**Figure 6.**
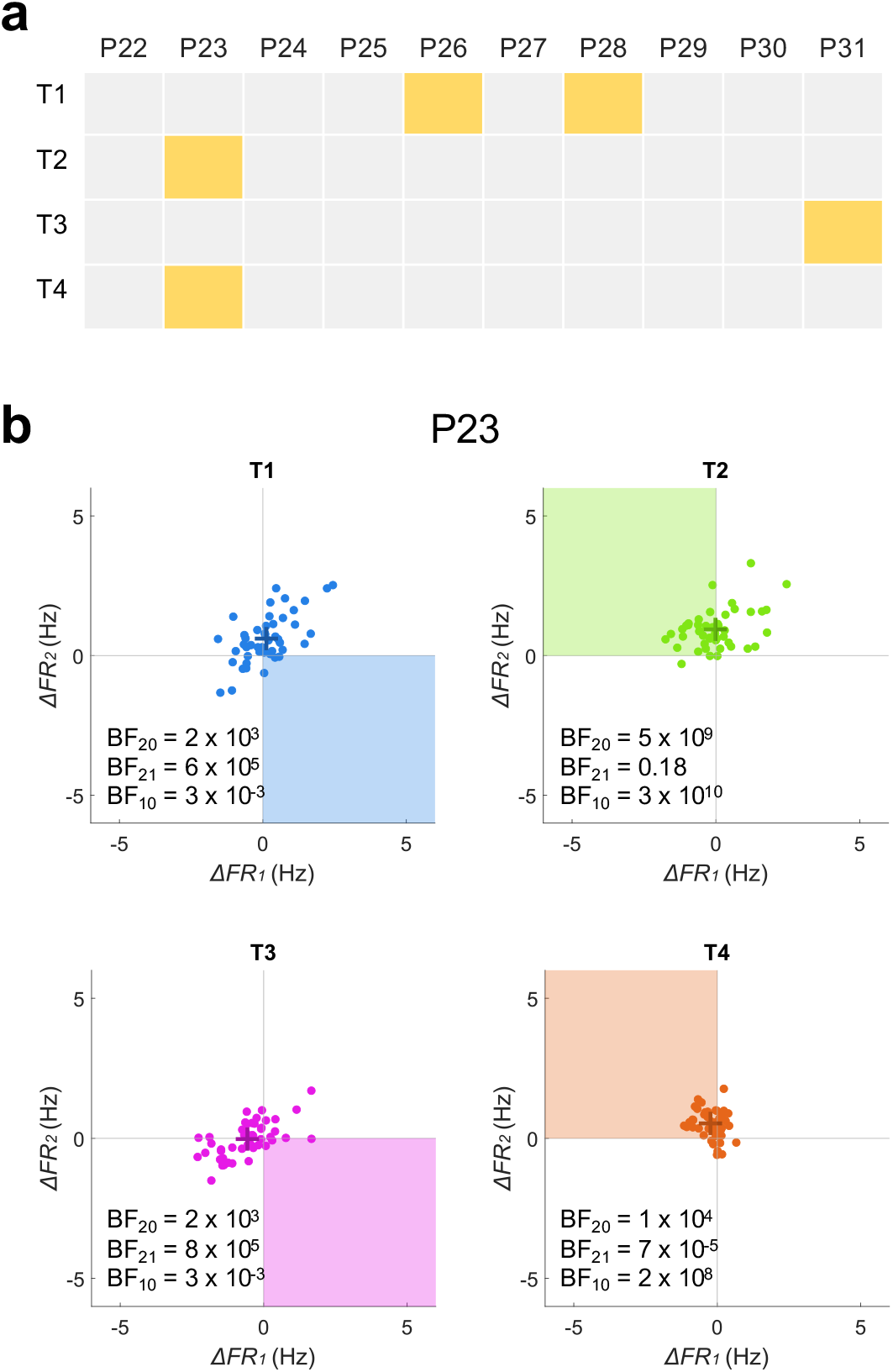
(a) Distribution of successful target reach for each participant (P22-P31). Cases with moderate or strong evidence in favor of successful target reach are marked yellow. The BFs can be found in Supplementary Table 2. (b) ΔFR_1_ and ΔFR_2_ across the whole experiment separated by targets for one participant (P23, dots: individual trials; +: mean across all trials from the target). Colored rectangular areas label the target zones for the individual target.

To test on the population level whether there was evidence for successful target reach to targets in opposite quadrants, we modified the test hypotheses from the previous section. *H*_1_ now considers both opposing targets:

*H*_0_: No change in firing rates.

*H*_1_: Successful target reach for targets in opposite quadrants, i.e., either T1&T2 or T3&T4.

*H*_2_: Firing rate changes not in line with the target area but at least one of the units changed its firing rate. *H*_2_ is the complement of *H*_0_ and *H*_1_.

On the population level, our results provide strong evidence against successful reach of opposing targets (BF_21_ > 10 and BF_20_ > 10 for both T1&T2 and T3&T4).

### No displacement was observed under less constrained control conditions

In a second experiment, referred to here as the difference control task, we investigated whether participants could modulate firing rates of pairs of MUs violating rigid control under a less restrictive control condition. In this task, the vertical cursor position was controlled by the sum of the firing rates of two MUs, while the horizontal cursor velocity was controlled by the difference of their firing rates. In this task, the targets were placed at both ends of the horizontal axis (Fig. 7a). Since participants only had to generate differences in the firing rates to reach the targets, the areas in the state space relevant to successful target reach included the respective halves of the state space (Fig. 7b). These areas included the displacement zones described in the previous section, as well as areas outside the displacement zones, as illustrated in Fig. 7b. A threshold on the firing rate differences was applied, below which the horizontal velocity was set to zero (see Methods for details). Participants first moved the cursor into the baseline area and held it there for 1 consecutive second (baseline phase), with the horizontal position fixed. After this baseline phase, the trial proceeded to the action phase, during which the cursor could move freely horizontally and vertically. A trial was considered successful if a participant could hold the cursor at the target for 3 consecutive seconds. Otherwise, the trial failed with a timeout after 30 seconds. The single-session task consisted of 5 blocks, with a total of 180 trials. Figures 7c-d show the firing rates of the two controlled MUs and the cursor trajectory over time from an example trial.

**Figure 7.**
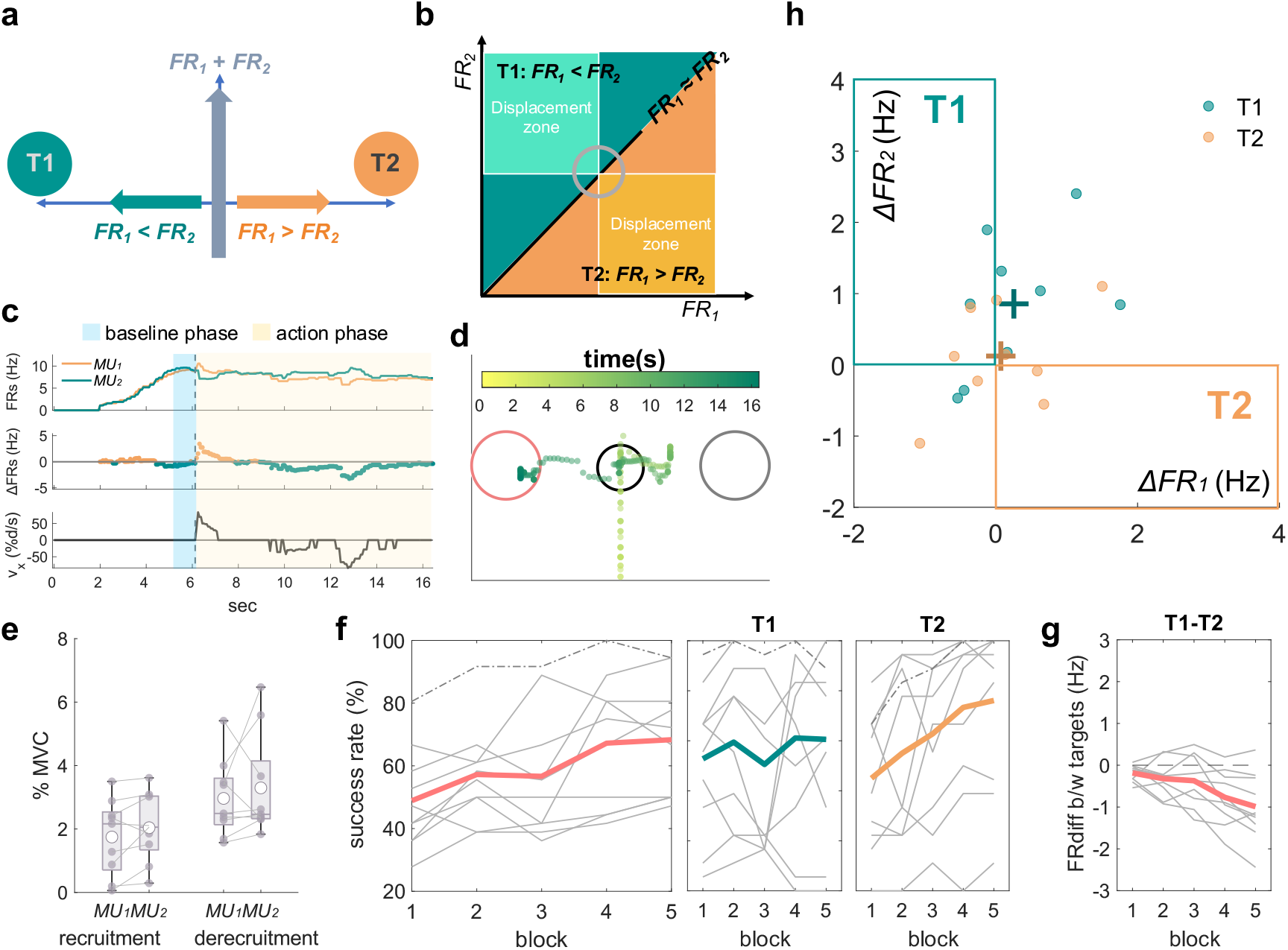
(a) Illustration of the control scheme. The cursor position along the vertical axis was controlled by the sum of the firing rates of the two MUs, and the cursor velocity along the horizontal axis was controlled by the difference between their firing rates. (b) Illustration of firing rate areas in the state space relevant for target reach. Each area is subdivided into regions within and outside the displacement zone. The baseline area is depicted by the grey circle. (c-d) Firing rates of the two MUs, horizontal velocity (c) and cursor trajectory (d) from an example trial. Middle panel in (c): ΔFRs refers to the difference between FR_1_ and FR_2_, with color coding indicating which MU exhibits a higher firing rate. Bottom panel in (c): velocity of the cursor movement in percentage of the total distance from the baseline area to the target per second. In (d) the black circle indicates the baseline area. The red circle represents the active target, while the grey circles represent the inactive target. (e) Recruitment and derecruitment thresholds of the two MUs used to control the cursor. (f) Success rate across the 5 control blocks. Left: combined T1 and T2; middle and right: separated for T1 and T2 (individual participants in gray; mean across participants in color). The data for the single participant whose neural data was excluded (see Methods) is represented by a dash-dot line. (g) Firing rate differences between the targets across 5 blocks during the action phase. FRdiff b/w targets (with ‘b/w’ denoting ‘between’) here refers to 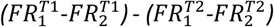 during the action phase, where 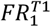 and 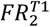 refer to averaged firing rates of MU_1_ and MU_2_, respectively, in trials with T1 (the same applies to T2 trials). (h) Mean ΔFR_1_ and ΔFR_2_ for individual participants (dots) and mean across participants (crosses). Mean ΔFR_1_ and ΔFR_2_, similar to the displacement task, were defined as the average firing rate differences between the action phase and the baseline phase. Colored rectangular boxes label the displacement zones for the two targets. FR represents the firing rate.

The mean recruitment and derecruitment thresholds were 1.7±1.2 % MVC and 3.0±1.2 % MVC for *MU*_1_ and 2.0±1.1 % MVC and 3.3±1.6 % MVC for *MU*_2_ (Fig. 7e, Bayesian Wilcoxon signed-rank test BF_10_ =1.98 for recruitment and BF_10_ = 0.51 for derecruitment thresholds). The results showed that participants modulated firing rate differences, and this modulation increased across blocks in line with the increased performance (Fig. 7f-g, f left: trend-BF_10_ >100, f middle and right: trend-BF_10_ =0.3 and 360 for T1 and T2, g: trend-BF_10_ = 40, Supplementary Fig. S1, S2a). The observed firing rate differences, however, do not necessarily imply deviations from the one-dimensional manifold (Fig. 2a), as these differences could be within or outside of the displacement zones (Fig. 7b). To clarify this point, we conducted an analysis comparing the firing rate changes between the baseline and action phases using the same three hypothesis tests as in the displacement control task.

There was no evidence for firing rates modulation towards the displacement zones of the corresponding targets for data aggregated across blocks (Fig. 7h and Table 3) in line with results from the analysis of individual blocks (see Supplementary Fig. S3 and Supplementary Table 3). Furthermore, there was strong evidence against successful target reach to both opposing targets (Table 4). Taken together, these results suggest that participants were able to learn the task under less strict control conditions.

**Table 3.** Bayes factors (BFs) quantifying relative strength of evidence for H_0_, H_1_ and H_2_ in the difference control experiment. Same hypotheses as in Table 1. The data provides inconclusive evidence for T1 and strong evidence forH_0_ for T2, i.e., no evidence for flexible MU control.

| | $BF_{20}$ | $BF_{21}$ | $BF_{10}$ |
| --- | --- | --- | --- |
| **T1** | 2.6 | 0.9 | 2.8 |
| **T2** | 0.18 | 1.4 | 0.1 |

**Table 4:** Bayes factors (BFs) quantifying relative strength of evidence for H_0_, H_1_ and H_2_ for trials from opposing targets. Same hypotheses as in Table 2.

| | $BF_{20}$ | $BF_{21}$ | $BF_{10}$ |
| --- | --- | --- | --- |
| **T1 & T2** | 29.1 | 129.6 | 0.23 |

Moreover, participants improved performance within a single-session training by increasing the difference in firing rates between the two MUs across blocks. However, the firing rate differences were not produced by modulating the firing rates of the MUs into the corresponding displacement zones.

## Discussion

### Lack of flexibility in MU control

The possibility of flexible MU control remains debated due to contradictory findings regarding the selective recruitment of MUs, and little is known about whether individuals can modulate the activity of two MUs independently. In this study, we investigated whether the activity of two MUs from the same pool can be voluntarily modulated in a way that the firing rate is increased in one unit and decreased in the other. Demonstrating this capability would provide compelling evidence for the human ability to selectively control individual MUs within a pool of a single muscle. We focused solely on flexible firing rate control without allowing for recruitment/derecruitment strategies and employed two tasks with different degrees of motor constraints. On the population level, our results from a group of participants revealed no evidence that MU firing rates could deviate from rigid control. Nevertheless, individual participant analysis showed that four out of ten participants in the displacement control task demonstrated moderate to strong evidence of successful individual target reach. However, it is important to note that no participant was able to reach opposing targets. When two targets were reached, they were located within the same quadrant relative to the baseline area. The inability to reach opposing targets suggests a lack of flexible MU control, as participants should have been able to voluntarily navigate the cursor to both opposing targets if MUs were flexibly controlled by independent inputs. Instead, target reach appeared to be affected by factors that consistently drove the firing rates toward the same quadrant. This outcome may be attributed to intrinsic properties of MUs in their response to synaptic inputs, such as the influence of persistent inward currents (PICs). One effect of PICs is the hysteresis in firing rates when MUs receive a triangular pattern of increasing and decreasing input [35]. When the two controlled MUs exhibited different ascending and descending slopes in their firing rates hysteresis, it can result in a positional shift from the baseline. By first increasing and then decreasing the exerted force, participants could have potentially guided the cursor into the target area as illustrated in Fig. 8. Depending on the relative slopes of the two MUs, participants could have reached either T1 and T3 or T2 and T4 in the displacement control task. However, such a control strategy would not allow the reaching of opposing targets, which aligns with our observations. Therefore, we believe that the individual target reach was not the result of flexible MU control. Based on our findings, we, therefore, conclude that flexible MU control in the human tibialis anterior muscle was not achieved within a single day of neurofeedback training in our experiments.

**Figure 8.**
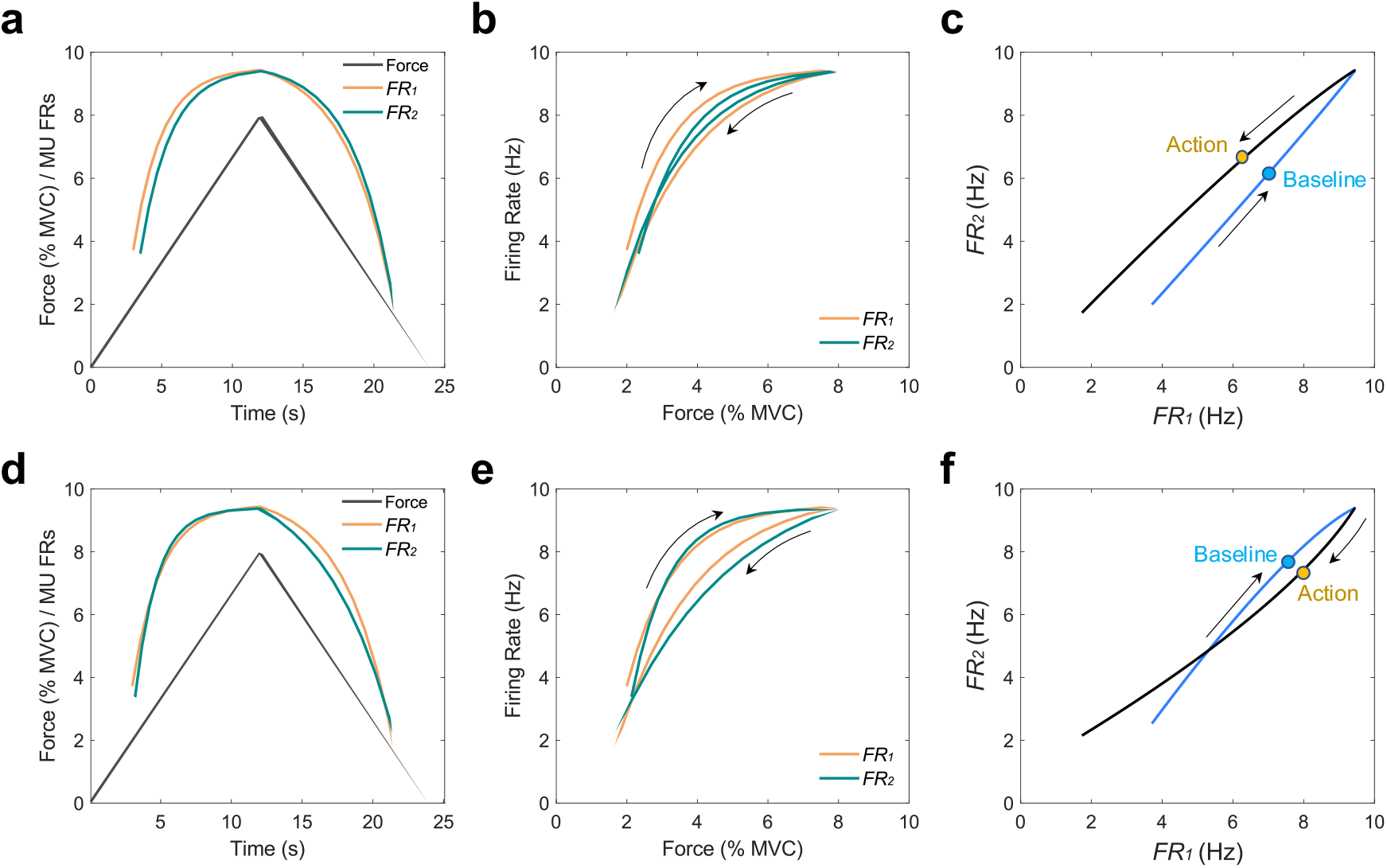
Illustration of how single-quadrant target reach could be achieved due to hysteresis from persistent inward currents. (a-c) Development of time vs. force and MU firing rates (a), force vs. MU firing rates (b), and the firing rates in the state space (c) from a pair of MUs. In this scenario, if participants increase and then decrease force, it would be possible to reach targets in the upper-left quadrant, such as T2 and T4, in the displacement control task, but not T1 or T3. Blue line: force ramp up. Black line: force ramp down. (d-f) Similar to (a-c). With MUs hysteresis properties as illustrated, participants would be able to reach targets in the lower-right quadrant, such as T1 and T3, in the displacement control task, but not T2 or T4. The arrows indicate the direction of the firing rate changes over time.

Some studies have demonstrated the selective activation of MUs from the human tibialis anterior [28, 30] and the biceps brachii [32] muscles. However, showing single MU activation without showing the reversal of the recruitment order cannot be considered as an indication of flexible control as it may be due to the derecruitment of the other MUs [26]. Additionally, results derived from multicompartment muscles should be interpreted with caution as MUs from different muscle compartments which can receive independent input might be involved [36]. While reversing MU recruitment order has been reported during fast force contractions in the human tibialis anterior muscle [29], the findings have not been consistent across different muscles [13, 14, 37, 38]. A recent study by Marshall et al. [25] demonstrated that the activities of MUs can deviate from a one-dimensional manifold during force modulation and also as a result of electrical stimulation of neighboring cortical sites. These findings suggest that direct cortical control over MUs is implemented in the motor system and that flexible MU control is a normal feature of skilled performance. It is therefore plausible to anticipate that participants can readily utilize this flexibility when given direct visual feedback of MU activities. However, our findings suggest that participants did not easily use this flexibility, at least not within a single training session.

The failure to identify flexibility in this study is unlikely attributable to the usage of sEMG recordings rather than intramuscular recordings, as the two methods show a high level of agreement, with over 80% of action potentials identified in common [39-41]. Yet, the selection of the test muscle may be a contributing factor. Like upper limb muscles, motoneurons in the human tibialis anterior muscle receive fast-conducting monosynaptic corticomotoneuronal (CM) connections [42, 43], suggesting potential for direct cortical control. However, the CM connectivity patterns between upper motoneurons and spinal motoneurons may differ between upper and lower limbs. The absence of robust MU firing rate modulation in the displacement zone of our experiments may be due to our focus on activated MUs (excluding any effects from recruitment and derecruitment) and participants were requested to perform tasks where trials lasted over seconds. In contrast, Marshall et al.[25] showed that deviations from rigid control could be observed when comparing MU activity during force modulations at varying speeds including rapid force offsets, such as in sinusoidal movements at different frequencies where a fast force offset was required to track the given force profile. Since our experiment specifically instructed participants to keep the controlled MUs active, fast force offsets were discouraged. However, if flexible control is indeed a normal feature of motor control, participants should be able to naturally utilize the flexibility to reach the targets in displacement zones without the need for excessive training and regardless of the required force modulation. It remains unclear why the flexibility observed in macaque upper limb muscles does not extend to the modulation of MU firing rates in the human tibialis anterior muscle. Could the distinct activity patterns observed across different force profiles be governed by motor programs associated with each force profile, potentially representing the most optimal strategy for motor unit control? If this is the case, the observations reported in Marshall et al. [25] may reflect learned responses tailored to specific motor tasks, rather than presence of readily available flexible MU control, where firing rates can be adjusted quickly and freely. Further research is required to address these unresolved questions.

When executing movement tasks, muscle groups involved in the same task work together as a synergy to increase the efficiency [44]. A similar concept may apply to motor neurons, where motor neuron synergies refer to functional groups of motor units from one or more muscles [18, 19, 45, 46]. The observed changes in recruitment order across different tasks involving the same muscle suggest that flexible MU activation is associated with the engagement of distinct synergy groups [45]. In this study, we found no evidence that participants could independently modulate the firing rates of MUs within the same muscle, despite potential behavioral changes, such as small ankle rotations. This may be attributed to the fact that the majority of MUs in the tibialis anterior muscle likely belong to the same MU synergy group [20, 46]. In the study by Rossato et al. [27], which employed a similar but distinct experimental paradigm to ours—using a less stringent condition without the constraints on derecruitment and with a target area extending beyond the displacement zones— participants were also unable to volitionally dissociate activities of a MU pair from either the same muscle or synergistic muscles during online feedback control. Future research may investigate if flexible control over MUs from a single muscle, but belonging to distinct motor unit synergies, can be achieved through neurofeedback.

### Limitations

The selection of MUs might have influenced the observed properties of the MUs: Firstly, one criterion during MUs selection was to select MUs with similar recruitment thresholds (Supplementary Fig. S4) to ensure that the firing rate of the lower recruitment threshold unit did not reach saturation when the other one was recruited. However, MUs with similar recruitment thresholds are more synchronized when firing at similar frequencies [47], implying that these MUs may receive a larger proportion of common input compared to independent input [23]. Secondly, the controlled MUs exhibited a limited range of firing rate variation of less than 2 Hz (Fig. 5), which may have been the result of MU selection. We selected MUs with low recruitment thresholds in particular so that the training time could be extended and the risk of fatigue was minimized. This may have led to reduced operational range of MU firing rates, as low recruitment threshold units (<10% MVC) typically exhibit an initial acceleration in firing rates upon recruitment, followed by a narrow rate modulation before reaching saturation [48, 49]. This non-linear acceleration of firing rates is attributed to the amplification of synaptic input through the activation of PICs at or below the recruitment threshold, a phenomenon more pronounced in MUs with lower recruitment thresholds [50, 51]. PICs not only limit the efficacy of additional input once they are activated [35, 52], but also enable motoneurons to generate sustained repetitive firing even after the cessation of the synaptic input [53]. These effects can cause the firing rates of MUs with low recruitment thresholds to be less reflective of the actual synaptic input. On the other hand, MUs with higher recruitment thresholds have a more linear force-firing rate relationship [54]. Using MUs with higher recruitment thresholds may offer a larger range for firing rate modulation and reduced synchronization, but it comes with the trade-off of increased likelihood of fatigue, potentially limiting the amount of available training time within a session.

A study in non-human primates suggested multiple cortical drives enabling flexible control of MUs [25]. However, it does not clarify whether these connections can be selectively and voluntarily controlled. In our study, we did not observe volitional control of MU activities that violates the rigid control strategy within single-session training in both tasks. The findings from the difference control task further suggest that violation of a rigid control strategy is not easy to achieve; otherwise, it would likely have been the preferred approach employed by the participants. Even though the displacement control task could not be accomplished within a single-session training, it is still possible that such a control skill could be acquired over an extended period of practice. Studies have shown that one day of training was insufficient for learning a novel activity pattern that falls outside the original manifold of the motor cortex [55], but the learning could occur over multiple days [56]. Similarly, adapting to an entirely new muscle synergy following a remapping of muscle activation patterns did not occur within a single day [57], but was achievable with multi-day training [58]. While it remains unclear whether long-term skill acquisition, such as in musicians, leads to greater flexibility in individual MUs, these individuals have been shown to exhibit greater motorcortical excitability and plasticity compared to untrained individuals [59, 60]. Technical challenges remain in conducting such an experiment over multiple days due to the difficulty of consistently tracking the same MUs across days [61], and learning might not transfer to another day when the selected MUs are varying. Improvements in cross-days MU tracking methodologies could enable future experiments addressing these questions.

## Materials and methods

### Participants

In total 33 able-bodied participants were recruited for this study. Ten participants carried out the displacement control task (1 female and 9 males, aged 23 to 33), and 21 participants carried out the difference control task (11 females and 10 males, aged between 21 and 39 years). Two more participants were recruited but did not perform any task because the MU decomposition quality was initially insufficient to continue the experiment. None of the participants took part in both tasks. Four participants in the difference control task (among them two authors of this study) were exposed to similar but not identical experimental paradigms before their participation. The remaining participants had no such prior experience. The study was approved by the institutional ethics committees of the University of Freiburg (approval number 21-1388-1), and all participants gave their informed consent before starting the experiment.

One participant’s data in the difference control task was not recorded during the baseline phase due to an oversight in the data collection protocol. For this reason, this participant’s data was excluded from all neural analyses but included in all behavioral analyses. In addition, only participants with sufficient MU decomposition quality throughout the entire recording session were included in the analysis in the main results (details are provided in *Decomposition stability and exclusion of participants* section below). Thus, in total nine participants from the displacement control task (1 female and 8 males), and ten participants from the difference control task (5 females and 5 males; 2 of them being an author of this study) were included in the main analyses.

### Experimental setup and data acquisition

Figure 1 illustrates the experimental setup. HD-sEMG signals were acquired from the tibialis anterior muscle of the self-reported dominant leg over the muscle belly aligned to the fiber direction using a 64 channels surface electrode grid (1 mm electrode diameter, 8 mm inter-electrode distance; GR08MM1305, OT Bioelettronica, Torino, Italy) with a 64-channel adapter (AD64F, OT Bioelettronica). Before applying the electrode grid, the area of the skin was shaved. The surface of the skin was then cleaned with an abrasive gel (everi, Spes Medical Srl, Genova, Italy) and subsequently with water to remove any residual gel. A self-adhesive foam layer (FOA08MM1305, OT Bioelettronica) was placed onto the electrode grid, and conductive paste (AC Cream, Spes Medical Srl) was applied between the skin and the electrode grid. After placing the electrode grid, medical adhesive tape was used to ensure the fixation of the grid on the leg. In addition, bipolar sEMG from the fibularis longus (FL), and the lateral and medial heads of the gastrocnemius muscle (GL and GM) were recorded using disposable adhesive surface electrodes (24 mm, CDES000024, Spes Medical Srl). All sEMG signals were filtered with a 10-500 Hz, 4th-order Butterworth bandpass filter, and sampled at 2048 Hz using a Quattrocento Amplifier system (OT Bioelettronica).

The force exerted by the dorsiflexion movement on the dominant leg was measured using an ankle dynamometer (NEG1, OT Bioelettronica) with a positioning system and a single-degree-of-freedom force transducer. The participant placed the dominant leg onto the dynamometer at a 90° angle of the ankle, and the participant’s foot was fixed to the dynamometer with a hook-and-loop tape. The recorded force signal was amplified and low-pass filtered at 33 Hz with a pre-amplifier (Forza, OT Bioelettronica), and then recorded using the Quattrocento amplifier. The communication between the amplifier and the computer was implemented with buffers of data packages consisting of 128 samples, corresponding to 62.5 ms.

### Experimental paradigm

In the beginning of the experiment, the participants’ dorsiflexion force was measured during maximum voluntary contraction (MVC). Afterwards, the participants completed a few trapezoidal force profiles to familiarize themselves with the setup and feedback. This was followed by a single trapezoidal profile to calibrate the MU decomposition. Subsequently, participants performed force tasks and proceeded to the MU control task (displacement or difference control). After the control task they repeated the force and MVC tasks. Further details are provided in the following sections.

#### Pre-experimental calibration

To measure the MVC force level, participants were first asked to perform a maximum isometric dorsiflexion of the ankle three times for 5 s each time, interrupted by a 15 s break. The maximum measured force from the dynamometer was then used to determine the MVC force level and served as a reference force for the rest of the experiment. During the following tasks in the pre-experimental calibration, the instantaneous force produced by isometric dorsiflexion was presented to participants on a computer screen in front of them, together with a force profile that they were supposed to follow. At first, participants were asked to follow a trapezoidal profile (trapezoid I: 10 s ramp-up from 0% to 10% MVC, 20 s hold at 10 % MVC, 10 s ramp-down from 10% to 0% MVC) for 2-5 repetitions as a familiarization to this type of force tracking task. Participants then tracked a second trapezoidal force profile with a longer hold period once (trapezoid II: 10 s ramp-up, 30 s hold at 10 % MVC, 10 s ramp-down). A motor unit decomposition algorithm [62] was calibrated with the HD-sEMG signals recorded during the trapezoid II profile. In the later stages of the experiment (see below), this decomposition algorithm was applied to obtain MU spiking activities in real-time [4, 26] from the HD-sEMG signals. At the end of the calibration sessions, the participants were asked to track the trapezoid I profile one more time. The force-to-firing rate relationships, as well as the spike-triggered averages of the HD-sEMG signals, were obtained from these data and used for MU selection (see below).

#### Force tracking tasks

After the pre-experimental calibration, participants performed three repetitions of the trapezoid I force tracking task, with 10 s rest between repetitions. This set of three repetitions was performed again after the control tasks (see below) to assess changes in MU properties, such as recruitment and derecruitment thresholds and to assess decomposition quality. Recruitment thresholds were defined as the average force values within a 200 ms window centered on the first instance of MU firing rates exceeding 5 Hz during ramp up, while the derecruitment thresholds were defined using the average force values within a 200 ms window but centered at the instance of when the firing rate dropped to 0 Hz during ramp-down [1]. Recruitment and derecruitment thresholds were determined for each trapezoid separately and then averaged across the three repetitions.

Those participants who would later carry out the displacement control task additionally tracked three chirp force profiles after the trapezoid I force tracking task. The amplitude of these force profiles was 2% MVC, and they were centered at 3%, 4%, and 5% MVC. For all chirp profiles the instantaneous frequency increased linearly from 0.1 Hz to 1.3 Hz during the first 30 seconds, and during the subsequent 30 seconds, it decreased linearly from 1.3 Hz to 0.1 Hz. Visual feedback of the applied force was provided throughout the task. The chirp profiles were recorded with the intention to investigate whether MU activities generated during the displacement task that are indicative of flexible control, could also be generated during dynamic force modulation. Since our results did not suggest flexible MU control during the displacement task, the chirp recordings were not further analyzed here.

#### MU selection

The MUs used for control in the displacement and difference control tasks were selected as follows. Participants controlled the vertical position of a cursor on a screen in front of them by the sum of the firing rates of two MUs. The horizontal position of the cursor was fixed. One or two circular areas were shown on the screen (for the difference and displacement control tasks, respectively), reflecting the baseline areas later used in the respective tasks (see below). Participants were asked to repeatedly move the cursor into the indicated circle, during which the experimenter selected two MUs from the available MU pool. The MU selection was based on the following criteria:

1. The recruitment thresholds of the two selected MUs should be as low as possible, preferably below 5 % MVC. MUs with lower recruitment thresholds were preferentially selected to minimize the risk of fatigue during the experiment.
2. The two selected MUs should have similar recruitment thresholds (to avoid the scenario in which one MU has reached its peak firing rate while the other was just being recruited).
3. MUs with more regular spiking, i.e., with a smaller coefficient of variation of inter spike intervals (ISIs) during the hold phase of the force tracking task were preferred, as irregular firing can indicate lower MU decomposition quality.
4. If more than two MUs met criteria 1-3 equally well, an MU pair with a similar range of firing rates across various force levels was selected.

After selecting the two control MUs, the MU with the lower recruitment threshold during the single repeat trapezoid I profile done at the end of the calibration sessions was assigned as *MU*_1_ and the one with higher recruitment threshold MU as *MU*_2_. Additional gain factors for each MU were manually set. The gain factors functioned as multipliers to the firing rates to ensure similar adjusted firing ranges of the two MUs. This was crucial for the displacement control task because the baseline area was centered along the diagonal of the firing rate state space. In the difference control experiment, ensuring similar firing rates after gain adjustment was important to minimize cursor movement along the horizontal axis once the cursor movement in the horizontal direction was unlocked at the beginning of the action phase. For both MUs, gain-adjusted firing rates between 7-9 Hz were targeted when reaching the baseline area in the difference control task. The gain factors ratio *MU*_1_/*MU*_2_ was 0.98 ± 0.03 (mean ± s.d) for the displacement control task, and 0.97 ± 0.09 (mean ± s.d) for the difference control task.

#### Displacement control task

In this task, participants controlled a cursor on the screen in front of them with the gain-adjusted firing rates from two MUs, *MU*_1_ and *MU*_2_. The firing rate of *MU*_1_ controlled the horizontal cursor position (along the x-axis), while the firing rate of *MU*_2_ controlled the vertical cursor position (along the y-axis). The cursor position was updated at the end of each data package.

The single-session experiment comprised 5 blocks and a total of 200 trials. Each block included 40 trials, with 10 trials per target per block, and the targets were presented in a random order. Each trial consisted of two phases: the baseline phase and the action phase (see Fig. 2c). To initiate a trial, participants had to move the cursor over the baseline area and hold it in the baseline area for 3 seconds. The holding time was indicated by a circular timer surrounding the baseline area. During the 3-second hold phase, the cursor was required to remain within the baseline area. If it left the baseline for more than one consecutive second, the timer was reset to zero. The baseline areas (grey) were represented by circles of 2 Hz radii centered at 6 Hz/6 Hz (A) and 8 Hz/8 Hz (B) (Fig. 2b). When the baseline phase ended, the circle showing the baseline area disappeared, and the trial entered the action phase.

Baseline firing rates were determined by averaging firing rates across the 3-second holding phase, and these baseline firing rates were used to set the position of the subsequent target. Targets appeared as a square box with a side length of 5 Hz with their upper left vertex (for T1 and T3) or their lower right vertex (for T2 and T4) at the average baseline firing rate (Fig. 2b). As a consequence, the targets visualized the displacement areas with regard to the baseline firing rates, as illustrated in Fig. 2a. In each trial, either T1 or T2 followed baseline area A, while either T3 or T4 was presented after baseline area B. The participants were given 10 seconds to move and hold the cursor in the target area for as long as possible. Participants received a reward of 10 points and a coin-earning sound each time the cursor remained within the target for 125 ms. The total number of points that participants had obtained was displayed at the top of the screen. In addition, two red bars were positioned along the x- and y-axes. The cursor touched the bars if one MU was derecruited, and participants received an immediate penalty of minus 50 points upon contact with the bars. Subsequent penalties were applied only if the cursor remained over the bar for an additional 600 ms or longer. If at any time during the action phase both MUs were derecruited simultaneously, the current trial was discontinued. A 5-second break occurred between the individual trials within a block and a 2-minute break between the individual blocks.

#### Difference control task

In this task, the vertical cursor position was controlled by the sum of the firing rates of *MU*_1_ and *MU*_2_, whereas the horizontal cursor velocity was controlled by the difference of the firing rates (Fig. 7a). The cursor position was updated at the end of each data package. Firing rate differences needed to exceed a threshold to generate a non-zero velocity. The x-position of the cursor (*x*_*t*_) was computed as

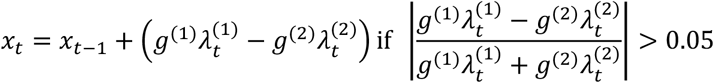

where *t* is the time step, *x*_*t*_ is the x-position of the cursor at time *t, g*^(1)^, *g*^(2)^ are the gain factors for *MU*_1_ and *MU*_2_ and 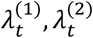 are the firing rates of *MU*_1_ and *MU*_2_ at time *t*. If the absolute difference of the gain-adjusted firing rates was not above the threshold (0.05), the velocity was set to 0 to reduce noisy cursor movements. The cursor position along the y-axis was set to the sum of the gain-adjusted firing rates, i.e.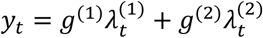 where *y*_*t*_ is the y-position of the cursor at time *t*.

The single-session experiment comprised 5 blocks and a total of 180 trials. Each block included 36 trials, with 18 trials per target presented in a random order. Participants engaged in a food delivery game involving two sequential steps in each trial (Supplementary Fig. 1a): 1) Baseline phase: Participants picked up the delivery by holding the cursor (in hand shape) over the pick-up area for one consecutive second. The cursor could only move vertically during the baseline phase. A circular timer surrounding the pick-up area indicated the passage of time. 2) Action phase: Once the baseline phase was successfully completed (the delivery was picked up), the participants proceeded by delivering the designated item (cursor) to one of the two houses (targets). The target house was denoted by a thought bubble appearing in the top-right corner of the house (see Supplementary Fig. 1a). During the action phase, the participants were asked to keep both MUs active throughout the trial. The states of the two MUs were indicated by two traffic lights on the screen, with green and grey indicating an active and inactive state, respectively. The cursor would only move when both MUs were active to discourage MU derecruitment. The cursor would stop moving if one of the MUs was derecruited and the trial would be discontinued when both MUs were derecruited. When the cursor entered the target, a circular timer surrounding the target house appeared to signify successful target reach and indicate the duration the cursor was held within the target. In each trial, the action phase was limited to a maximum duration of 30 s, during which participants needed to hold the cursor within the target for three consecutive seconds for a trial to be considered successful. A trial was discontinued once the target was reached successfully. Upon success, the participants were awarded a score of 100 points, accompanied by a coin-earning sound. If one MU was derecruited during the 3-second holding period over the target, the timer around the target would be reset to zero so that a trial could only be successful without MU derecruitment during the hold period. Between trials, there was a 5-second pause followed by a 3-second countdown to allow participants to prepare for the next trial. In addition, there were 2-minute breaks between the individual blocks.

Note that we present the results from displacement control task first in the result section while the experiments involving the displacement control task were conducted chronologically after the experiments involving the difference control task.

### Decomposition stability and exclusion of participants

The HD-sEMG decomposition algorithm used in this study has previously been validated and shown to yield accurate decomposition when compared to intramuscular recordings [62]. We performed a systematic offline analysis of the decomposition stability in our experiments and excluded participants from further analysis when at least one of the two control MUs exhibited insufficient decomposition stability. The following measures were used to assess decomposition stability of a motor unit:

#### (1) Pulse trains

The decomposition algorithm iteratively computes motor unit specific separation vectors onto which the HD-sEMG signals are projected to obtain the motor unit pulse trains. The discharge times of a motor unit are then acquired by k-means clustering (with k=2) of the peaks of the squared motor unit pulse train (pp^2^) [62]. Separation vectors and cluster parameters were obtained from the recordings during a trapezoid force profile in the pre-experimental calibration phase (referred to here as the decomposition ramp). A silhouette measure (SIL) was computed to evaluate the consistency of the clustering and only motor units with SIL>0.9 for the data from the decomposition ramp were used [62]. The separation vectors and cluster parameters obtained from the decomposition ramp were used in the control blocks to obtain MU spike trains from HD-sEMG recordings online.

Non-stationarities during subsequent recordings can lead to degrading cluster separability over time and thus, wrong cluster assignment of new incoming data during online experiments. As a consequence, actual action potentials may be missed (false negatives) or be detected where there are none (false positives). To assess the stability of cluster separability, we re-applied the k-means clustering algorithm offline after the experiments, to the recordings during the control blocks. Here, k-means was applied separately to the data from each block to obtain individual cluster thresholds that were optimized for each block. See supplementary figures S8a and S9a for units from two participants with low and high stability, respectively. Occasional extreme pp^2^ values were removed before performing k-means offline, using a threshold of five times the median of the pp^2^ values from action potentials classified during the decomposition ramp. For participants included in the main results, the MUs used for control had SIL values obtained from offline clustering exceeding 0.89 across all control blocks.

Even if cluster separability remains high, non-stationarities in the HD-sEMG signals can result in reduced pp^2^ peak values and thus, shifts of the action potential cluster to lower values (Supplementary Fig. S8a in contrast to Supplementary Fig. S9a). In such situations, a static cluster threshold as used in the online experiments can lead to missed spikes. We compared the spike trains obtained after re-application of k-means to the spike trains obtained online and computed the fraction of missed spikes under the assumption that the spike trains obtained after re-application of k-means reflect the ground truth (Supplementary Fig. S8c and S9c, blue lines). This estimate may overstate the number of missed spikes due to false positives in the clustering after re-application, for example resulting from decreased cluster separability. For the participant shown in Supplementary Fig. S8a this is reflected in the less separable clusters in later blocks. To improve the estimate of missed spikes, we introduced the relative inter-spike interval (*rISI*), which measures the duration of an ISI in relation to the median duration of adjacent ISIs:

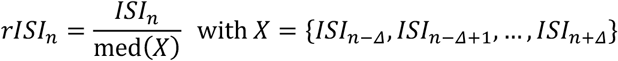

Here, *n* indexes the sequence of ISIs across the recording and Δ specifies the number of adjacent ISI before and after the current ISI that are used to calculate the median ISI. Δ was set to 12 in this analysis. For slowly changing firing rates and given the regularity of MU spiking, *rISI* values should fluctuate around 1 while missed spikes would yield peaks around 2 and 3 in the *rISI* histograms (Supplementary Fig. S8b, S9b). We therefore estimated the fraction of missed spikes by using only those additional spikes after re-application of k-means that were found in ISIs with *rISI* values above 1.2 or 1.5 (Supplementary Fig. S8c, S9c). This led to a slightly smaller fraction of estimated missed spikes in blocks where clusters were less separable (Supplementary Fig. S8c). If the fraction of the estimated missing spikes was too high or showed a strong increase across blocks, the participant was excluded from the analysis.

#### (2) ISI histograms

We assessed stability also by visual inspection of the ISI histograms during the control blocks. The emergence of distinct second or third peaks in the ISI histograms (see Supplementary Fig. S8d for an example and in contrast Supplementary Fig. S9d), located at two or three times the ISI of the highest peak, may also indicate the presence of missing spikes. When such distinct peaks were observed we typically also observed a higher fraction of missed spikes according to the measure described in the previous paragraph and therefore, the participant was excluded.

Besides false negatives, false positive spike detections could also lead to incorrect estimation of firing rates. False positives could result from a degradation in decomposition quality, additional noise in the recorded EMG signal (e.g., external interference), or the conflation of signals from newly recruited MUs with those of the decomposed MUs, compromising the accuracy of the spike detection. False positives would produce an excess of short ISI that would not be expected for the recorded MU during the used force levels and tasks. If an excess of such short ISIs was observed in the ISI histograms for the control blocks, the participant was also excluded.

We also investigated the ISI histograms from the trapezoid force profiles performed before and after the control tasks (pre and post ramps). The ISI histograms of two example participants with excess short ISIs during the pre/post ramps are shown in Supplementary Fig. S8e, f. In the example shown in the upper row of Supplementary Fig. S8e, an excess of short ISIs was observed during the hold phase of the first trapezoid (ramp1) of the force tracking task before the control task (ramp hold pre) and during the first and third trapezoids (ramp1 and ramp3) after the control blocks (ramp hold post). However, the ISI histograms from the control blocks of the same participant (lower row of Supplementary Fig. S8e) show no excess of short ISIs, except for a small peak during block 1 (black arrow, note that this peak is not relevant for the control feedback as ISIs <20ms were excluded from firing rate estimation). While all ramps were carried out at 10% MVC, most participants produced forces below 5% MVC during control blocks (see Fig. 3b and Supplementary Fig. S1d). This suggests that the short ISI during the pre/post ramps were a result of the higher forces. Therefore, we computed ISI histograms from the ramps using only ISIs occurring during force levels up to the 99th and 90th percentiles of the force levels observed during the control blocks (Supplementary Fig. S8e). The observed decrease in the number of short ISIs with decreasing force thresholds corroborates the previous interpretation. In such cases and when in addition no excess of short ISIs were observed during the control blocks, we, therefore, concluded that the decomposition stability was not affected during the control blocks and did not exclude the respective participant. Conversely, if short ISIs persisted after applying lower force thresholds and/or an excess of short ISIs were observed during the control blocks (as illustrated for the unit from another participant shown in Supplementary Fig. S8f), the degradation in decomposition quality or interference from external noise was likely relevant for the control blocks, and the participant was, thus, excluded from the main results.

Note that the exclusion of participants did not affect our main conclusions as Bayesian hypotheses tests using all participants also indicate that there is no evidence supporting target reach (Supplementary Fig. S5 and Supplementary Table 4, 5).

Supplementary figures S8 and S9 show the results from the analyses to assess decomposition stability of individual units from selected example participants. The analyses for both control MUs of all participants are provided in the Supplementary figures S10-S40.

### Analysis

#### Firing rate estimation

Two methods were used to estimate firing rates from MU spike trains: (i) Convolution of the MU spike trains with a 400 ms Hanning window. This method was used when determining the recruitment and de-recruitment thresholds. (ii) Estimation of firing rates from interspike intervals (ISI). This method was used to compute the MU firing rates online during the experiments with MU controlled cursor movement. For (ii), the real-time firing rate estimate *λ* was updated at each new spike according to the following equation:

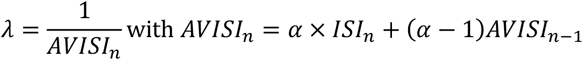

where *ISI*_*n*_ is the ISI between the current and the previous spike, *AVISIn*_−1_is the weighted average of previous ISIs updated at each new spike and *n* the spike index. The current firing rate estimate *λ* was computed as the inverse of the last updated average ISI. The parameter *α* was set to 0.2. If no spike was detected for over 500 ms, the firing rate was set to 0 Hz. The maximum value for *AVISI*_*n*_ was capped at 1000 ms to prevent a slow increase of the firing rate estimate after a longer period of inactivity (5 seconds or longer). A slow firing rate increase could lead to excessive force production by the participants and thus increase the likelihood of muscle fatigue.

#### Removal of data points and trials

This study focused on the continuous modulation of motor unit activities and investigated the flexibility of MU firing rate modulation while MUs were active. For this purpose, we excluded all firing rate data from the analysis within a 250 ms window centered around any period where any of the two MUs had a firing rate of 0 Hz. This approach reduced the influence of derecruitment and recruitment in the analysis. When analyzing the firing rates in the action phases, only trials that lasted longer than 3 seconds after the removal of the time windows were used. Out of 1800 trials from 9 participants in the displacement control experiment, a total of 1740 trials met this criterion and were used for the analysis. In the difference control experiment, 1589 out of 1800 trials from 10 participants met this criterion. To assess robustness, a separate analysis of MU firing rates was conducted excluding all trials with derecruitment in one or both control MUs and using only the second half of the action phase (Supplementary Fig. S7 and Table 8,9 and 10).

#### Determination of recruitment order for MU_1_ and MU_2_

For the analyses of the difference control task, we assigned *MU*_1_ and *MU*_2_ in such a way that *MU*_1_ had a lower recruitment threshold than *MU*_2_. The recruitment order of the two units was obtained from the tracking of a trapezoidal force profile (as described above), and separately from the control tasks. For the control tasks, we used the instantaneous firing rate estimates from each individual trial and the average of the measured forces at the first occurrence of firing rates above 5 Hz, if present, as the recruitment thresholds. If the recruitment order for the control and the force tracking tasks was different, a Bayesian Wilcoxon signed-rank test was conducted to compare the recruitment thresholds obtained from the trials of the control task to the recruitment threshold obtained from the trapezoidal force profile. If the resulting Bayes Factor was larger than 10 (indicating strong evidence for a difference), the recruitment thresholds from the control task were used to determine the recruitment order of the two MUs. Otherwise, the recruitment thresholds from the trapezoidal force profile were used. Since the trapezoidal force profile is the conventional method for determining recruitment thresholds and given the higher variability of the force increase at the beginning of the control trials (the time course of the force increase was not queued but freely chosen by the participants), we only used the recruitment thresholds from the control task if there was strong evidence for a difference to the trapezoidal force task.

For the displacement control task, the recruitment order obtained from the trapezoidal force profile before the control blocks was maintained in the analysis of the main results as it reflected the visual feedback during the online experiments. A robustness analysis was performed using recruitment thresholds derived from both the trapezoidal force profile and individual control trials as described above for the difference control task. Here, the MUs were reassigned, such that the recruitment threshold of *MU*_1_ was lower than that of *MU*_2_. The results after reassignment were consistent with those obtained when the recruitment order was not changed (Supplementary Fig. S6 and Table 6 and 7).

### Statistics

We used Bayesian methods for hypothesis testing. The Bayes Factor BF_10_ measures the level of support for comparing two hypotheses (H_1_ and H_0_) on a range from 0 to +∞ based on data. A BF_10_ value of 1 indicates that the data do not provide evidence in favor of either H_0_ or H_1_ while a value below or above 1 provides evidence in favor of H_0_ or H_1_, respectively. To interpret Bayes Factors, Lee & Wagenmakers (2014) [63] suggested a heuristic classification where values of BF_10_ between 1 and 3 would be considered anecdotal, values between 3 and 10 moderate, values between 10 and 30 strong, values between 30 and 100 very strong and values above 100 extreme evidence for H1. We applied this heuristic with a modification: any Bayes Factor value greater than 10 was consistently interpreted as strong evidence, without further differentiation into very strong or extreme evidence. Correspondingly, for values of BF_10_ below 1, the range between 1 and 1/3 was considered anecdotal, the range between 1/3 and 1/10 moderate and values below 1/10 strong evidence for H0. Note, that BF_01_ is given by 1/ BF_10_.

#### One sample and paired samples designs

For one sample and paired sample comparisons, we used the Bayesian Wilcoxon signed-rank test [64] implemented in JASP (https://jasp-stats.org). The default Cauchy prior with location zero and scale 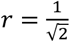 was used. To assess the robustness of the Bayes factor with regard to the prior, the Bayes factor was additionally computed for prior scales 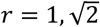 that include also wider priors. As the Bayes factor did not change qualitatively across the prior scales, we reported the Bayes factor obtained with the default prior scale of 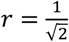 in the main text. The results for the different prior scales can be found in Supplementary Table 11.

#### Repeated measures ANOVA designs

For repeated-measures (RM) ANOVA designs, we used the Bayesian RM ANOVA implemented in JASP with random slopes [65] and the Jeffreys-Zellner-Siow prior [66]. The inclusion Bayes factor (BF_inclusion_) was used to assess the evidence for a predictor to be included [65, 67]. We computed Bayes factors for three different prior scales for fixed effects 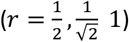 with the prior scale for random effects set to twice that of the fixed effects. As the Bayes Factors were not qualitatively different across the scales, we reported the Bayes Factors for the default scales 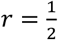 for fixed and *r* = 1 for random effects) in the main text. The complete results for all prior scales are reported in Supplementary Table 11. Post hoc tests were conducted using the default settings for multiple comparisons in JASP which uses a correction for multiple comparisons according to [68].

#### Trend analysis across blocks

We conducted trend analyses across blocks by linear regression of the measurement (e.g. success rate) across blocks. The value for the slope coefficient obtained from the linear least-squares fit was used as a measure of the trend. We performed a Bayesian Wilcoxon signed-rank test on the obtained slopes to quantify evidence for non-zero trends. The obtained Bayes factors BF_10_, with H_0_ reflecting a slope of 0 and H_1_ a slope different from 0, were reported as trend-BF_10_. To compare trends in repeated measurements, a Bayesian repeated-measures ANOVA was applied to assess differences between slopes, with the resulting inclusion Bayes factor reported as trend-BF_inclusion_. We used the same prior scales for the Bayesian Wilcoxon signed-rank test and the Bayesian RM ANOVA for the trends as described in the previous sections. Bayesian Wilcoxon signed-rank tests and Bayesian RM ANOVA were computed using JASP.

#### Analysis of motor unit firing rate changes

##### Single target analysis

We analyzed the firing rate changes between the baseline and action phases for all four targets separately. With *ΔFR*_1_ and *ΔFR*_2_ referring to the changes of the firing rates of *MU*_1_ and *MU*_2_ between baseline and action, the following three hypotheses were considered (see Fig. 9):

**Figure 9.**
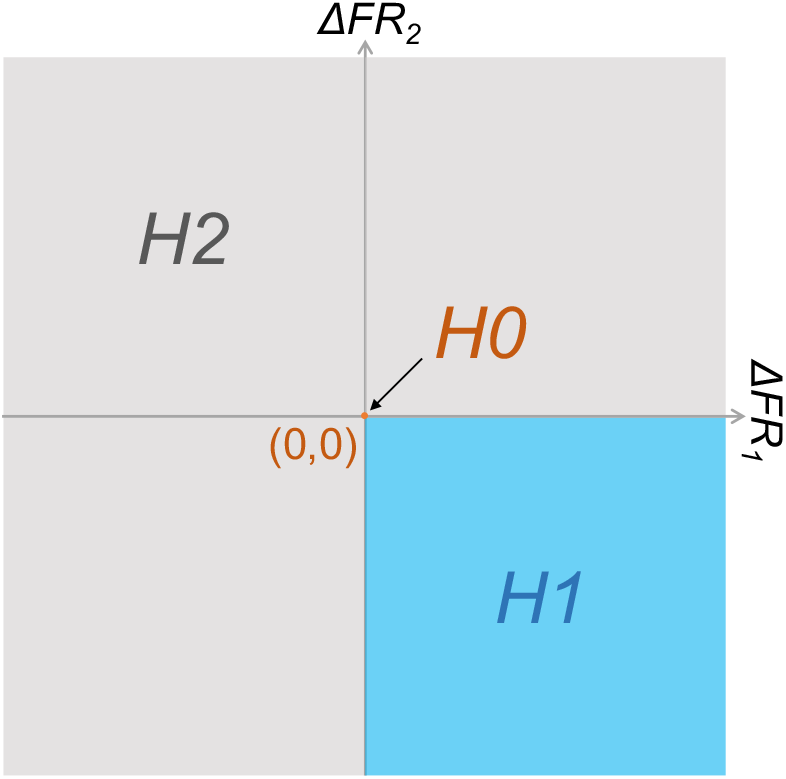
Illustration for three hypotheses used to test MU firing rates changes using the T1/T3 area as an example. Orange color for *H*_0_, light blue color for *H*_1_ and grey color for *H*_2_.

H_0_: no change in firing rates, i.e. *ΔFR*_1_ = 0 ∧ Δ*FR*_2_ = 0

H_1_: successful target reach, i.e. *ΔFR*_1_ > 0 ∧ Δ*FR*_2_ < 0 for T1 and T3, *ΔFR*_1_ < 0 ∧ Δ*FR*_2_ > 0 for T2 and T4.

H_2_: no successful target reach but a change of firing rate in at least one of the two units. This hypothesis reflects the complement of H_0_ and H_1_.

Bayes factors between all pairs of hypotheses were computed using the R package BFpack [69]. BFpack uses default priors that do not require the specification of the scale of the expected effects (for details see [69]). The analysis provides Bayes Factors for all pairwise combinations of the three hypotheses yielding in total six Bayes Factors (BF_xy_ with *x* ∈ {0,1,2} and *y* ∈ {0,1,2} and *x* ≠ *y*). Given that BF_xy_=1/BF_yx_ and that BF_xy_=BF_xz_ · BF_zy_ any two Bayes Factors that include each hypothesis at least once contain the complete information.

The Bayes Factors were interpreted as follows:

1. If the Bayes Factors of one hypothesis over both of the other two hypotheses exceeded 10 (or 3), we concluded that the data provides strong (or moderate) evidence for that hypothesis. In other words, if BF_20_>10 and BF_21_>10 the data provides strong evidence for H_2_. Correspondingly, if BF_01_>10 and BF_02_>10 or if BF_10_>10 and BF_12_>10 the data provides strong evidence for H_0_ or H_1_, respectively. For all three cases, if both BFs were larger than 3 but at least one of them not larger than 10, we concluded that our data provides moderate evidence for H_0_, H_1_ or H_2_ respectively.
2. In cases where no single hypothesis dominates over both of the other two hypotheses, we may still find moderate or strong evidence against H_1_. If none of the conditions of (1) apply but if either BF_21_>10 or BF_01_>10 the data provides strong evidence against H_1_ regardless of the values of the other Bayes Factors in each case. Likewise, if either 3<BF_21_≤10 or 3<BF_01_≤10, we concluded that our data provides moderate evidence against H_1_.
3. If none of the conditions of (1) or (2) apply, the data provides only weak or anecdotal evidence with regard to H_1_.

We report the Bayes Factors BF_20_, BF_21_ and BF_10_ from which all other Bayes Factors can be computed. While two Bayes Factors would be sufficient, we report three Bayes Factors for convenience. For the analyses of the population of participants, trial-averaged firing rates were used assuming statistical independence between participants. For single participant analyses, firing rates from individual trials were used assuming statistical independence between trials.

##### Opposing targets analysis

We jointly analysed the firing rate changes for pairs of opposing targets (T1 and T2 or T3 and T4). With 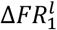 and 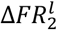 referring to the firing rate changes of *MU*_1_ and *MU*_2_ for a lower target (i.e. either T1 or T3) and 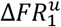 and 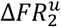 to the firing rate changes for an upper target (i.e. either T2 or T4) we considered the following three hypothesis for each pair of opposing targets:

H_0_: no change in firing rates for both targets, i.e. 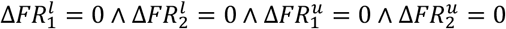

H_1_: successful reach of both targets, i.e. 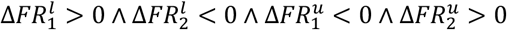

H_2_: no successful reach of both targets but a change of firing rate in at least one of the two units during at least one of the two targets. This hypothesis reflects the complement of H_0_ and H_1_.

The rest of this analysis was carried out in the same way as in the single target case.

#### Fitting and visualizing bivariate Gaussian distributions

When fitting bivariate Gaussian distributions to firing rates, we used the sample mean 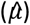 and the sample covariance matrix 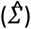. To visualize the bivariate Gaussian distributions, we used ellipses centered at 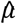 and with axes orientated along the eigenvectors of 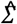. The width and height of the ellipses were chosen so that a random number drawn from the bivariate Gaussian lies within the ellipse with a probability of 0.8. This was achieved by setting the width/height of the ellipse to the square roots of the eigenvalues multiplied by 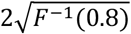(0.8) where *F*^−1^ is the inverse of the 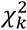 cumulative distribution function with *k* = 2 degrees of freedom [70].

## Acknowledgements

We thank Deren Yusuf Barsakcioglu for assistance with the decomposition algorithm and Tobias Pistohl for his assistance in resolving various technical challenges. This research was supported in part by the European Commission grant H2020 NIMA (FETOPEN 899626), European Research Council (ERC) under the Synergy Grant Natural BionicS (810346), the EPSRC Transformative Healthcare for 2050 project NISNEM Technology (EP/T020970/1), the German Research Foundation (DFG) through grant no INST 39/1014-1, the state of Baden-Württemberg through the Struktur-und Innovationsfonds (SI-BW), ECHOES (ERC Starting) under Grant 101077693 and in part by a Consolidación Investigadora grant (CNS2022-135366) funded by MCIN/AEI/10.13039/ 501100011033 and UE’s NextGenerationEU/PRTR funds. We acknowledge support by the Open Access Publication Fund of the University of Freiburg.

## Competing interest disclosure

Dario Farina is inventor of two patents (Neural Interface. UK Patent application no. GB1813762.0. August 23, 2018 and Neural interface. UK Patent application no. GB2014671.8. September 17, 2020) related to the methods and applications of this work. Dario Farina is also Scientific Advisor for neural interfacing for Meta, Reality Labs, and for high-density EMG technology for OT Bioelettronica, Italy.

