## Supplementary material for "Rigid Control of Motor Unit Firing Rates in the Human Tibialis Anterior Muscle Persists during Neurofeedback"

| | $BF_{20}$ | $BF_{21}$ | $BF_{10}$ |
|---|---|---|---|
| ***T1*** | | | |
| **B1** | 18 | 16 | 1 |
| **B2** | 2.8 | 10.9 | 0.26 |
| **B3** | 58 | 18 | 2.6 |
| **B4** | 12 | 44 | 0.27 |
| **B5** | 97 | 190 | 0.5 |
| ***T2*** | | | |
| **B1** | $1.2 \times 10^4$ | $2.3 \times 10^3$ | 5.4 |
| **B2** | $5.4 \times 10^3$ | 71 | 75 |
| **B3** | $1.4 \times 10^7$ | 93 | $1.5 \times 10^5$ |
| **B4** | $5 \times 10^3$ | 24 | 210 |
| **B5** | $1.3 \times 10^6$ | 78 | $1.7 \times 10^4$ |
| ***T3*** | | | |
| **B1** | 154 | 83 | 1.85 |
| **B2** | 1.5 | 9.8 | 0.16 |
| **B3** | 1.7 | 17 | 0.09 |
| **B4** | 11 | 6 | 1.86 |
| **B5** | 2.2 | 10 | 0.22 |
| ***T4*** | | | |
| **B1** | 57 | 22 | 2.6 |
| **B2** | 54 | 2.99 | 18 |
| **B3** | **97** | **0.4** | **230** |
| **B4** | 10.4 | 2.1 | 4.9 |
| **B5** | 31 | 1.9 | 16 |

Supplementary Table 1. Bayes factors for the displacement experiment for individual targets and blocks.

| P22 | $BF_{20}$ | $BF_{21}$ | $BF_{10}$ |
|---|---|---|---|
| T1 | $6.6 \times 10^{17}$ | $2.5 \times 10^{11}$ | $2.6 \times 10^6$ |
| T2 | 2590 | $4 \times 10^4$ | 0.065 |
| T3 | $5.8 \times 10^{12}$ | 336 | $1.7 \times 10^8$ |
| T4 | 199 | $4 \times 10^4$ | 0.005 |

| P23 | $BF_{20}$ | $BF_{21}$ | $BF_{10}$ |
|---|---|---|---|
| T1 | 1991 | $6.4 \times 10^5$ | 0.003 |
| **T2** | **$5.4 \times 10^9$** | **0.18** | **$3 \times 10^{10}$** |
| T3 | 1921 | $7.7 \times 10^5$ | 0.003 |
| **T4** | **$1.2 \times 10^4$** | **$7.2 \times 10^{-5}$** | **$1.7 \times 10^8$** |

| P24 | $BF_{20}$ | $BF_{21}$ | $BF_{10}$ |
|---|---|---|---|
| T1 | $1 \times 10^9$ | $5 \times 10^{11}$ | $2 \times 10^{-3}$ |
| **T2** | **$1 \times 10^9$** | **0.44** | **$2.4 \times 10^9$** |
| T3 | $1.7 \times 10^{14}$ | $1.1 \times 10^{16}$ | 0.015 |
| T4 | 0.021 | 0.26 | 0.083 |

| P25 | $BF_{20}$ | $BF_{21}$ | $BF_{10}$ |
|---|---|---|---|
| T1 | $4.3 \times 10^9$ | $9 \times 10^{10}$ | 0.048 |
| T2 | $5.4 \times 10^{17}$ | 362 | $1.5 \times 10^{15}$ |
| T3 | 0.43 | 90 | 0.005 |
| **T4** | **$5.5 \times 10^9$** | **0.84** | **$6.6 \times 10^9$** |

| P26 | $BF_{20}$ | $BF_{21}$ | $BF_{10}$ |
|---|---|---|---|
| **T1** | **$1.5 \times 10^5$** | **0.04** | **$3.7 \times 10^6$** |
| T2 | $3.5 \times 10^{14}$ | $2.8 \times 10^{12}$ | 124 |
| T3 | $1.5 \times 10^5$ | 3.3 | $4.5 \times 10^4$ |
| T4 | 3592 | 21 | 171 |

| P27 | $BF_{20}$ | $BF_{21}$ | $BF_{10}$ |
|---|---|---|---|
| T1 | 798 | 42.8 | 18.6 |
| T2 | $5.3 \times 10^{12}$ | $3.9 \times 10^9$ | $1.4 \times 10^3$ |
| T3 | $3 \times 10^8$ | $9.9 \times 10^9$ | 0.031 |
| T4 | 11.3 | 255 | 0.044 |

| P28 | $BF_{20}$ | $BF_{21}$ | $BF_{10}$ |
|---|---|---|---|
| **T1** | **43.7** | **0.15** | **300** |
| T2 | $7.2 \times 10^{16}$ | $1.3 \times 10^{15}$ | 56.8 |
| T3 | $3.9 \times 10^4$ | 3525 | 11 |
| T4 | $9.1 \times 10^7$ | 9304 | 9800 |

| P29 | $BF_{20}$ | $BF_{21}$ | $BF_{10}$ |
|---|---|---|---|
| T1 | 641 | $5.5 \times 10^4$ | 0.012 |
| T2 | $1.2 \times 10^{15}$ | $1.4 \times 10^9$ | $8.6 \times 10^5$ |
| T3 | 0.93 | 97 | 0.01 |
| T4 | $6 \times 10^6$ | 3566 | 168 |

| P30 | $BF_{20}$ | $BF_{21}$ | $BF_{10}$ |
|---|---|---|---|
| T1 | $2.9 \times 10^{13}$ | $7.9 \times 10^{10}$ | 367 |
| T2 | $5.2 \times 10^6$ | $5.2 \times 10^4$ | 100 |
| T3 | $1.8 \times 10^7$ | 13.4 | $1.4 \times 10^6$ |
| T4 | 1.15 | 259 | 0.004 |

| P31 | $BF_{20}$ | $BF_{21}$ | $BF_{10}$ |
|---|---|---|---|
| T1 | $6.6 \times 10^7$ | 1.02 | $6.5 \times 10^7$ |
| T2 | $1.2 \times 10^{11}$ | $5.2 \times 10^{10}$ | 2.25 |
| **T3** | **$8.1 \times 10^4$** | **0.004** | **$1.9 \times 10^7$** |
| T4 | $7.6 \times 10^9$ | $1.1 \times 10^{10}$ | 0.67 |

Supplementary Table 2: Bayes factors for individual participants and targets. See methods section for a more detailed explanation of the interpretation of BF. Evidence for target reach include P23(T2 and T4), P24(T2), P25(T4), P26(T1), P28(T1) and P31(T3). Participant P28 was excluded from the main results analysis due to instability in MU decomposition.

#### Real time difference control with motor unit activities: experimental design and performance evaluation

In this experiment, participants were told to play a food delivery game by first moving the cursor in cartoon hand shape over the baseline area (Fig. S1a, white open circle) and hold it there for 1 second (baseline phase) to pick up the delivery (Fig. S1a$_1$). During this phase, the horizontal cursor position was fixed. After the delivery was picked up, the cursor turned from hand to delivery icon, and the trial entered the action phase, during which the cursor was free to move horizontally and vertically. A delivery item was randomly selected in each trial from a pool of six options: cheese, a basket of apples, milk, apple juice, ice cream, and an apple pie. The varied selection of items was used to increase participant engagement. The delivery was only successful if the cursor was held at the target house for 3 consecutive seconds within 30 seconds after the delivery was picked up (Fig. S1a$_2$). Participants were asked to keep both motor units active throughout the trial and keep the cursor around the same vertical position (to not make a drastic change in the exerted force level). Activation of the MUs appeared as two traffic lights on the screen (Fig. S1a; green for active, grey for inactive). The trial ended if both MUs were de-recruited at the same time. In this experiment, rather than altering motor unit firing rates in opposing directions, participants were only required to generate a difference in the firing rates to move the cursor towards the targets (Fig. S1b). Figure S1c shows the activity map of the two control MUs from the same example trial as in Fig. 7c-d in the main text.

For the purpose of the analysis, the two MUs were numbered such that MU1 had a lower recruitment threshold than MU2 (see Methods). There was no evidence for a trend of the measured forces across blocks (Fig. S1d, trend-$BF_{10}$ = 0.5). We employed two metrics to assess performance: success rate and latency. Success rate was computed as the percentage of successful trials within each block. Latency was defined by the time from the beginning of the action phase until successful target reach (cursor hold on target area for 3 consecutive seconds) and thus only computed for successful trials. Our results showed an increase of the success rate from 49% in the first block to 68% in the last block (Fig. 7f in the main text, trend-$BF_{10}$ >100) in parallel to a decrease in latency from 14 s in the first to 11 s in the last block (Fig. S1e, trend-$BF_{10}$ = 20).

There was strong evidence for a change of the success rate and latency across blocks for T2 but not for T1 (Fig. 7f in the main text for success rate, Fig. S1f, trend-$BF_{10}$ =0.4 and 76 for T1 and T2), however, there was only anecdotal evidence for a difference in the trends of the success rates (Fig. 7f right in the main text , Bayesian Wilcoxon signed-rank test for trend-$BF_{10}$ =1.6) and moderate evidence for a difference in the trends of the latencies (Fig. S1f, Bayesian Wilcoxon signed-rank test for trend-$BF_{10}$ = 8) between the targets. The delivery item had no impact on the success rate (Bayesian repeated measures ANOVA, $BF_{inclusion}$ = 0.26 for delivery item).

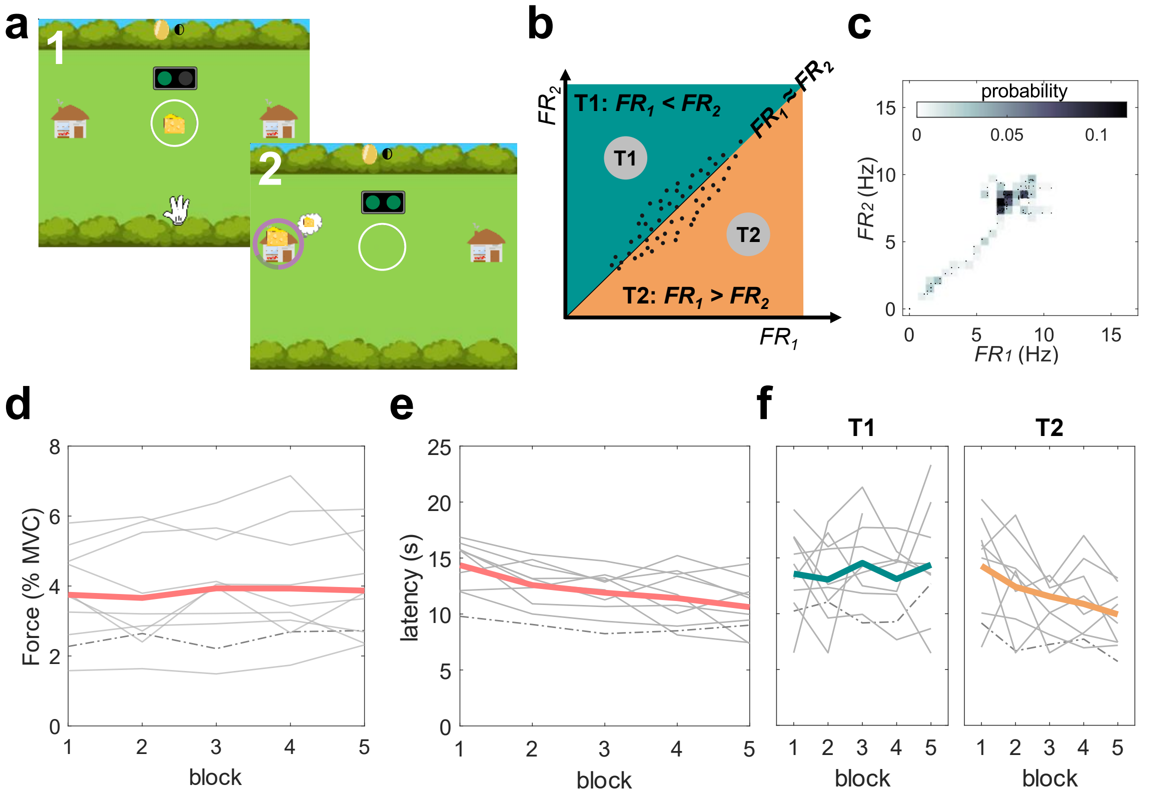


Supplementary Figure S1. (a) Screen images of the experiment. Participants first controlled the movement of a cartoon hand-shaped cursor moving only along the y-axis to pick up the delivery item (cheese) in the baseline area indicated by a white circle (a1). Participants then had to move the delivery to the target house (a2). (b) Schematic diagram of firing rates of control-MUs during the experiment. Black dots show typical correlated firing patterns observed during force modulation when both control-MUs are active. Firing rates of control-MUs were re-scaled to a similar level so that the activity observed during isometric contraction lied approximately on the diagonal. (c) Firing rate distribution from an example trial. Probability of the activities of the controlled-MUs in one trial using a bin width of 0.6 Hz. (d-e) Force (d) and target reach latency (e) across the 5 blocks (individual participants in gray; mean across participants in red). (f) Same as (e) but separately for targets T1 (left) and T2 (right) (individual participants in gray; mean across participants in color). In (d)-(f) the dash-dotted line reflects the participant whose neural data was excluded (see Methods for details).

#### Modulation of firing rates differences across blocks in the difference control task

We examined the firing rates during the baseline and action phases. Note that the firing rates here were adjusted with the gain values. We separately analyzed the sum of the firing rates of the two MUs as well as the difference ($FR_1 - FR_2$) between their firing rates. In the experiment, the firing rate sum controlled the horizontal cursor position while the firing rate difference controlled the vertical cursor velocity. The firing rate difference, thus, needs to be modulated to move the cursor to the targets. All analyses were carried out separately using the MU firing rates during the baseline and the action phases, as well as the rates during the action phase after subtracting the baseline rates.

We found no evidence (beyond anecdotal) for a difference in the sum of the firing rates between the two targets, neither for the action phase nor for the baseline phase nor for the action phase rates after subtracting the baseline rates (Fig. S2, a left: Bayesian Wilcoxon signed-rank test $BF_{10}$ = 0.84; f left: Bayesian Wilcoxon signed-rank test $BF_{10}$ = 0.33; k left: Bayesian Wilcoxon signed-rank test $BF_{10}$ = 1.2). This remained to be the case when analyzing the summed firing rates across blocks (Fig. S2, b: Bayesian repeated measures ANOVA, $BF_{inclusion}$ = 1, 0.13 and 1.2 for target, block and target – block interaction; c: trend-$BF_{10}$ = 1.3; g: Bayesian repeated measures ANOVA, $BF_{inclusion}$ = 0.4, 0.3 and 0.5 for target, block and target – block interaction; h: trend-$BF_{10}$ = 0.43; l: Bayesian repeated measures ANOVA, $BF_{inclusion}$ = 0.95, 0.11 and 0.42 for target, block and target – block interaction; m: trend-$BF_{10}$ = 1.2).

For the firing rate difference, there was no evidence for target specificity during the baseline phase (Fig. S2, f right: Bayesian Wilcoxon signed-rank test $BF_{10}$ = 0.46; i: Bayesian repeated measures ANOVA, $BF_{inclusion}$ = 0.4, 0.6 and 0.3 for target, block and target – block interaction; j: trend-$BF_{10}$ = 0.3). During the action phase, the firing rate differences were target specific (Fig. S2a right, Bayesian Wilcoxon signed-rank test $BF_{10}$ = 14.7) and the target specificity increased slightly across blocks (Fig. S2e, trend-$BF_{10}$ = 40; Fig. S2d, Bayesian repeated measures ANOVA, $BF_{inclusion}$ = 10, 2 and >100 for target, block and target – block interaction) in parallel to the increasing performance (Fig. 7f and Fig. S1f). After subtracting the baseline rates, the firing rate differences remained different between the two targets (Fig. S2k right: Bayesian Wilcoxon signed-rank test $BF_{10}$ >100) and the target specificity continued to increase across blocks (Fig. S2o, trend-$BF_{10}$ = 21; Fig. 2n, Bayesian repeated measures ANOVA, $BF_{inclusion}$ = 8.9, 0.25, 2.8 for target, block and target – block interaction). Yet, visually the changes of the rate differences for T1 and T2 across blocks during the action phase (Fig. S2d) are affected by the subtraction of the baseline phase (Fig. S2n).

Taken together, these results show that the difference but not the sum of the firing rates of the two MUs was target specific. The target specificity of the firing rate differences increased across blocks in parallel to the performance.

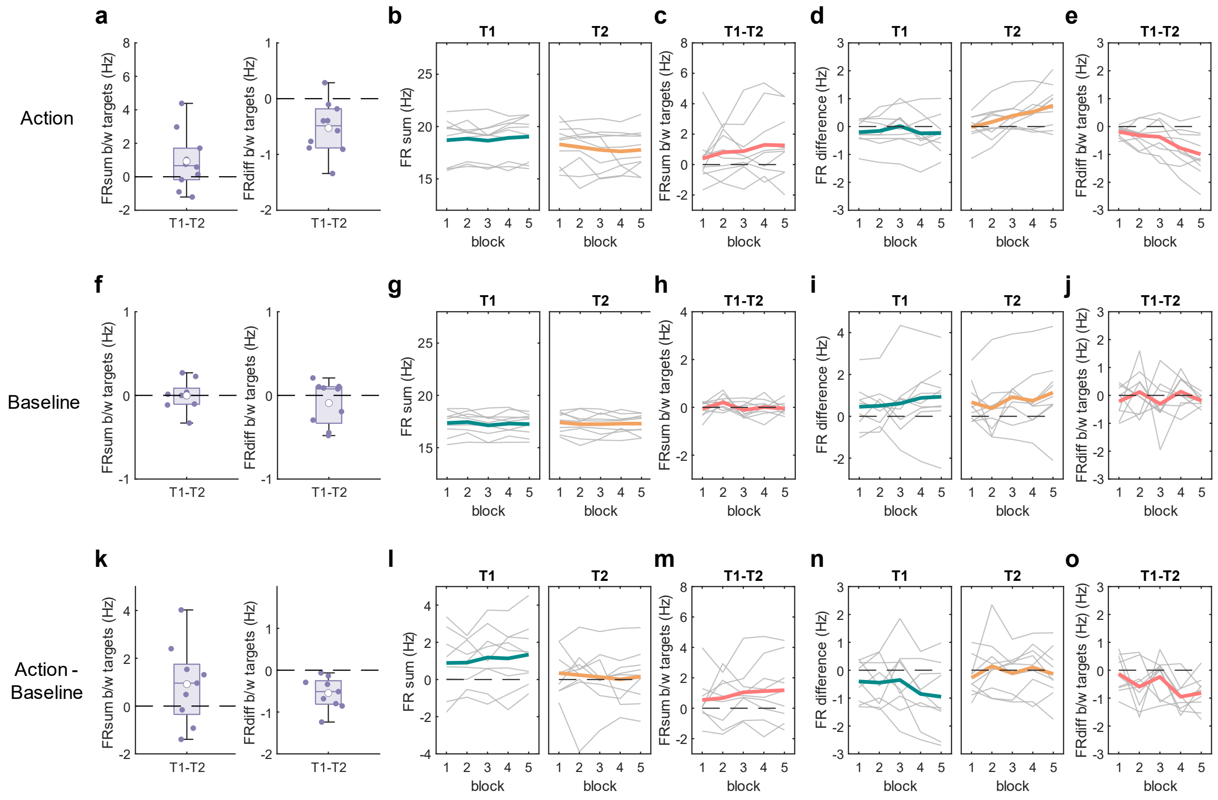


Supplementary Figure S2. (a) Difference of firing rate sum (left) and differences (right) between T1 and T2 during action phase. Each filled dot depicts one participant. White dots depict the mean; horizontal lines the median. (b-e) Firing rate sum (b) and differences (d) across 5 blocks for T1 (left) and T2 (right) trials during the action phase. Differences between the targets across blocks for firing rate sum (c) and difference (e). Individual participants in gray; mean across participants in color. Dashed lines depict zero. (f-j) same as a-e but for baseline phase. (k-o) same as in a-e but after subtraction of baseline rates.

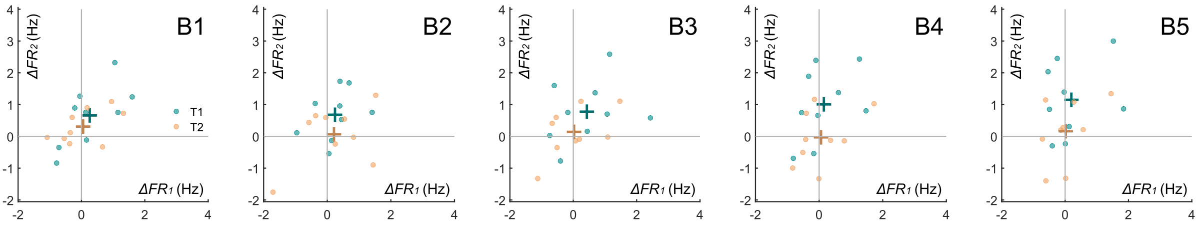


Supplementary Figure S3. Averaged $\Delta FR_1$ and $\Delta FR_2$ separately for each block for the difference control task (dots: individual participants; +: mean across participants).

| | $BF_{20}$ | $BF_{21}$ | $BF_{10}$ |
|---|---|---|---|
| ***T1*** | | | |
| **B1** | **1.1** | **0.7** | **1.5** |
| **B2** | 2.5 | 1.3 | 2 |
| **B3** | 1.9 | 1.7 | 1.1 |
| **B4** | **2.4** | **0.5** | **4.4** |
| **B5** | **3.8** | **0.7** | **5.2** |
| ***T2*** | | | |
| **B1** | 1 | 13 | 0.07 |
| **B2** | 0.16 | 0.7 | 0.24 |
| **B3** | 0.19 | 2.1 | 0.09 |
| **B4** | 0.14 | 0.7 | 0.2 |
| **B5** | 0.18 | 1.7 | 0.1 |

Supplementary Table 3. Bayes factors for difference control experiment for individual targets and blocks.

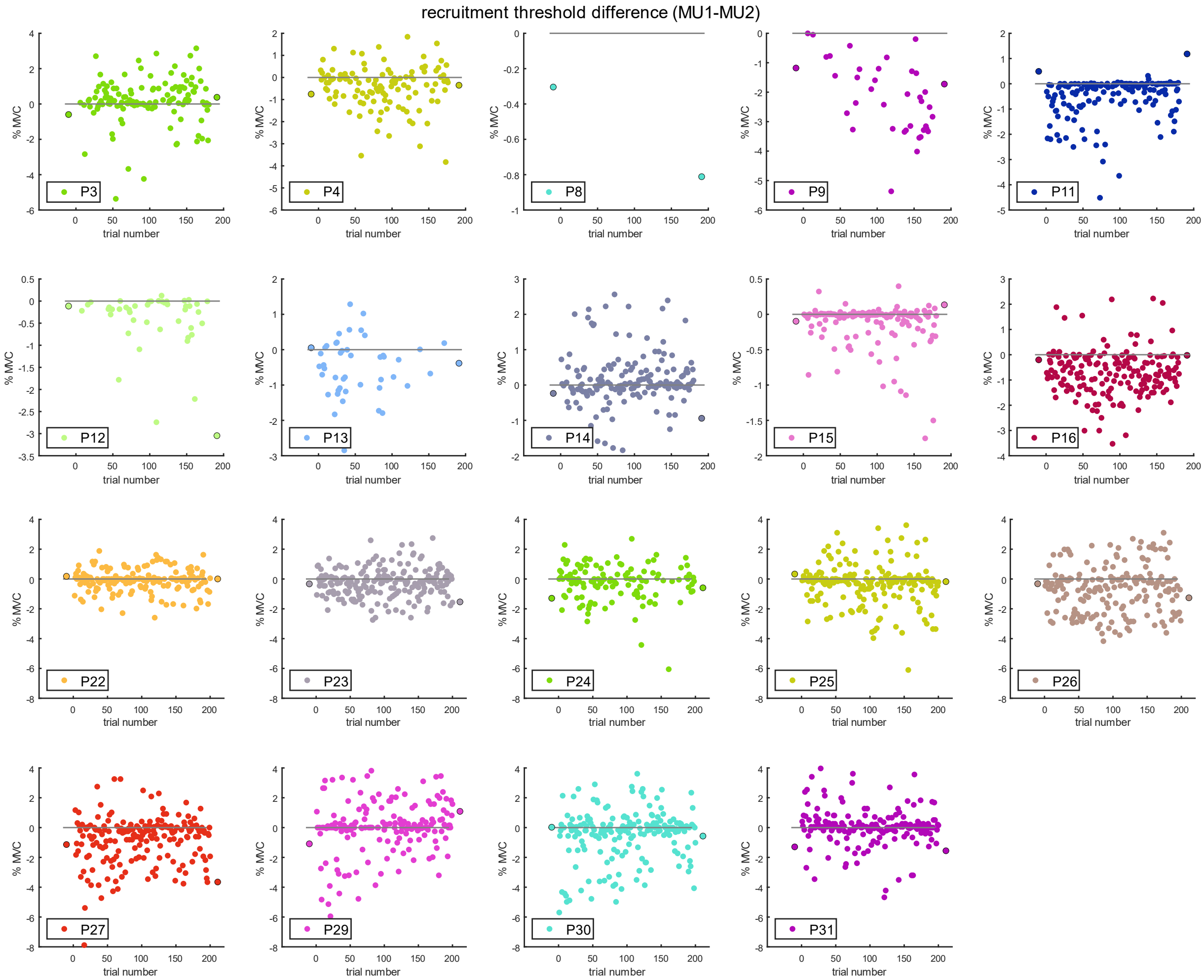


Supplementary Figure S4. Recruitment threshold difference from the two MUs during ramp task and control trials. The filled circles with black edges at the very beginning and end represent the recruitment threshold differences from the ramp tasks conducted before (left) and after (right) the control tasks. The filled circles without edges depict the recruitment threshold differences observed during each trial. Missing data points may be due to either unrecorded data (P8) or the units being already active before the trial began.

#### Robustness Check

The robustness of the neuronal analyses was assessed with regard to the exclusion of participants, the removal of data around derecruitment, the recruitment thresholds, and the time window during the action phase. The results from these robustness checks are presented in the next paragraphs. None of these analyses yielded results supporting successful target reach.

##### ***Inclusion of data from all participants***

Repeating the analysis of firing rate differences including the data from all participants yielded qualitatively the same conclusions as the results from the selected participants for both the displacement and the difference control tasks. (Supplementary Figure S5, Supplementary Tables 4 and 5).

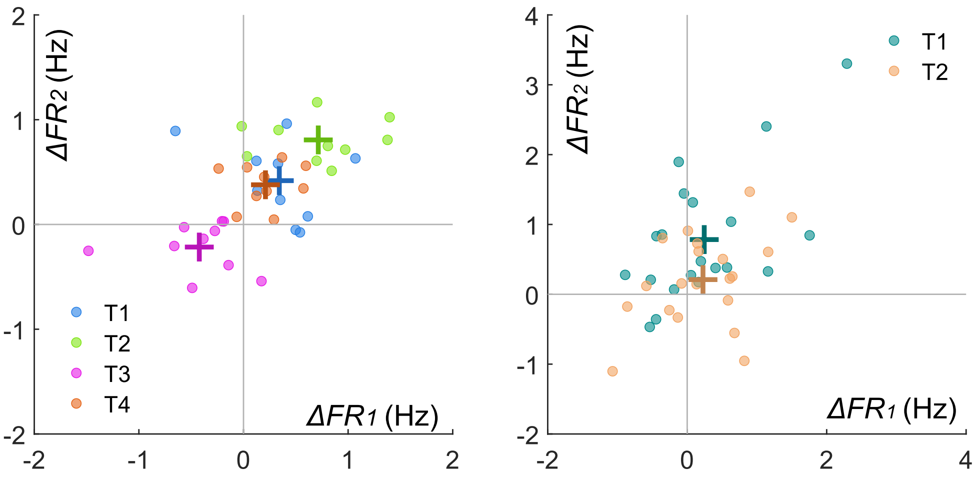


Supplementary Figure S5. $\Delta FR_1$ and $\Delta FR_2$ averaged across all blocks from all participants (dots: individual participants; +: mean across participants). Left: displacement control task experiment. Right: difference control task experiment.

| | **$BF_{20}$** | **$BF_{21}$** | **$BF_{10}$** |
|---|---|---|---|
| **T1** | 263 | 90 | 2.9 |
| **T2** | 5.9 x $10^5$ | 266 | 2231 |
| **T3** | 52 | 34 | 1.6 |
| **T4** | 560 | 12.9 | 43 |

Supplementary Table 4. Bayes factors for the displacement experiment including all participants.

| | **$BF_{20}$** | **$BF_{21}$** | **$BF_{10}$** |
|---|---|---|---|
| **T1** | 24.8 | 1.5 | 16.7 |
| **T2** | 0.3 | 3 | 0.1 |

Supplementary Table 5. Bayes factors for the difference control experiment including all participants.

##### ***Rearrangement of $MU_1$ and $MU_2$***

The re-analysis after rearranging $MU_1$ and $MU_2$ for the displacement control task (see materials and methods section "Determination of recruitment threshold order for $MU_1$ and $MU_2$"), revealed no evidence supporting successful target reach (Supplementary Table 6 and 7, Supplementary Figure S6), thus supporting the main conclusion of the original analyses, though with overall weaker evidence. The re-analysis provided weaker evidence against target reach for T4, amounting to only anecdotal evidence against $H_1$ on its own. For the opposing targets T3 and T4, the evidence from the re-analysis shifted from strong to moderate evidence against $H_1$.

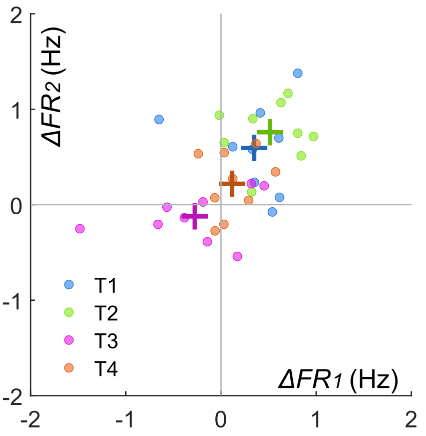


Supplementary Figure S6. $\Delta FR_1$ and $\Delta FR_2$ averaged across all blocks after rearranging $MU_1$ and $MU_2$, ensuring that $MU_2$ is the MU with the higher recruitment threshold for each participant. (dots: individual participants; +: mean across participants).

| | $BF_{20}$ | $BF_{21}$ | $BF_{10}$ |
|---|---|---|---|
| **T1** | 90 | 94 | 0.96 |
| **T2** | 4.6 x $10^3$ | 124 | 37 |
| **T3** | 0.7 | 2.7 | 0.26 |
| **T4** | 1.4 | 2.4 | 0.6 |

Supplementary Table 6. Bayes factors for the displacement task (Table 1 in main text) after rearranging $MU_1$ and $MU_2$, ensuring that $MU_2$ is the motor unit with the higher recruitment threshold for each participant.

| | $BF_{20}$ | $BF_{21}$ | $BF_{10}$ |
|---|---|---|---|
| **T1 & T2** | 8.3 x $10^3$ | 1.4 x $10^3$ | 5.8 |
| **T3 & T4** | 1.4 | 6.2 | 0.2 |

Supplementary Table 7. Bayes Factors for opposing targets in the displacement task (Table 3 in main text) after rearranging $MU_1$ and $MU_2$, ensuring that $MU_2$ is the motor unit with the higher recruitment threshold for each participant.

#### ***Modified trial duration and trial exclusion***

We re-analysed the firing rate changes using only the second half of the action phase (i.e. corresponding to 5-10 s after the onset of the action phase) and excluding trials entirely when one or both MUs were derecruited during the trial (which resulted in total in 1437 out of 1800 trials being used from 9 participants). The results are shown in Supplementary Figure S7 and Supplementary Tables 8-10). For the data aggregated across blocks, none of the targets showed evidence in favour of target reach but the evidence against target reach decreased for T4 compared to the previous analysis (cf. Supplementary Table 8 with Table 1). The re-analysis also yielded moderate to strong evidence against the reaching of opposing targets (Supplementary Table 9). The weaker evidence against target reach in the re-analyses may be the result of the larger variability due to the smaller amounts of data (namely, fewer trials and shorter time windows to estimate the firing rate during the action phase). When the re-analysis was conducted for individual blocks (Supplementary Table 10), the evidence against $H_1$ further declined for several blocks. Moreover, three individual blocks showed anecdotal to moderate evidence for successful target reach, one of which (B3 for T4) consistent with the original analysis (Supplementary Table 1). Given the anecdotal evidence for target reach for two out of these three cases (B2 and B1 for T1) and the issue of multiple comparisons (separate hypotheses tests for each of the five blocks), we consider that the re-analysis of individual blocks as a whole also does not provide evidence supporting target reach.

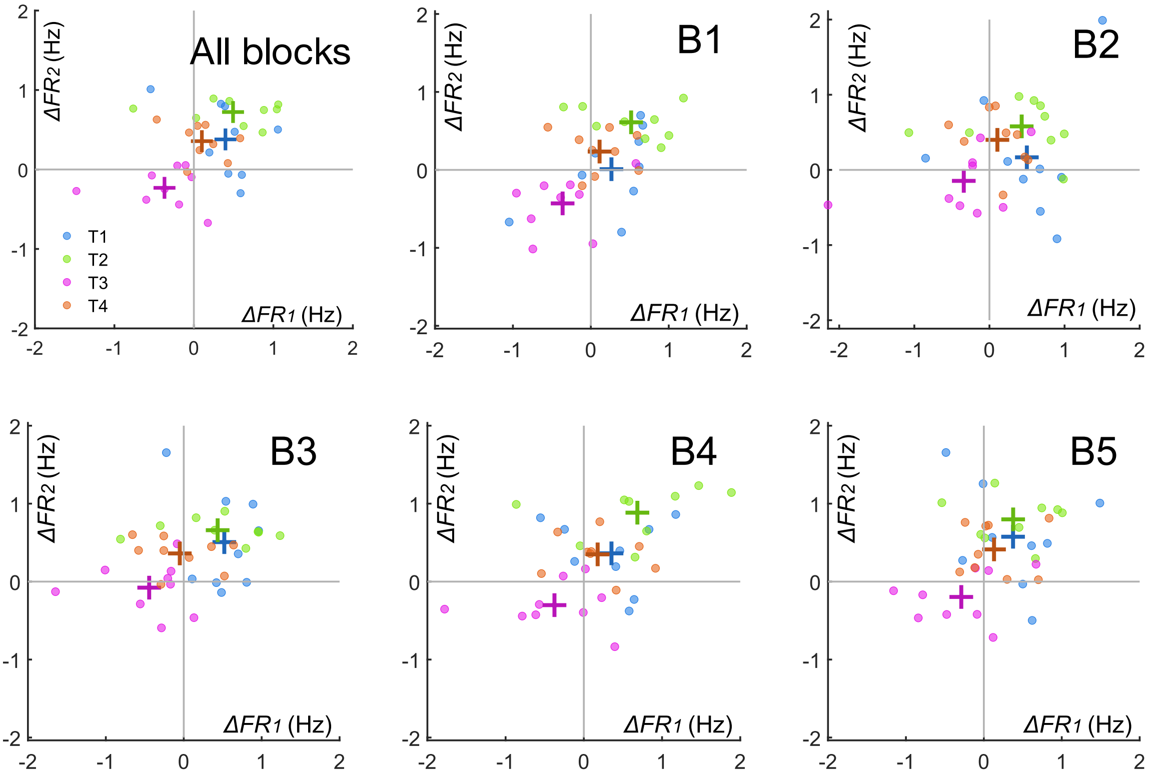


Supplementary Figure S7. $\Delta FR_1$ and $\Delta FR_2$ averaged across blocks and separately for each block for the displacement control task after excluding trials with derecruitment in one or both MUs and considering only the second half of the action phase (dots: individual participants; +: mean across participants).

| | $BF_{20}$ | $BF_{21}$ | $BF_{10}$ |
|---|---|---|---|
| **T1** | 144 | 18 | 8 |
| **T2** | $1.3 \times 10^6$ | 14.3 | $8.9 \times 10^4$ |
| **T3** | 19.5 | 10.4 | 1.9 |
| **T4** | 173 | 1.8 | 95.6 |

Supplementary Table 8. Bayes factors for the displacement task (Table 1 in main text) after excluding trials with derecruitment in one or both MUs and considering only the second half of the action phase.

| | $BF_{20}$ | $BF_{21}$ | $BF_{10}$ |
|---|---|---|---|
| **T1 & T2** | $7.9 \times 10^5$ | 312 | $2.6 \times 10^3$ |
| **T3 & T4** | $3.1 \times 10^5$ | 8.8 | $3.5 \times 10^4$ |

Supplementary Table 9. Bayes Factors for opposing targets in the displacement task (Table 3 in main text) after excluding trials with derecruitment in one or both MUs and considering only the second half of the action phase.

| | $BF_{20}$ | $BF_{21}$ | $BF_{10}$ |
|---|---|---|---|
| ***T1*** | | | |
| **B1** | **0.39** | **0.295** | **1.33** |
| **B2** | **1.2** | **0.74** | **1.55** |
| **B3** | 218 | 14.8 | 14.8 |
| **B4** | 10.8 | 14.3 | 0.75 |
| **B5** | 20.2 | 20 | 1 |
| ***T2*** | | | |
| **B1** | $4 \times 10^4$ | 28.3 | $1.4 \times 10^3$ |
| **B2** | 258 | 5.7 | 45.5 |
| **B3** | $4.9 \times 10^5$ | 6.4 | $7.7 \times 10^4$ |
| **B4** | $3.9 \times 10^3$ | 9.2 | 425 |
| **B5** | $8.2 \times 10^3$ | 8.6 | 956 |
| ***T3*** | | | |
| **B1** | 15.4 | 7 | 2.2 |
| **B2** | 0.46 | 3.03 | 0.15 |
| **B3** | 2.6 | 16 | 0.16 |
| **B4** | 11.8 | 4.1 | 2.9 |
| **B5** | 1.1 | 2.9 | 0.39 |
| ***T4*** | | | |
| **B1** | 3.6 | 1.4 | 2.5 |
| **B2** | 20.8 | 1.7 | 12.6 |
| **B3** | **51** | **0.25** | **202** |
| **B4** | 36.3 | 2 | 18 |
| **B5** | 15.8 | 1.4 | 11.6 |

Supplementary Table 10. Bayes factors for the displacement experiment for individual targets and blocks after excluding trials with derecruitment in one or both MUs and considering only the second half of the action phase.

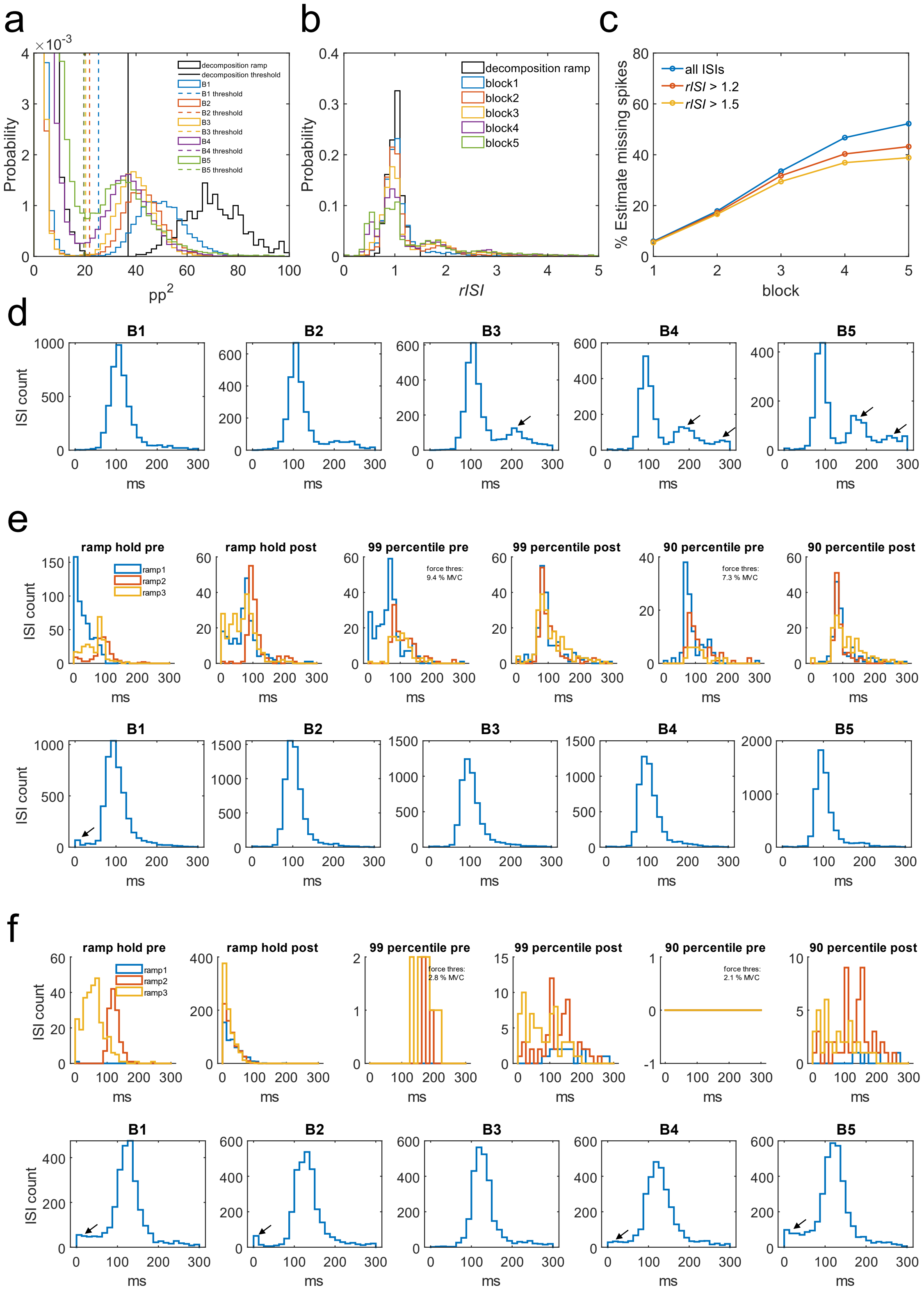


Supplementary Figure S8. (a-d) Data from the same participant. (a) Histograms of peaks of the squared motor unit pulse train (pp$^2$) obtained from recordings during a trapezoid force profile (decomposition ramp) to initialize decomposition algorithm (black) and control blocks (colored). The vertical lines show the thresholds determined by k-means clustering for classification. (b) *rISI* from the decomposition ramp (black) and the control blocks (colored). (c) Estimated fraction of missing spikes without applying *rISI* (blue), only using ISIs with *rISI* >1.2 (red) or *rISI* > 1.5(yellow). (d) Example ISI histograms from a single participant during control blocks. The second and third peaks showing in the histogram (black arrows) indicate missing spikes. (e-f) Example ISI histograms from two participants during trapezoid force tasks performed before and after the control blocks (upper row) and during control blocks (bottom row). The trapezoid force tasks were repeated three times before and after the control blocks, with each repetition labeled as ramp1 to ramp3. The ISI histograms show ISIs from either before (pre) or after (post) the control blocks using different parts of the data. The two subplots on the right show ISI histograms from the hold part of the ramps. The other subplots include ISIs that were extracted from the ramp-up and ramp-down phase where the force level is either below the 99th and 90th percentile of forces recorded during the control blocks. Black arrows point at peaks of short ISIs in the ISI histograms in the control blocks. B1 to B5 refer to block 1 through block 5, respectively.

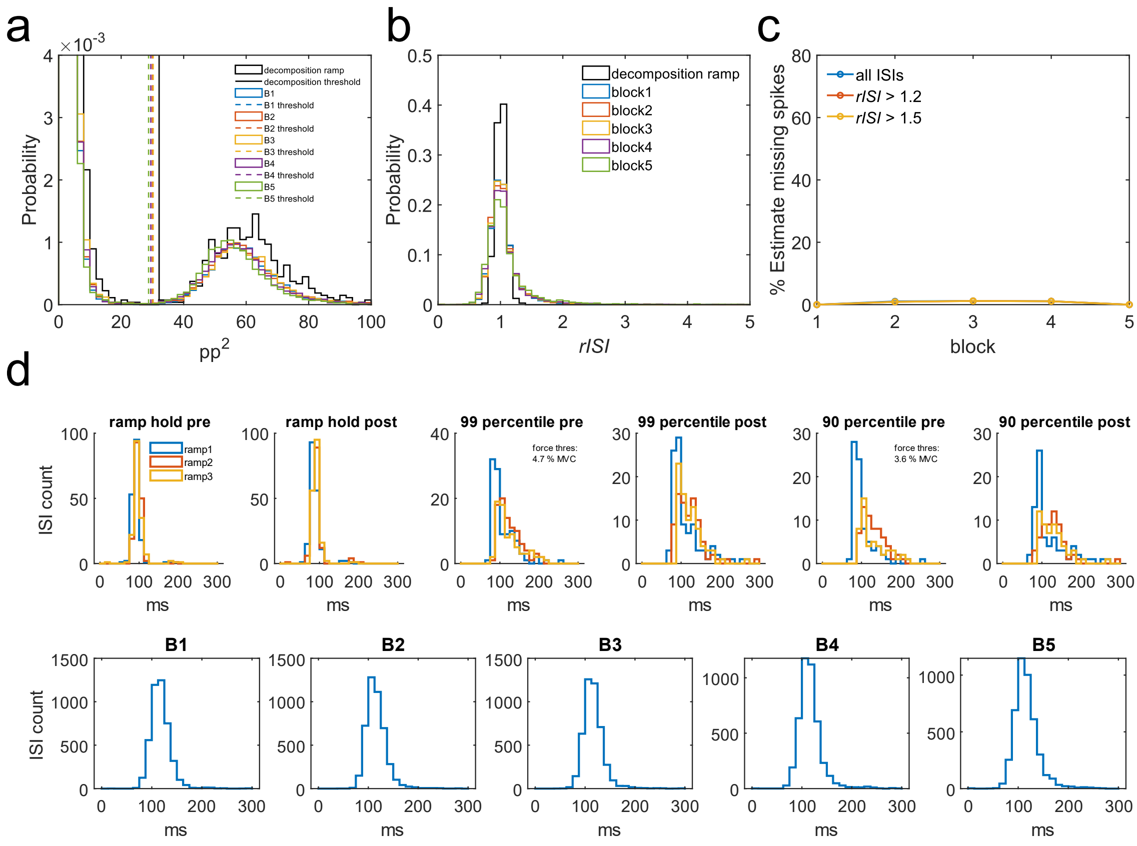


Supplementary Figure S9. Example participant with high decomposition stability. (a-c) Same as Supplementary Figure S8 (a-c). (d) Same as Supplementary Figure S8 (e) and (f).

| Figure label | Figure description | Default prior | Wide prior | Ultra-wide prior |
|---|---|---|---|---|
| Fig3a | MU recruitment / derecruitment | 1.71 / 0.37 | 1.63 / 0.27 | 1.12 / 0.21 |
| Fig3b | Force across blocks | trend-$BF_{10}$ = 0.4 | trend-$BF_{10}$ = 0.3 | trend-$BF_{10}$ = 0.2 |
| Fig3c | Force across blocks per target | $BF_{inclusion}$>100 for target<br>$BF_{inclusion}$= 0.4 for phase<br>$BF_{inclusion}$= 4 for target-phase interaction | $BF_{inclusion}$ >100 for target<br>$BF_{inclusion}$= 0.3 for phase<br>$BF_{inclusion}$= 3.8 for target-phase interaction | $BF_{inclusion}$ >100 for target<br>$BF_{inclusion}$= 0.2 for phase<br>$BF_{inclusion}$= 3.2 for target-phase interaction |
| Fig3d | Time in target | $BF_{inclusion}$= 0.5 for target | $BF_{inclusion}$= 0.4 for target | $BF_{inclusion}$= 0.2 for target |
| Fig3e | Time in target across blocks | trend-$BF_{10}$ = 0.6 | trend-$BF_{10}$ = 0.7 | trend-$BF_{10}$ = 0.4 |
| Fig3f | Time in target across blocks per target | trend-$BF_{inclusion}$= 0.9 for target | trend-$BF_{inclusion}$= 0.7 for target | trend-$BF_{inclusion}$= 0.4 for target |
| Fig5c | s.d. of of $\Delta$FR difference across blocks | trend-$BF_{10}$ = 0.4 | trend-$BF_{10}$ = 0.4 | trend-$BF_{10}$ = 0.3 |
| Fig5d | s.d. of of $\Delta$FR sum across blocks | trend-$BF_{10}$ = 0.3 | trend-$BF_{10}$ = 0.2 | trend-$BF_{10}$ = 0.2 |
| Fig5e | s.d. of of $\Delta$FR difference across blocks per target | trend-$BF_{inclusion}$= 0.6 for target | trend-$BF_{inclusion}$= 0.4 for target | trend-$BF_{inclusion}$= 0.3 for target |
| Fig5f | s.d. of of $\Delta$FR sum across blocks per target | trend-$BF_{inclusion}$= 0.7 for target | trend-$BF_{inclusion}$= 0.5 for target | trend-$BF_{inclusion}$= 0.3 for target |
| Fig7e | MU recruitment / derecruitment | $BF_{10}$ = 1.98 / 0.51 | $BF_{10}$ = 2.08 / 0.4 | $BF_{10}$ = 2.17 / 0.33 |
| Fig7f-left | Success rate across blocks | trend-$BF_{10}$ >100 | trend-$BF_{10}$ > 100 | trend-$BF_{10}$ = 56 |
| Fig7f-right T1/T2 | T1 Success rate across blocks | trend-$BF_{10}$ = 0.3 / >100 | trend-$BF_{10}$ = 0.2 / 96 | trend-$BF_{10}$ = 0.2 / 86 |
| Fig7g-left | FR difference between the 2 MUs in 2 target blocks across blocks | trend-$BF_{10}$ = 40 | trend-$BF_{10}$ = 28.3 | trend-$BF_{10}$ = 24.8 |
| S1d | Force across blocks | trend-$BF_{10}$ = 0.5 | trend-$BF_{10}$ = 0.4 | trend-$BF_{10}$ = 0.3 |
| S1e | Latency across blocks | trend-$BF_{10}$ >100 | trend-$BF_{10}$ = 74 | trend-$BF_{10}$ = 65 |
| S1f | Latency across blocks by target | trend-$BF_{10}$ = 8 | trend-$BF_{10}$ = 7 | trend-$BF_{10}$ = 7 |
| S2a-left / right | FR difference from the two targets during action | $BF_{10}$ = 0.8 / 14.7 | $BF_{10}$ = 0.7 / 12.6 | $BF_{10}$ = 0.6 / 8.9 |
| S2b | FR sum across blocks by target during action | $BF_{inclusion}$= 1 for target<br>$BF_{inclusion}$= 0.13 for block<br>$BF_{inclusion}$= 1.2 for target-block interaction | $BF_{inclusion}$= 1.1 for target<br>$BF_{inclusion}$= 0.06 for block<br>$BF_{inclusion}$= 0.9 for target-block interaction | $BF_{inclusion}$= 1.1 for target<br>$BF_{inclusion}$= 0.03 for block<br>$BF_{inclusion}$= 0.4 for target-block interaction |
| S2c | FR sum T1-T2 across blocks during action | trend-$BF_{10}$ = 1.3 | trend-$BF_{10}$ = 1.3 | trend-$BF_{10}$ = 1.1 |
| S2d | FR difference across blocks by target during action | $BF_{inclusion}$= 10 for target<br>$BF_{inclusion}$= 2 for block<br>$BF_{inclusion}$>100 for target-block interaction | $BF_{inclusion}$= 11 for target<br>$BF_{inclusion}$= 1.7 for block<br>$BF_{inclusion}$>100 for target-block interaction | $BF_{inclusion}$= 11 for target<br>$BF_{inclusion}$= 0.9 for block<br>$BF_{inclusion}$>100 for target-block interaction |
| S2d T1 / T2 | | trend-$BF_{10}$ = 0.4 / 85 | trend-$BF_{10}$ = 0.3 / 42 | trend-$BF_{10}$ = 0.2 / 41 |
| S2e (same as 7g-left) | FR difference T1-T2 across blocks during action | trend-$BF_{10}$ = 40 | trend-$BF_{10}$ = 28.3 | trend-$BF_{10}$ = 24.8 |
| S2f-left / right | FR difference from the two targets during baseline | $BF_{10}$ = 0.33 / 0.46 | $BF_{10}$ = 0.25 / 0.38 | $BF_{10}$ = 0.18 / 0.3 |
| S2g | FR sum across blocks by target during baseline | $BF_{inclusion}$= 0.4 for target<br>$BF_{inclusion}$= 0.3 for block<br>$BF_{inclusion}$= 0.5 for target-block interaction | $BF_{inclusion}$= 0.3 for target<br>$BF_{inclusion}$= 0.2 for block<br>$BF_{inclusion}$= 0.4 for target-block interaction | $BF_{inclusion}$= 0.2 for target<br>$BF_{inclusion}$= 0.09 for block<br>$BF_{inclusion}$= 0.2 for target-block interaction |
| S2h | FR sum T1-T2 across blocks during baseline | trend-$BF_{10}$ = 0.43 | trend-$BF_{10}$ = 0.35 | trend-$BF_{10}$ = 0.26 |
| S2i | FR difference across blocks by target during baseline | $BF_{inclusion}$= 0.4 for target<br>$BF_{inclusion}$= 0.6 for block<br>$BF_{inclusion}$= 0.3 for target-block interaction | $BF_{inclusion}$= 0.3 for target<br>$BF_{inclusion}$= 0.4 for block<br>$BF_{inclusion}$= 0.2 for target-block interaction | $BF_{inclusion}$= 0.2 for target<br>$BF_{inclusion}$= 0.3 for block<br>$BF_{inclusion}$= 0.08 for target- block interaction |
| S2j | FR difference T1-T2 across blocks during baseline | trend-$BF_{10}$ = 0.3 | trend-$BF_{10}$ = 0.25 | trend-$BF_{10}$ = 0.18 |
| S2k-left / right | FR difference from the two targets action-baseline | $BF_{10}$ = 1.2 / >100 | $BF_{10}$ = 1 / 74 | $BF_{10}$ = 0.7 / 67 |
| S2l | FR sum across blocks by target during action-baseline | $BF_{inclusion}$= 0.95 for target<br>$BF_{inclusion}$= 0.11 for block<br>$BF_{inclusion}$= 0.42 for target- block interaction | $BF_{inclusion}$= 1.07 for target<br>$BF_{inclusion}$= 0.05 for block<br>$BF_{inclusion}$= 0.24 for target- block interaction | $BF_{inclusion}$= 1.08 for target<br>$BF_{inclusion}$= 0.02 for block<br>$BF_{inclusion}$= 0.15 for target- block interaction |
| S2m | FR sum T1-T2 across blocks during action-baseline | trend-$BF_{10}$ = 1.2 | trend-$BF_{10}$ = 1.2 | trend-$BF_{10}$ = 0.9 |

| | | | | |
|---|---|---|---|---|
| S2n | FR difference across blocks by target during action-baseline | $BF_{inclusion}$= 8.9 for target<br>$BF_{inclusion}$= 0.25 for block<br>$BF_{inclusion}$= 2.8 for target-block interaction | $BF_{inclusion}$= 8.9 for target<br>$BF_{inclusion}$= 0.15 for block<br>$BF_{inclusion}$= 2.4 for target-block interaction | $BF_{inclusion}$= 8.6 for target<br>$BF_{inclusion}$= 0.07 for block<br>$BF_{inclusion}$= 1.8 for target-block interaction |
| S2n T1 / T2 | | trend-$BF_{10}$ = 1 / 0.3 | trend-$BF_{10}$ = 0.98 / 0.2 | trend-$BF_{10}$ = 0.8/ 0.18 |
| S2o | FR difference T1-T2 across blocks during action-baseline | trend-$BF_{10}$ = 21 | trend-$BF_{10}$ = 18 | trend-$BF_{10}$ = 17 |
| - | Delivery items success rate | trend-$BF_{inclusion}$= 0.26 for item | trend-$BF_{inclusion}$= 0.14 for item | trend-$BF_{inclusion}$= 0.06 for item |

Supplementary Table 11. Bayes factor robustness check with various Cauchy prior scales.
For the Bayesian Wilcoxon signed-rank sum test, the prior scales used were: default ($r = \frac{1}{\sqrt{2}}$), wide ($\gamma$ = 1), and ultra-wide ($\gamma$ = $\sqrt{2}$). For the Bayesian repeated measures ANOVA, the priors scales used were: default ($\gamma$ = 0.5), wide ($r = \frac{1}{\sqrt{2}}$), and ultra-wide ($\gamma$ = 1) for the fixed effects with the prior scale for random effects set to twice that of the fixed effects.

**Post Hoc comparison -Target**

| | | Prior Odds | Posterior Odds | $BF_{10,U}$ |
|---|---|---|---|---|
| T1 | T2 | 0.414 | 1.511 | 3.647 |
| | T3 | 0.414 | 7118.624 | 17185.879 |
| | T4 | 0.414 | 1255.354 | 3030.692 |
| T2 | T3 | 0.414 | 711.487 | 1717.681 |
| | T4 | 0.414 | 4439.311 | 10717.446 |
| T3 | T4 | 0.414 | 0.135 | 0.326 |

**Post Hoc comparison -Phase**

| | | Prior Odds | Posterior Odds | $BF_{10,U}$ |
|---|---|---|---|---|
| Baseline | Action | 1.000 | 0.212 | 0.212 |

Supplementary Table 12. Post hoc tests from Fig. 3c were conducted in JASP. The posterior odds have been corrected for multiple testing by fixing the prior probability that the null hypothesis holds across all comparisons according to Westfall[1]. Individual comparisons are based on the Bayesian t-test with a Cauchy prior with $r = \frac{1}{\sqrt{2}}$. The "U" in the Bayes factor denotes that it is uncorrected.

[1] Westfall, P.H., *Multiple testing of general contrasts using logical constraints and correlations.* Journal of the American Statistical Association, 1997. **92**(437): p. 299-306.

**Supplementary Figures S10–S40** present the decomposition stability analysis for the two control MUs of each participant. Supplementary figures S10–S28 present participants with stable decomposition that were included in the main analysis: S10-19 from the difference control task and S20–S28 from the displacement control task. Supplementary figures S29–S39 present participants from the difference control task that were excluded from the main analysis, and S40 presents the participant from the displacement control task that was excluded from the main analysis. Each figure follows the same format as Supplementary Figure S9, with panels (a–d) displaying data from $MU_1$ and panels (e–h) from $MU_2$. A detailed explanation of each subplot can be found in the caption of Supplementary Figure S8.

### Included participants-difference control

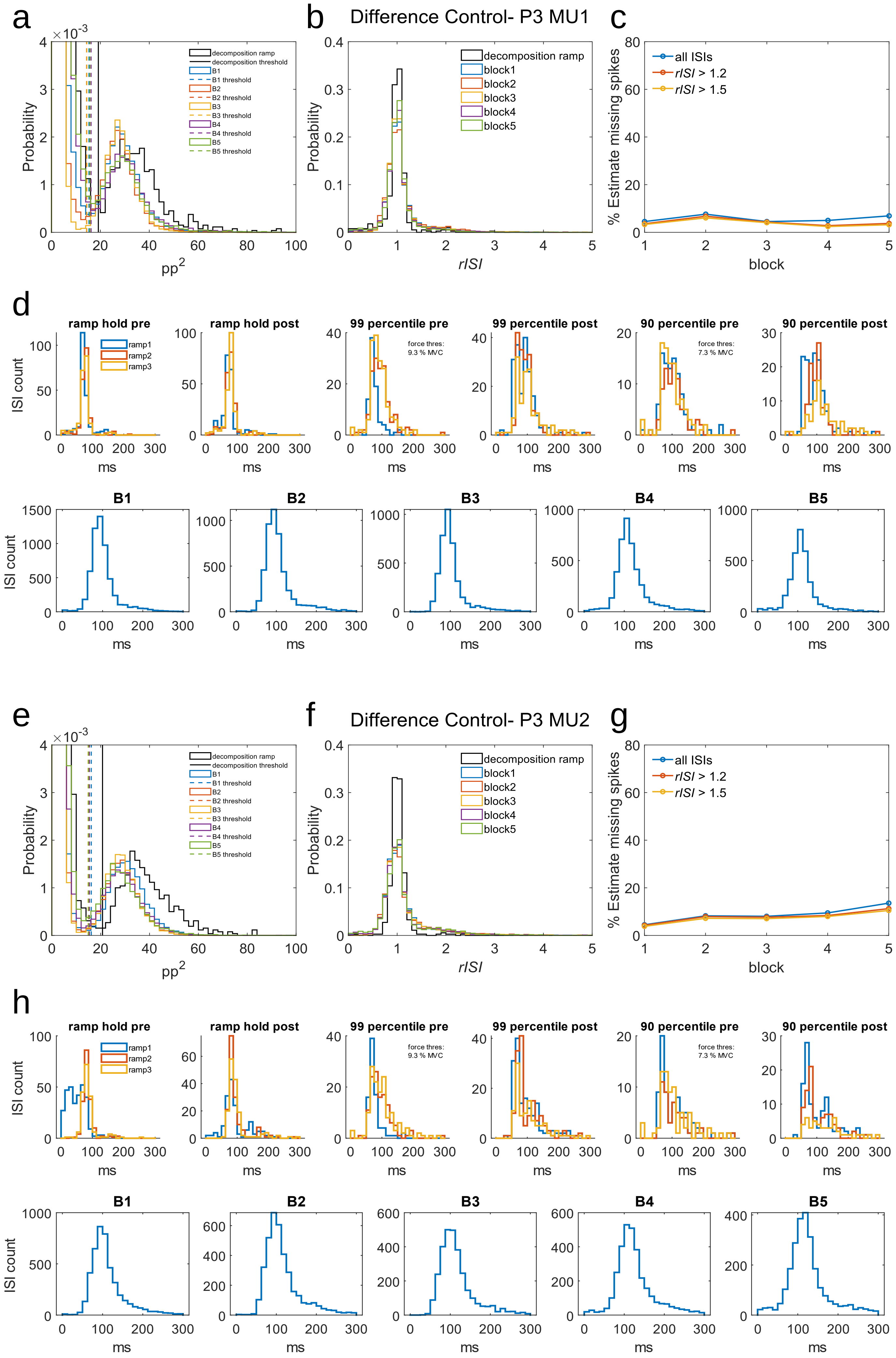


Supplementary Figure S10

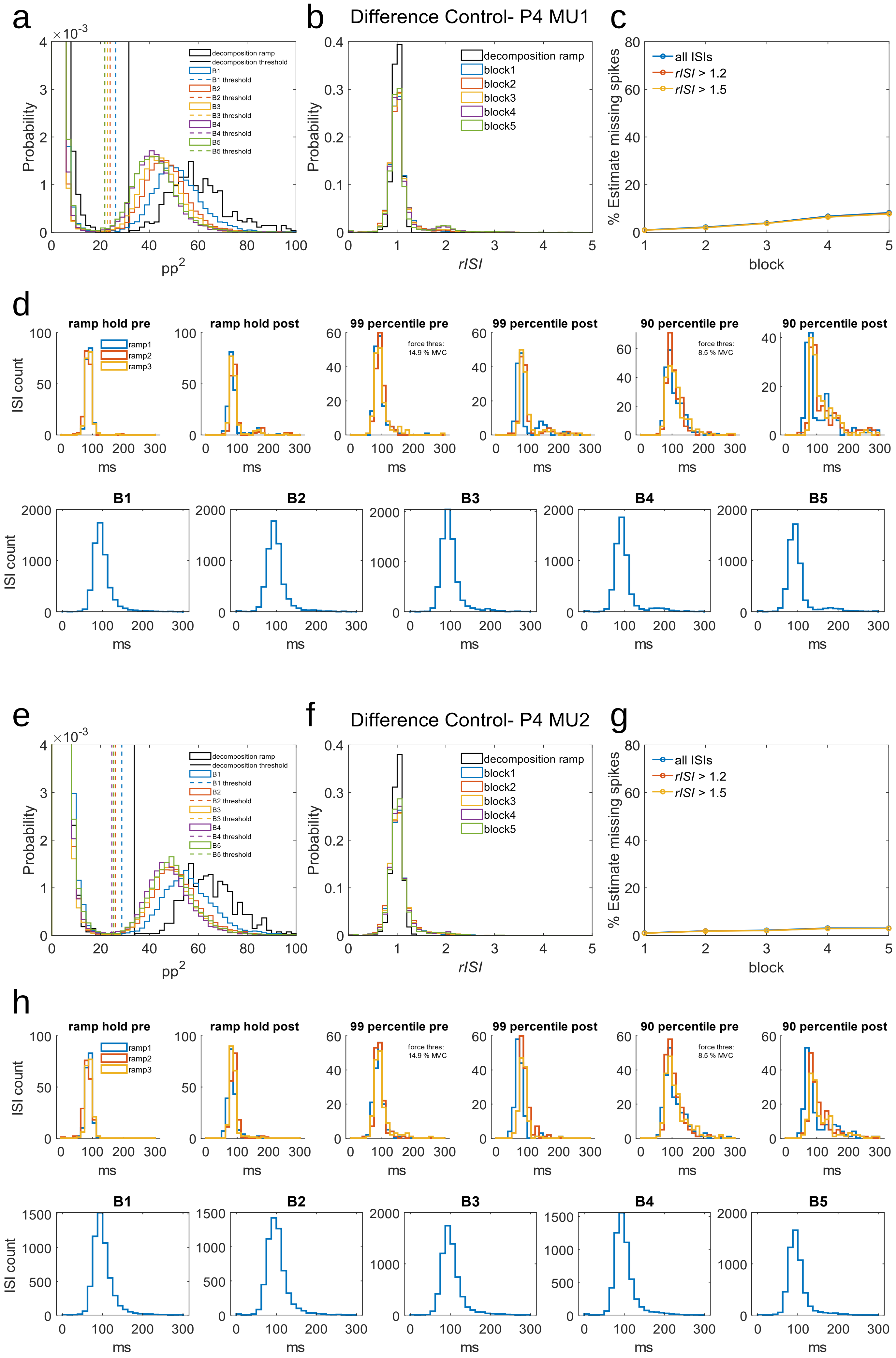


Supplementary Figure S11

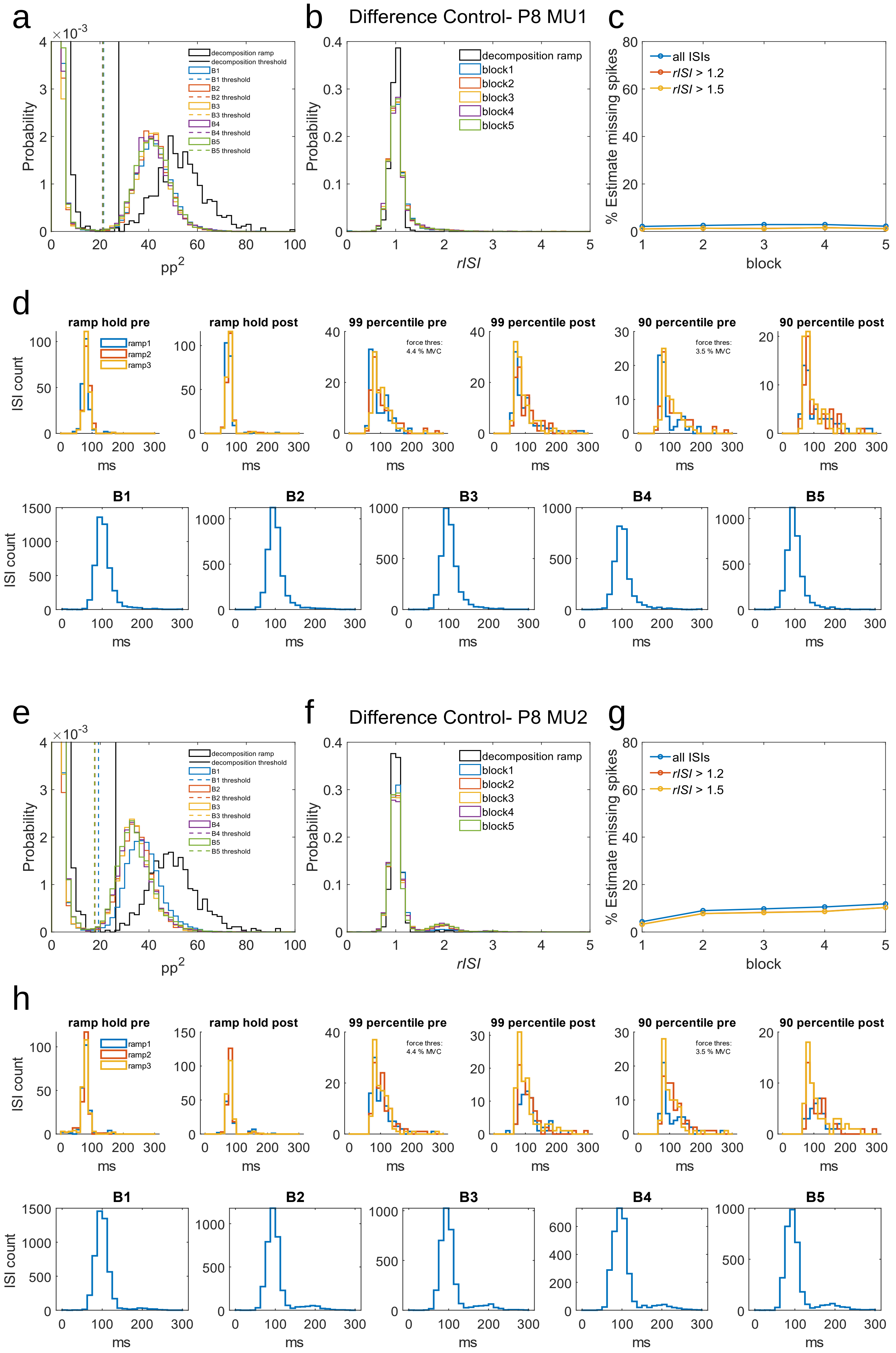


Supplementary Figure S12

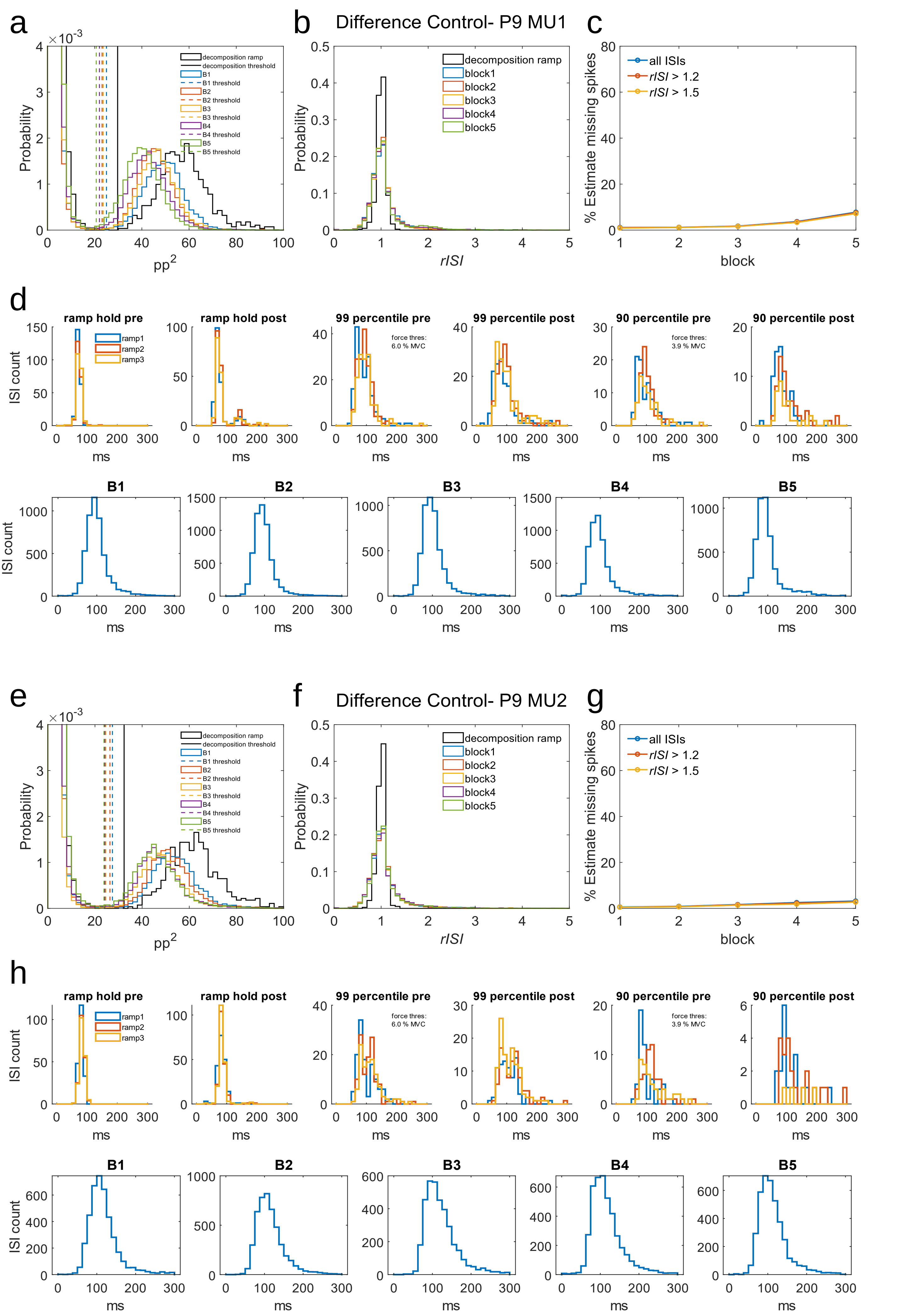


Supplementary Figure S13

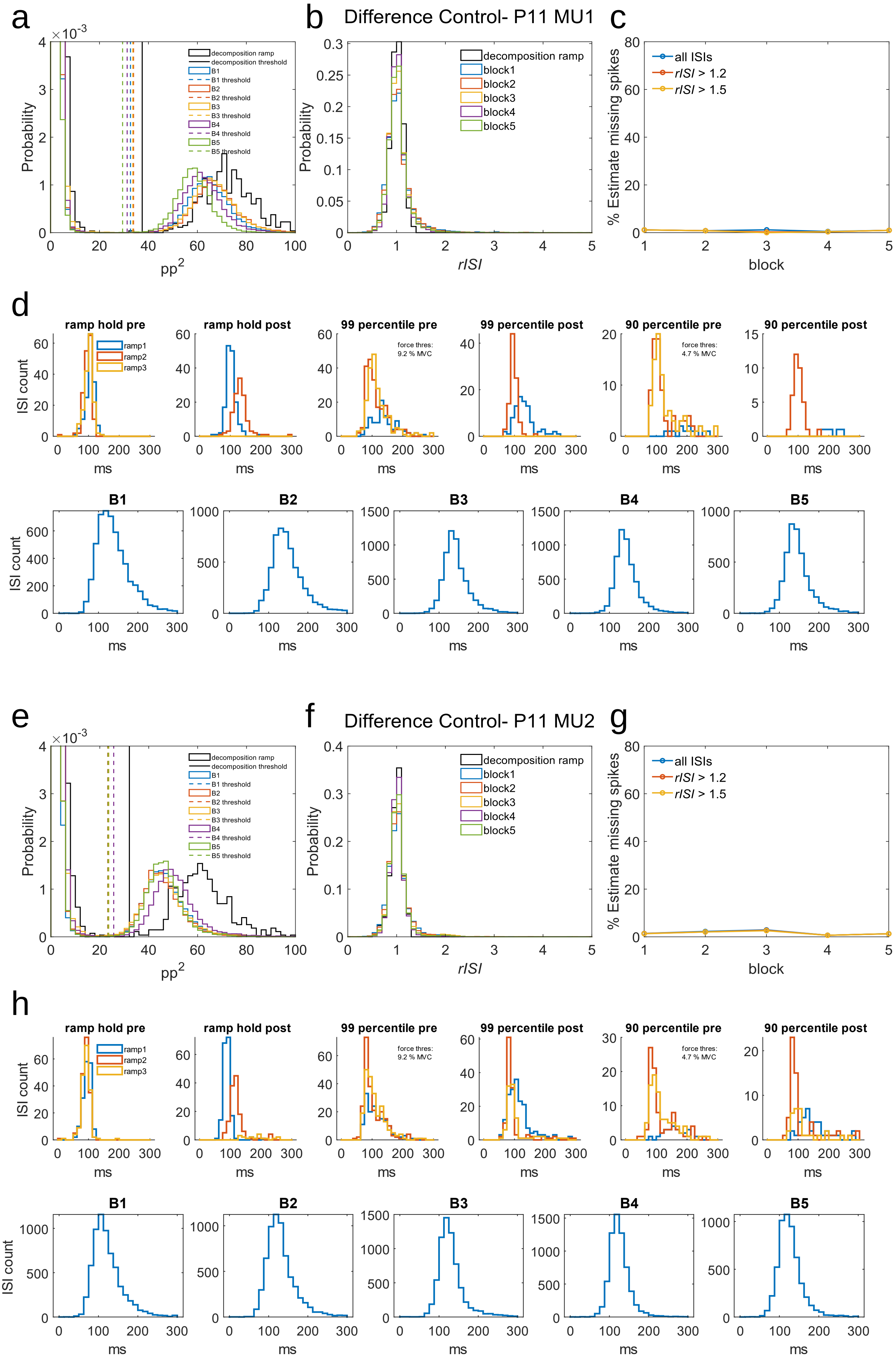


Supplementary Figure S14

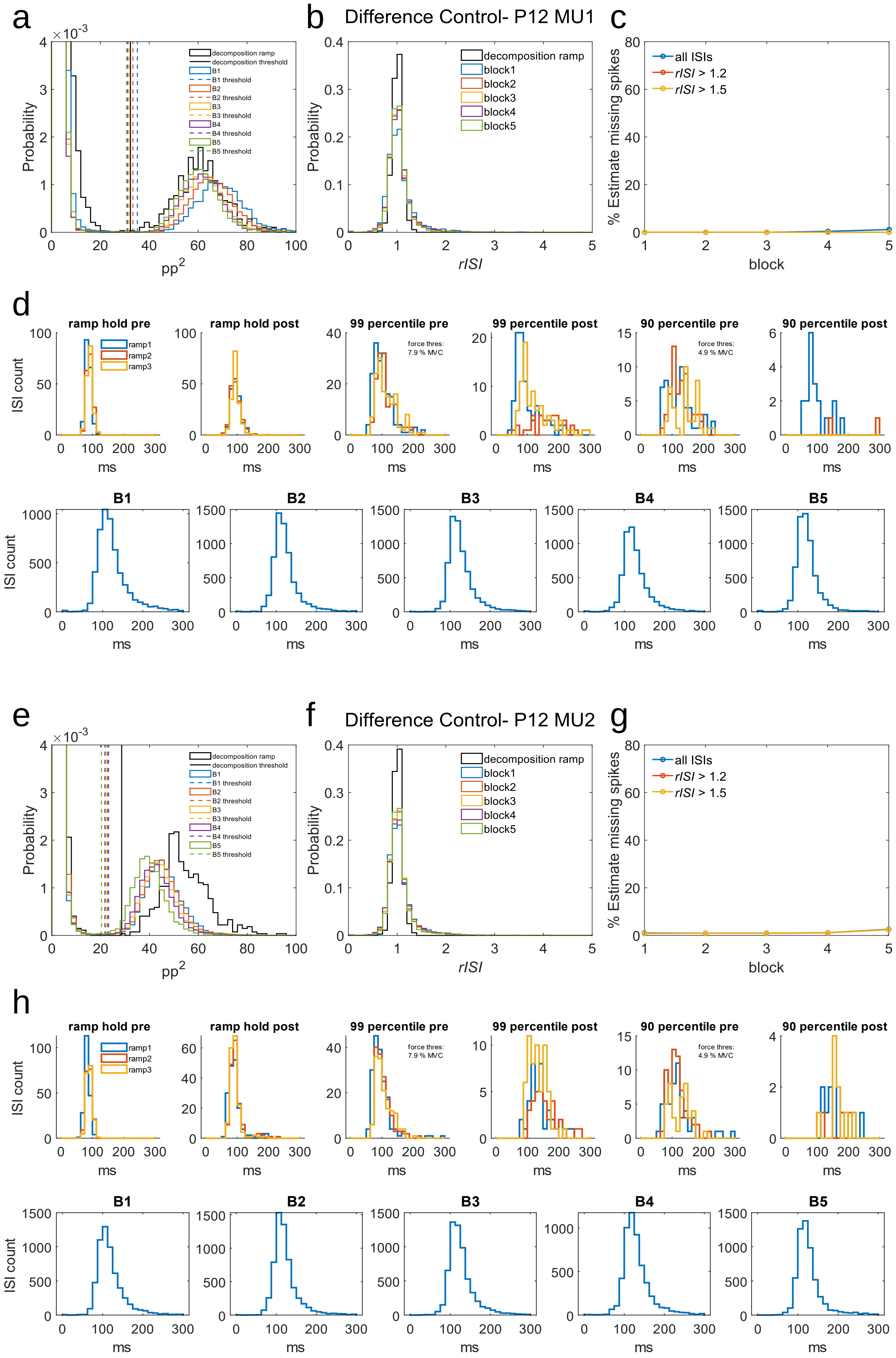


Supplementary Figure S15

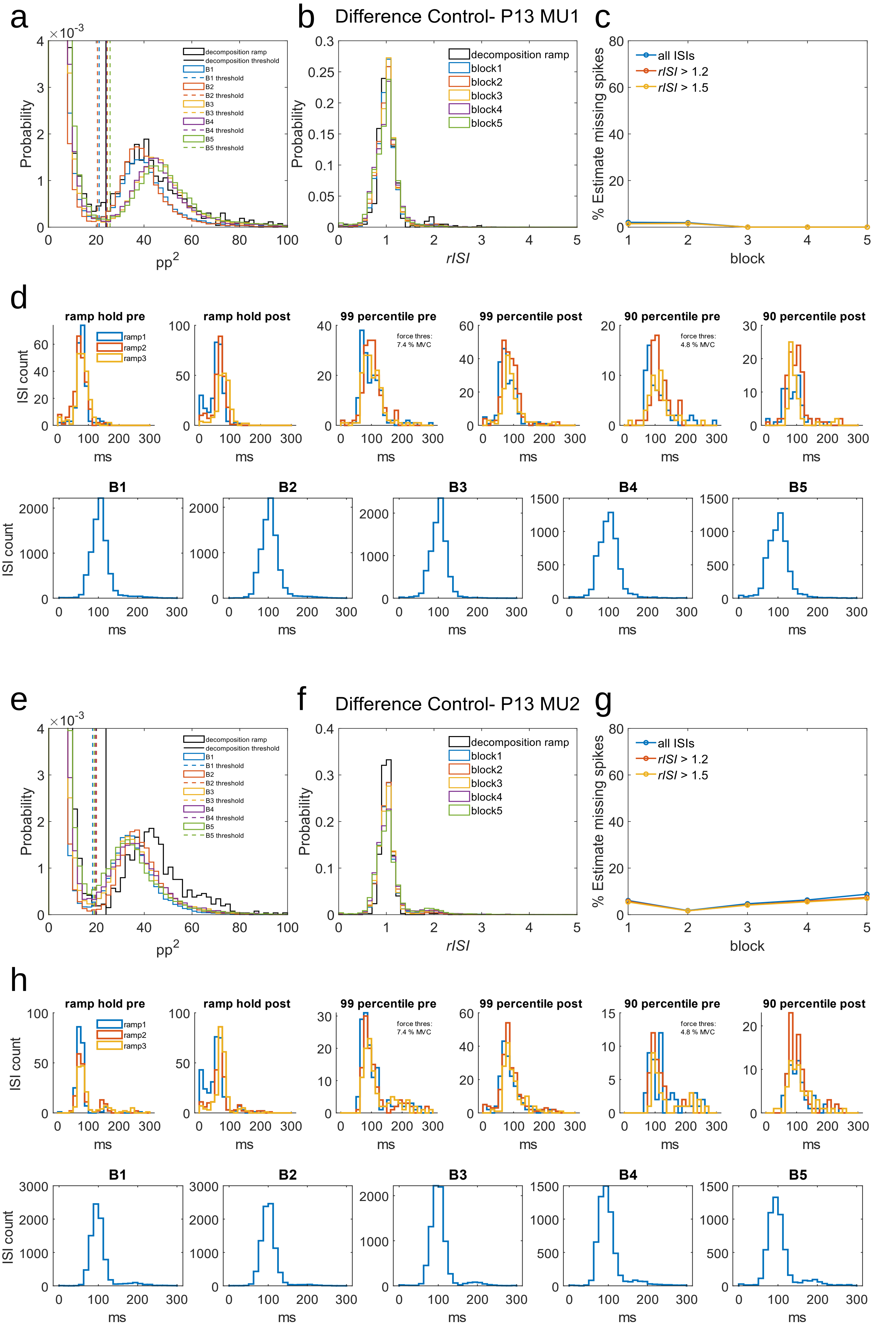


Supplementary Figure S16

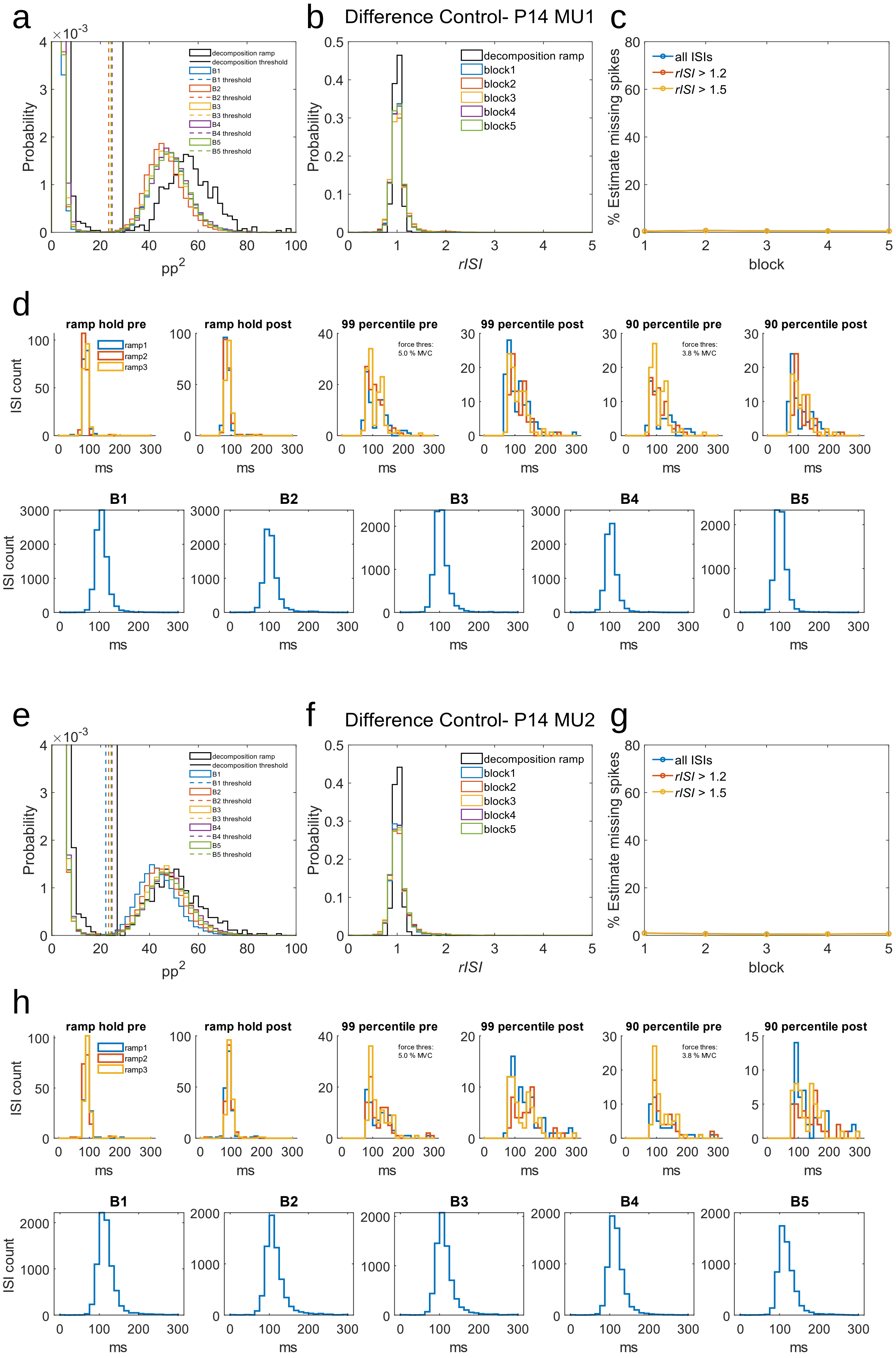


Supplementary Figure S17

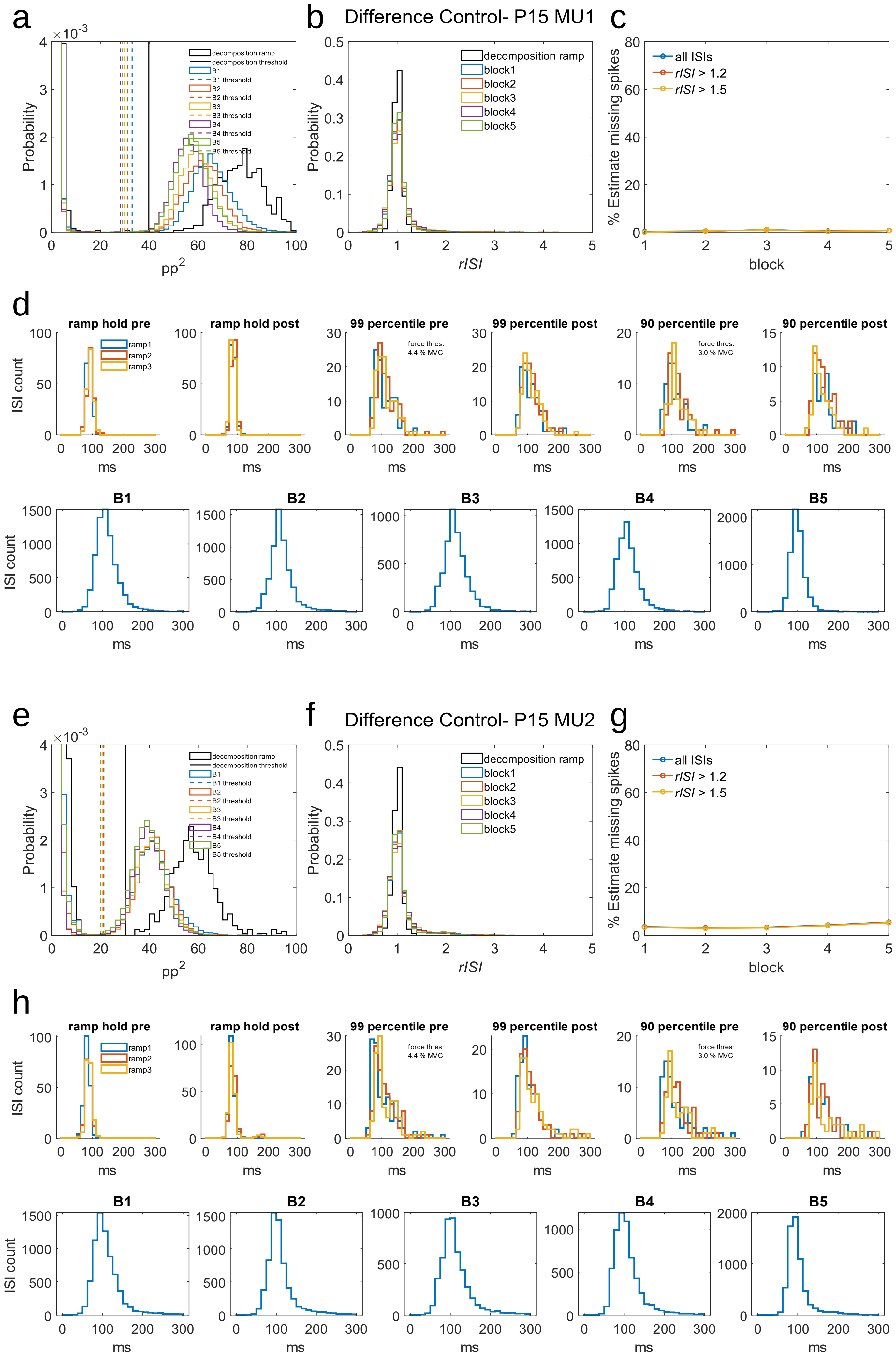


Supplementary Figure S18

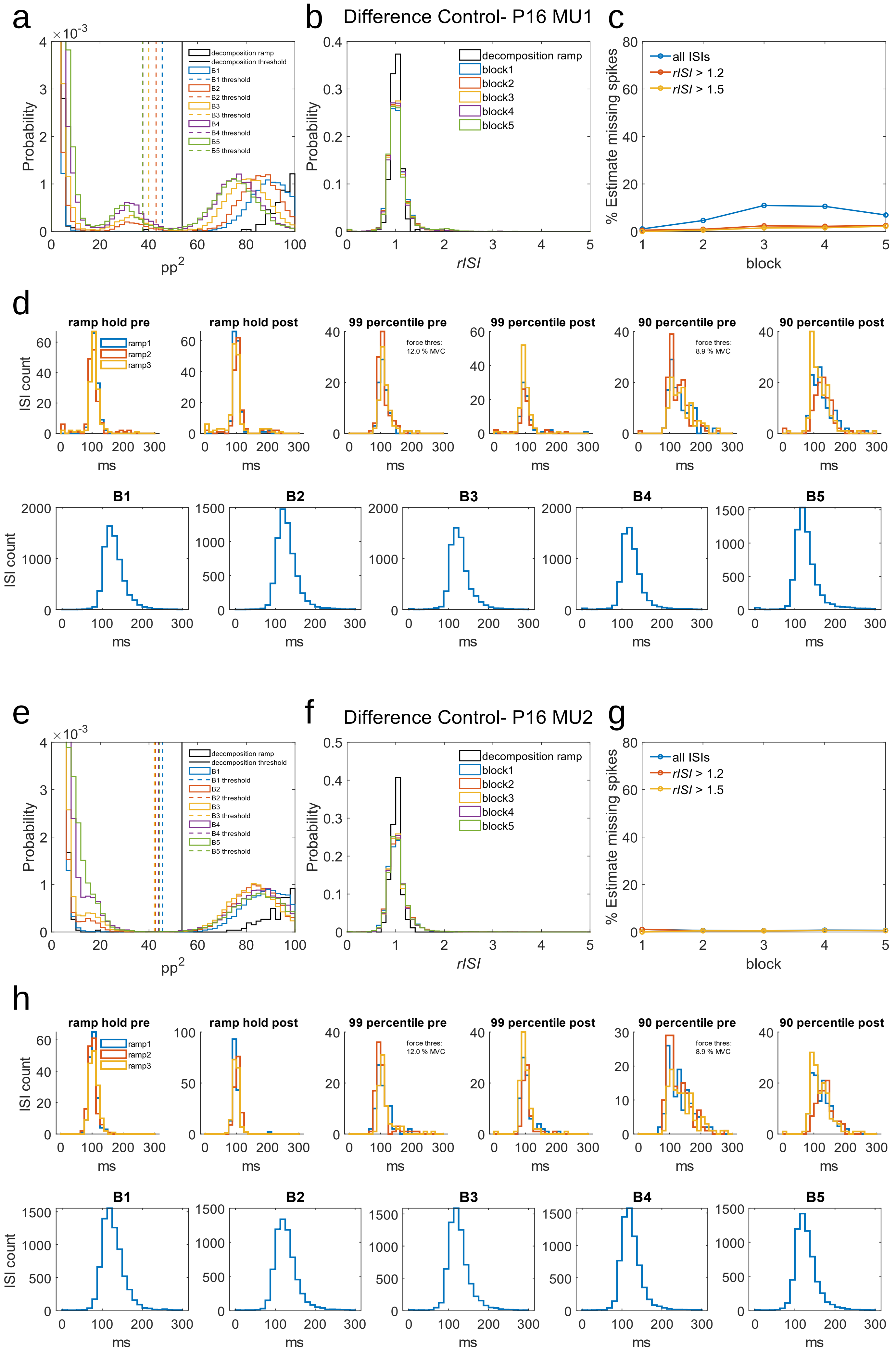


Supplementary Figure S19

### Included participants-displacement control

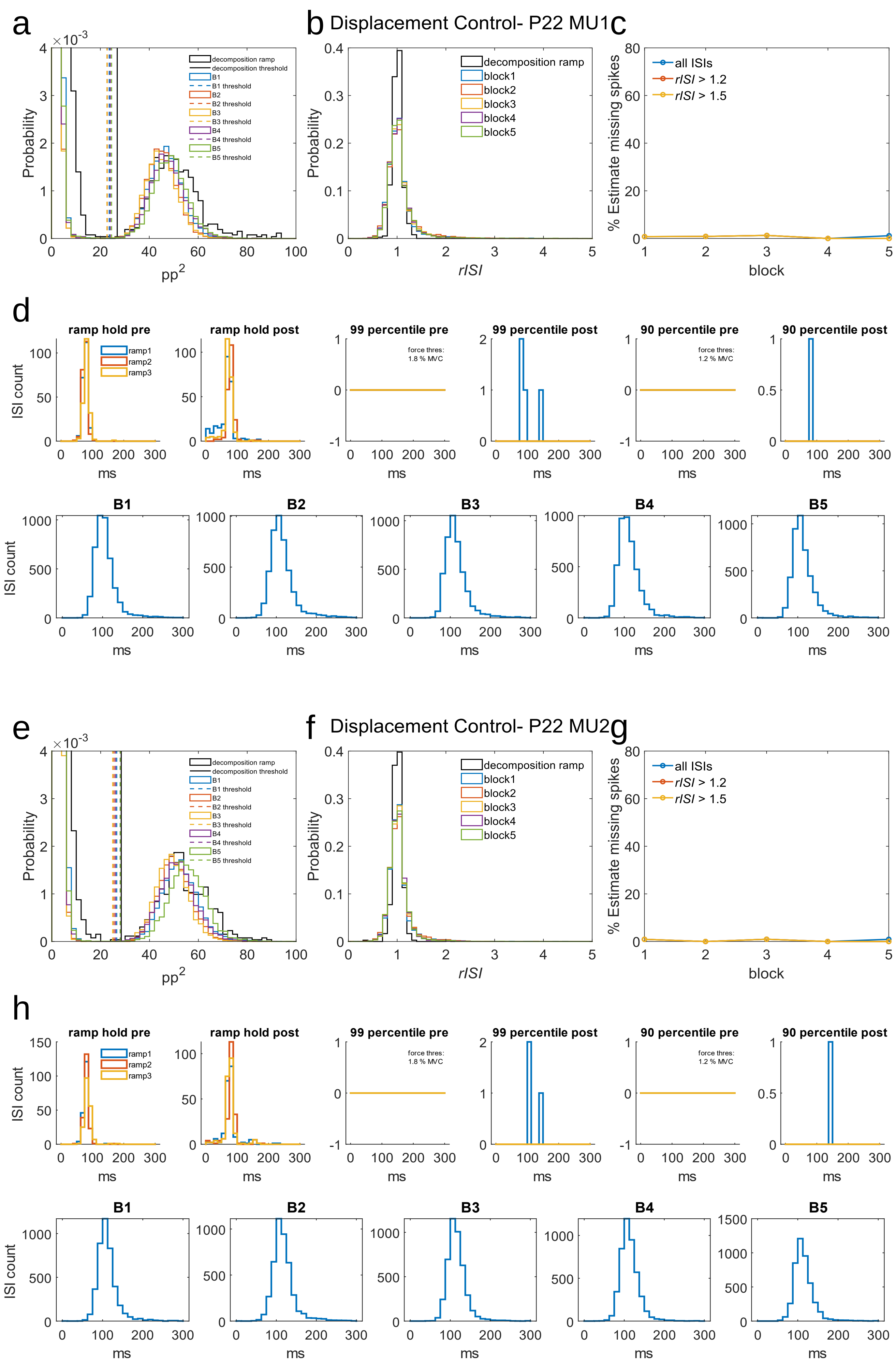


Supplementary Figure S20

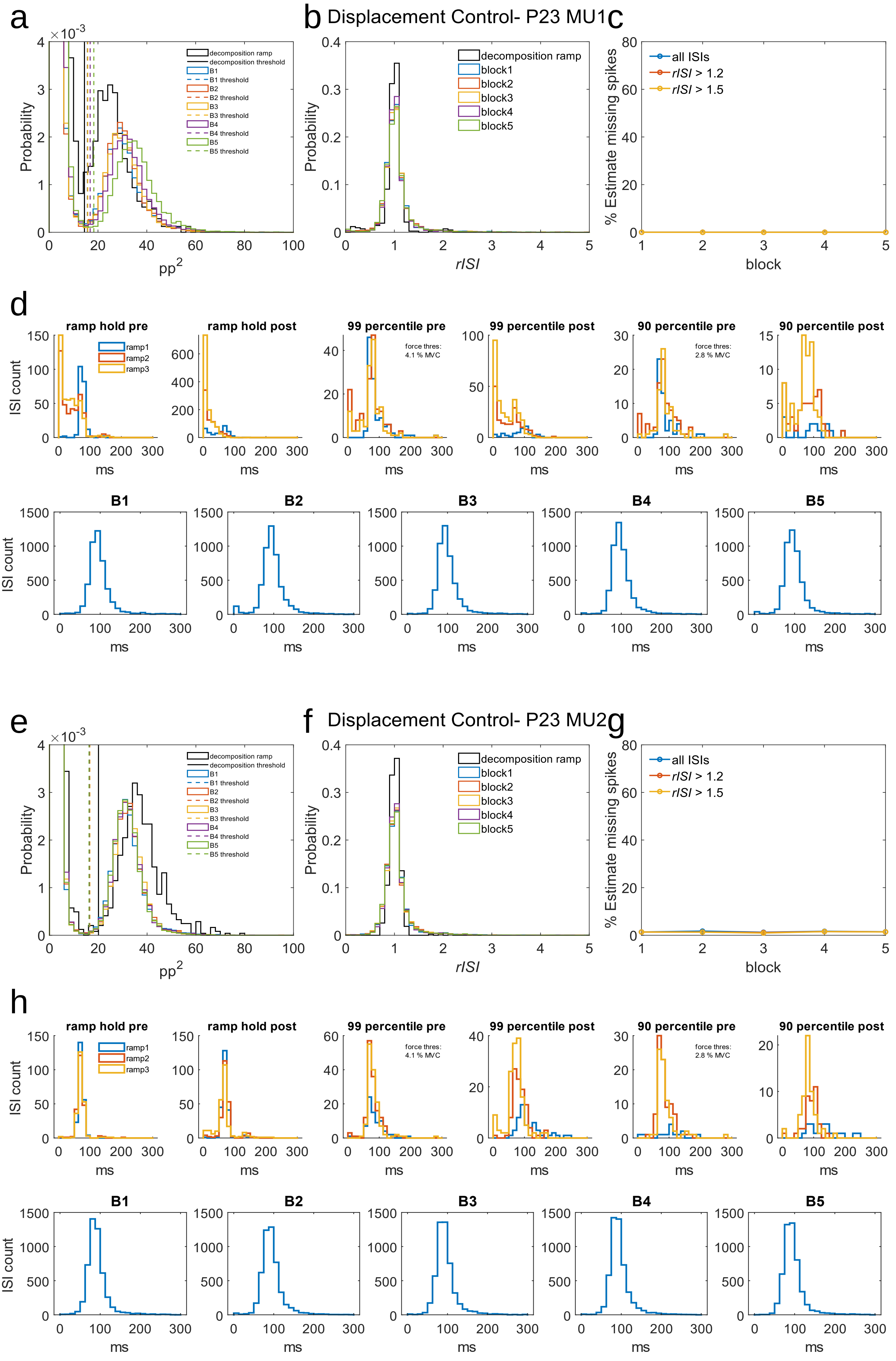


Supplementary Figure S21

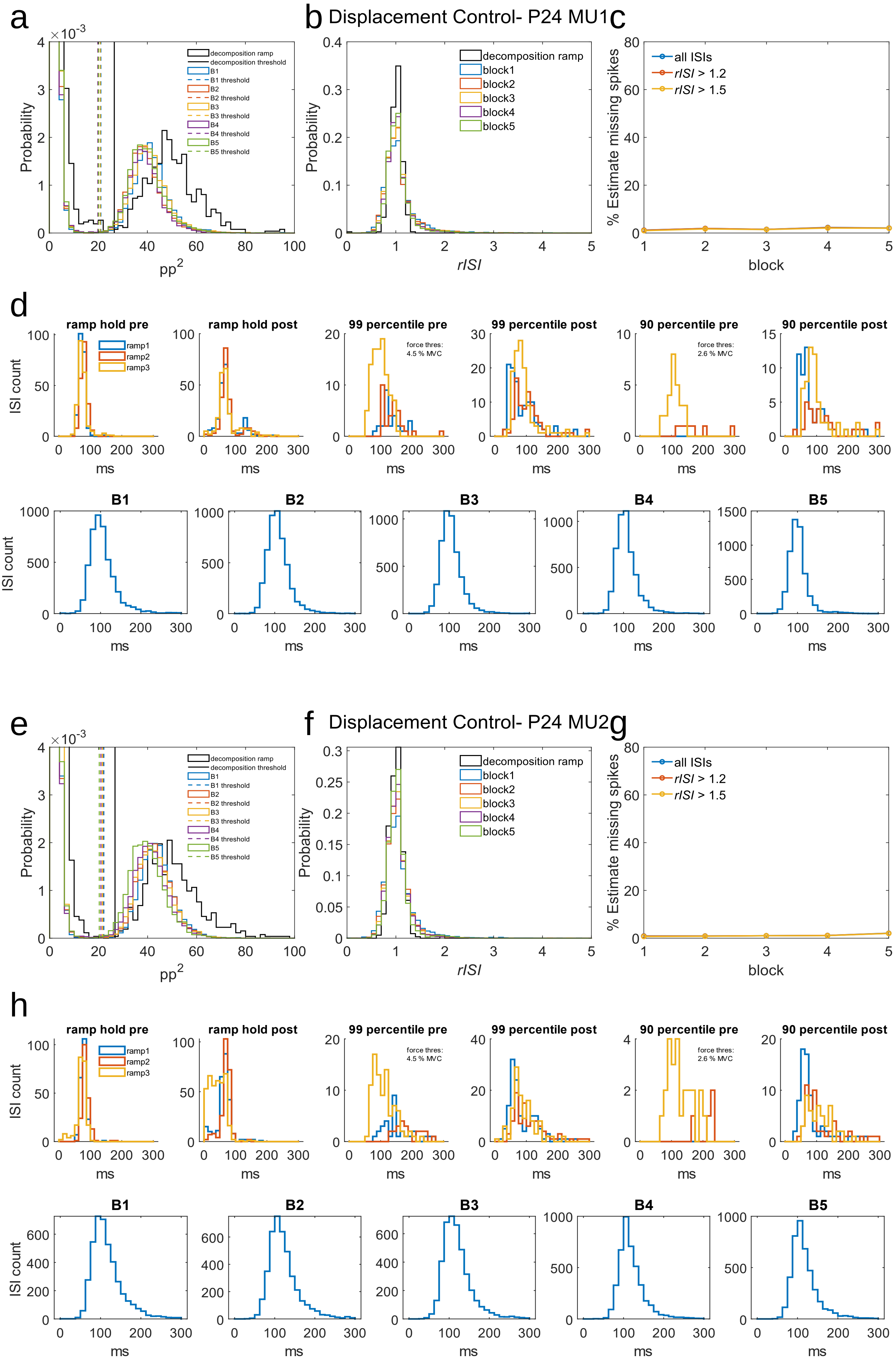


Supplementary Figure S22

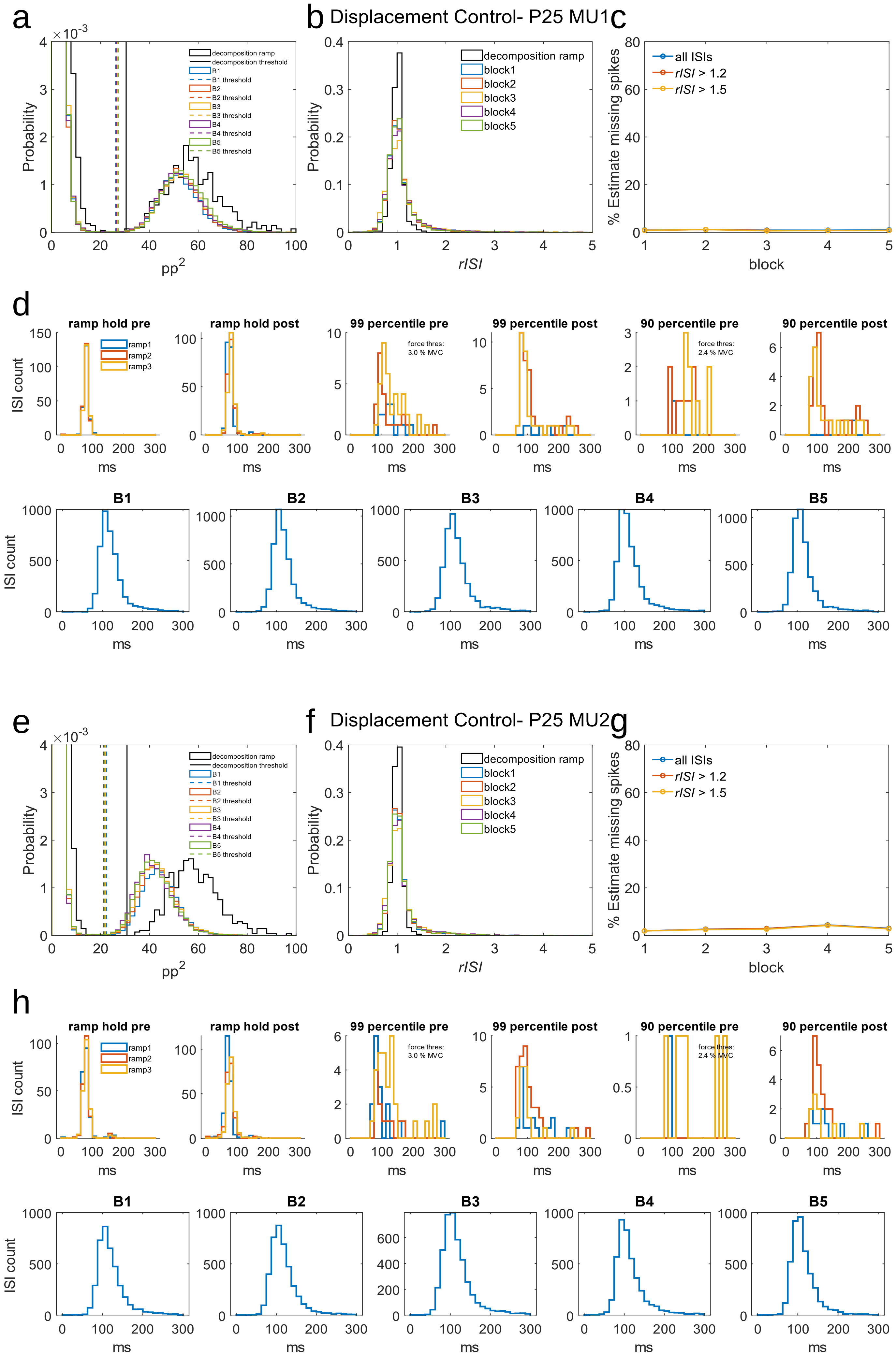


Supplementary Figure S23

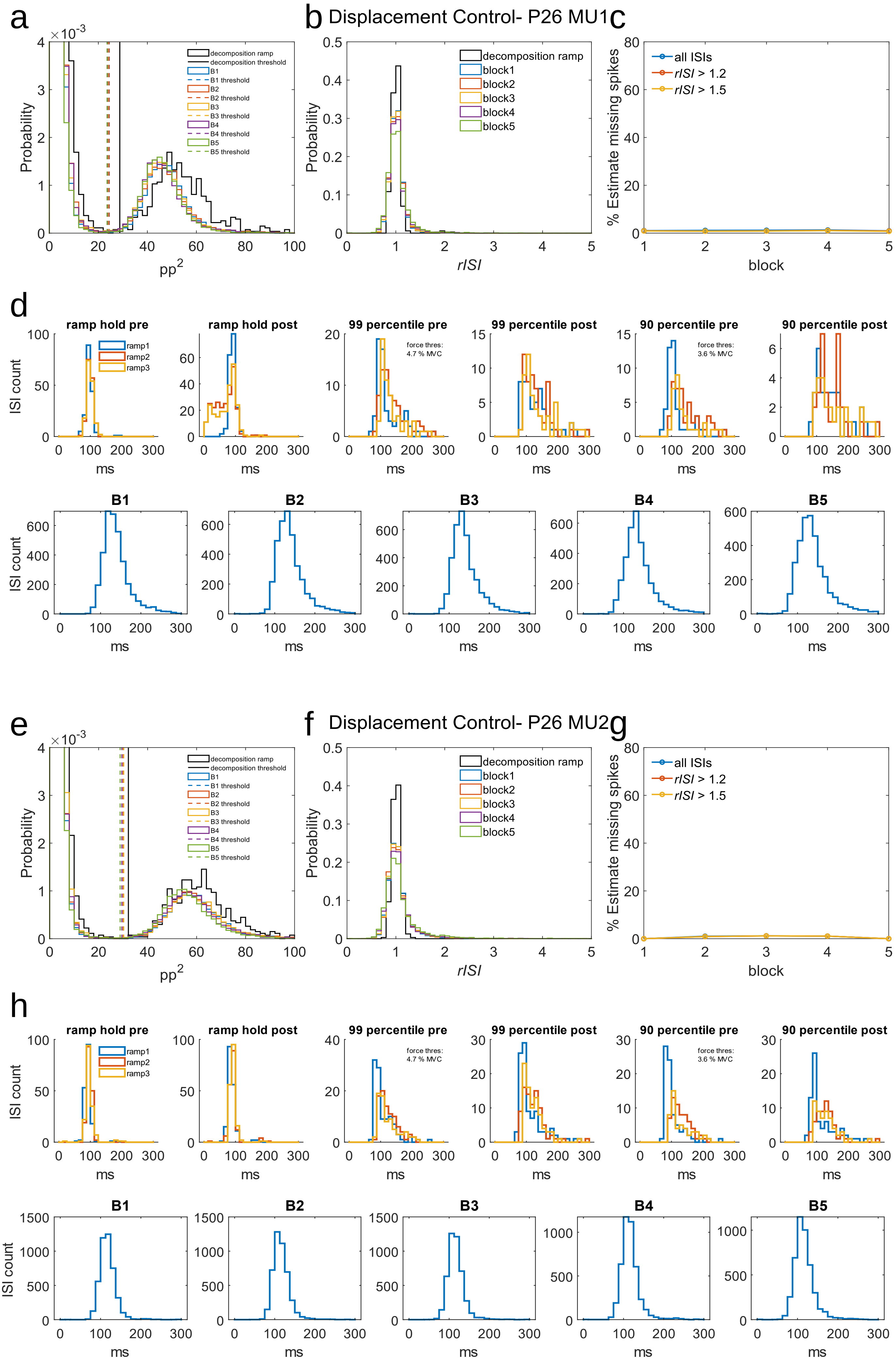


Supplementary Figure S24

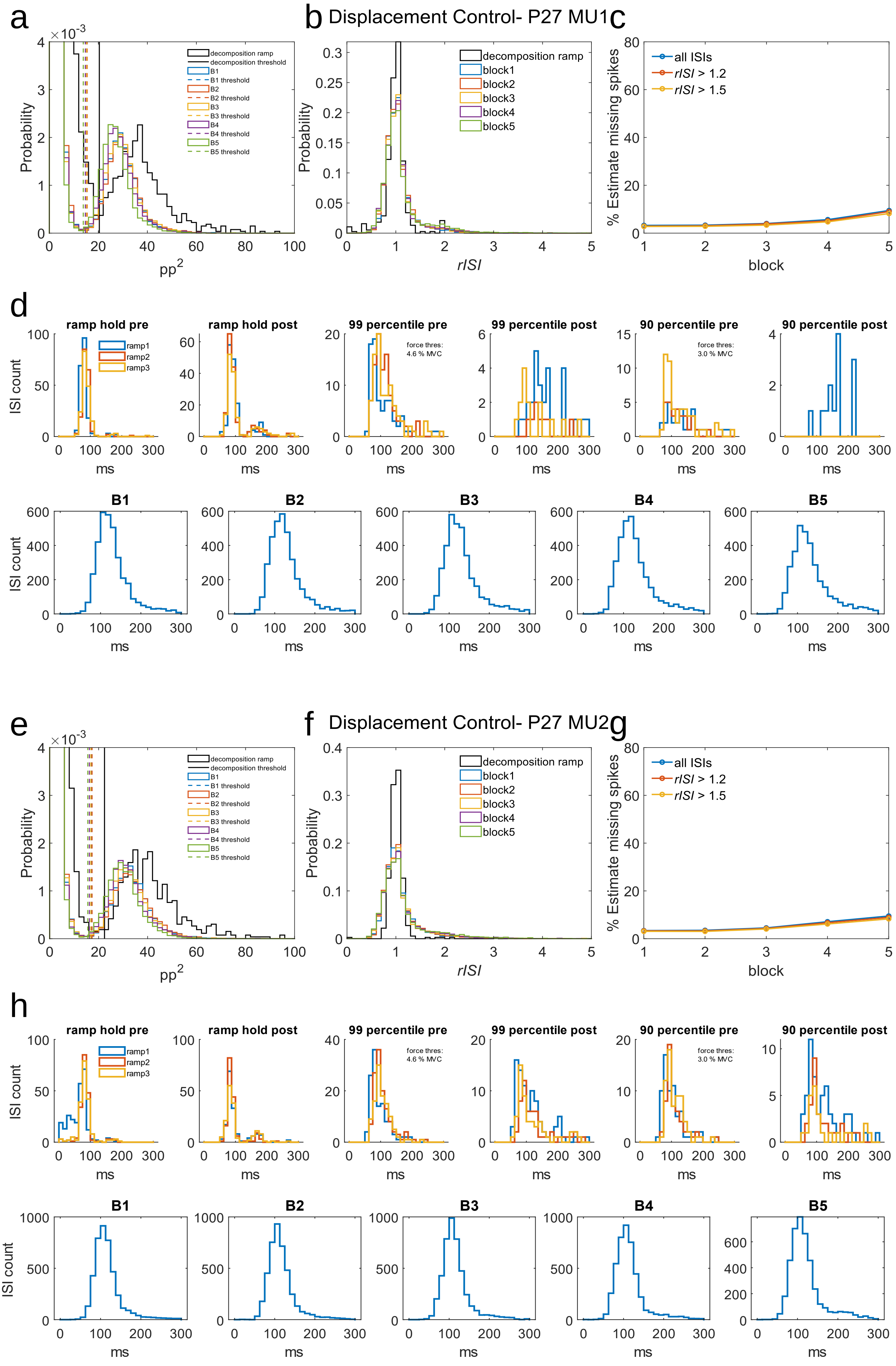


Supplementary Figure S25

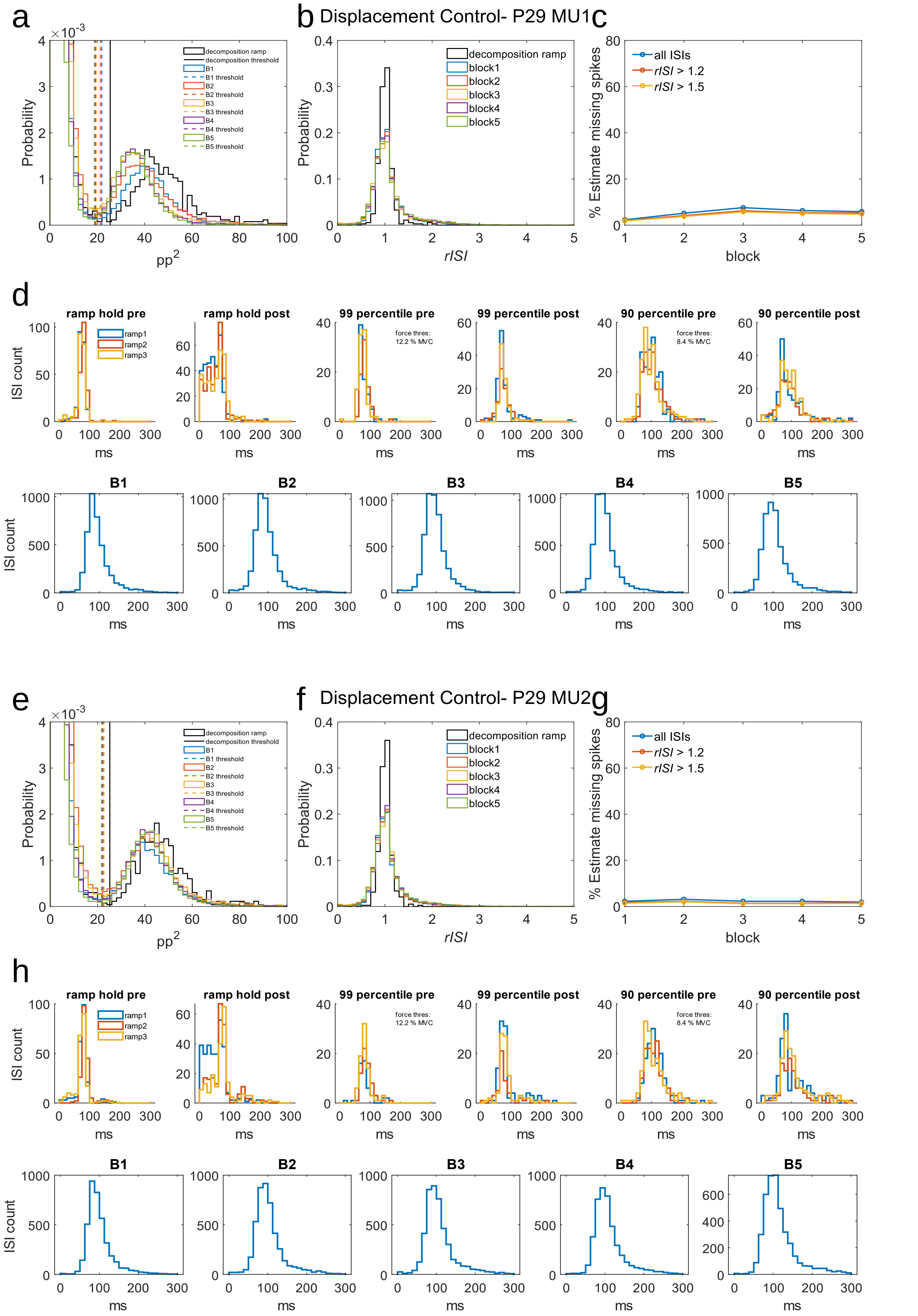


Supplementary Figure S26

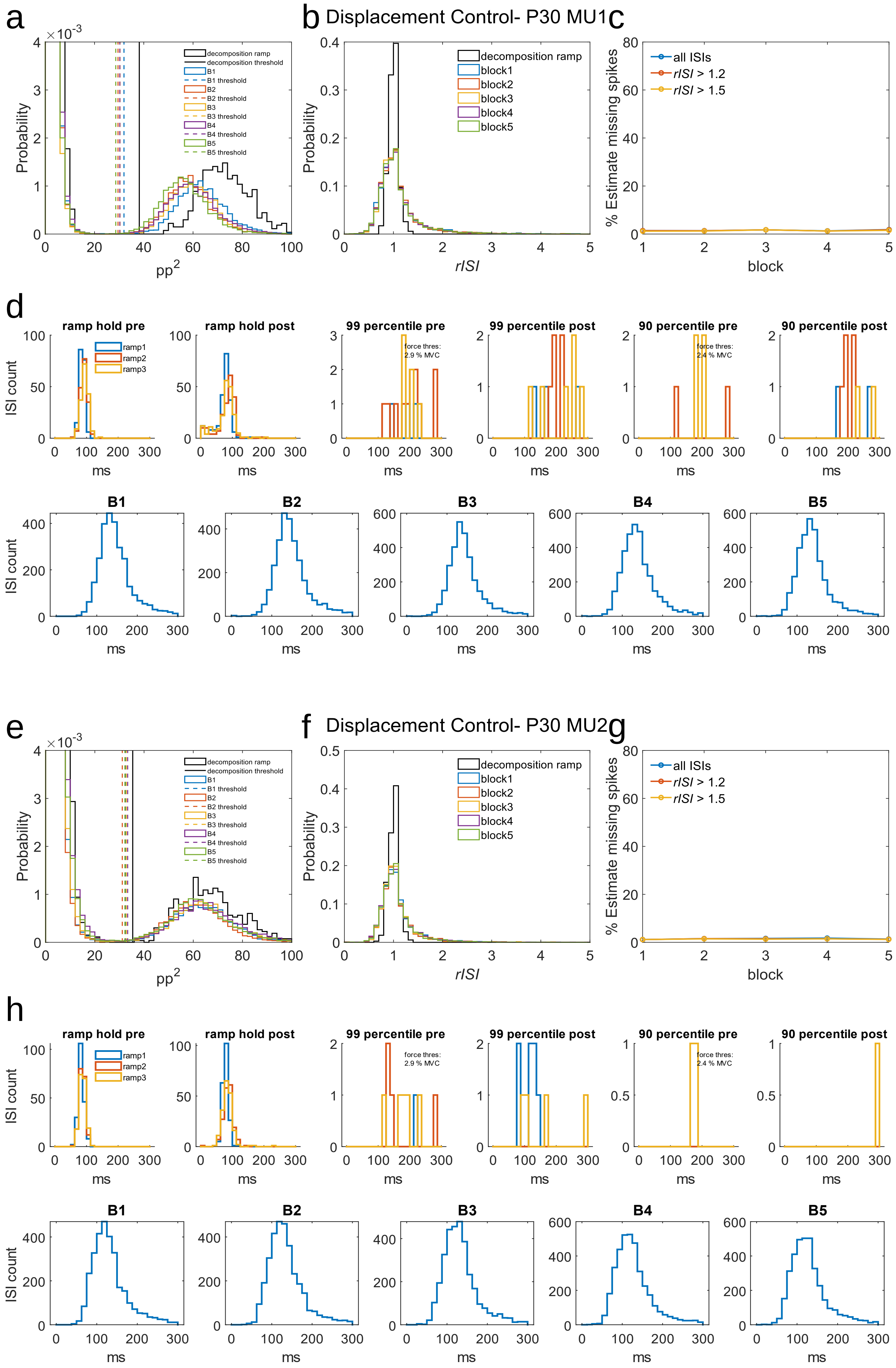


Supplementary Figure S27

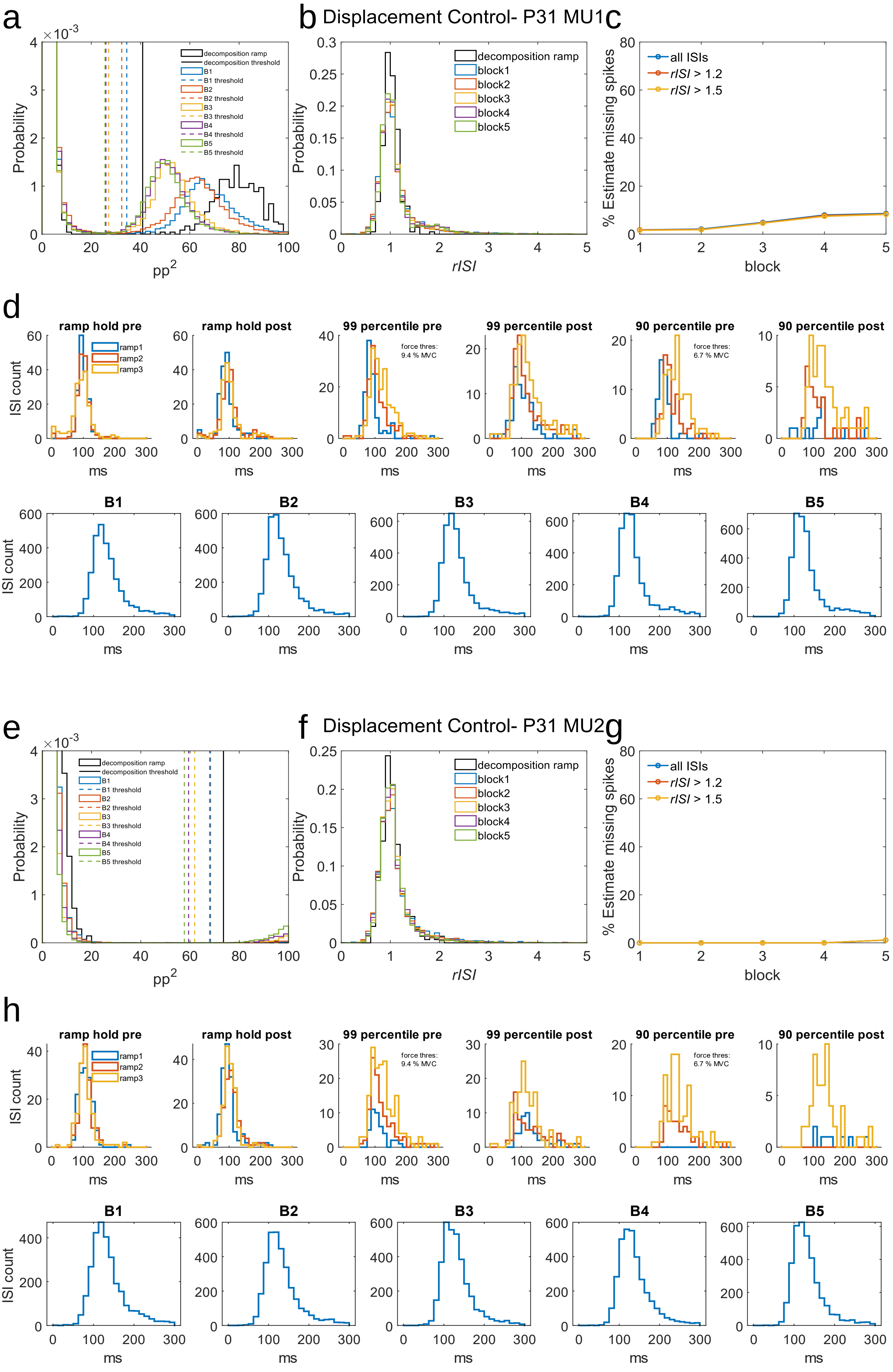


Supplementary Figure S28

### Excluded participants-difference control

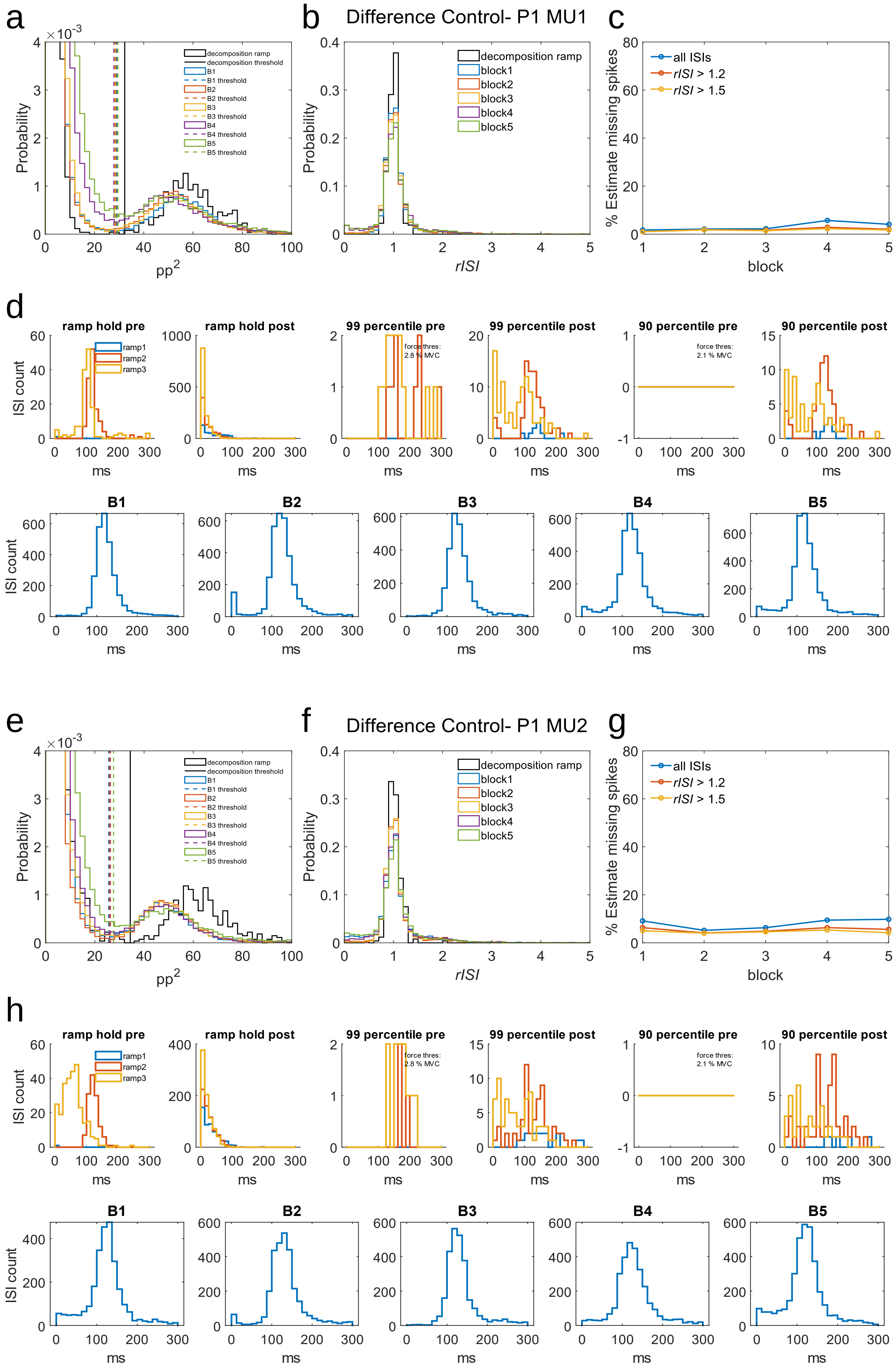


Supplementary Figure S29

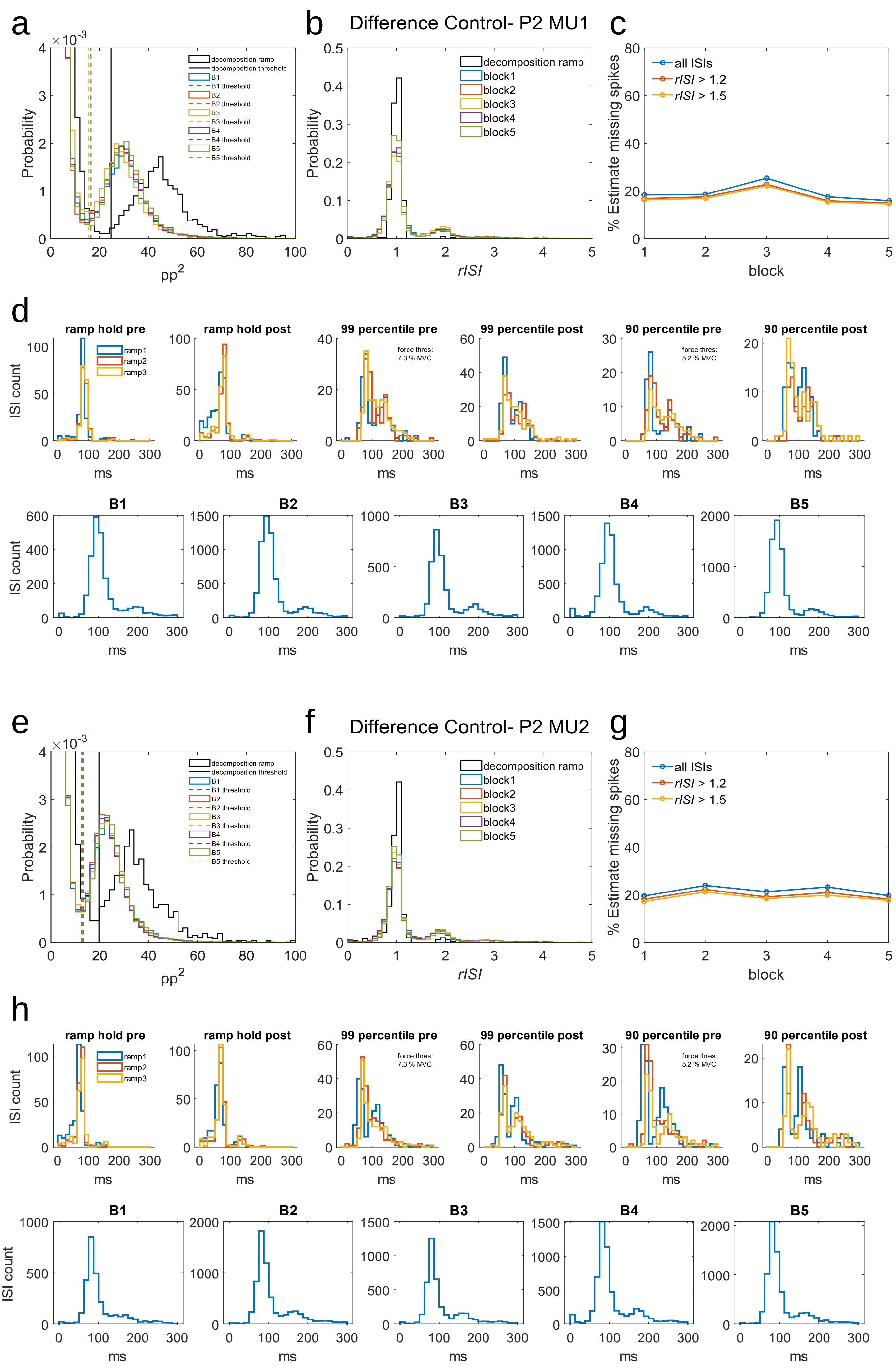


Supplementary Figure S30

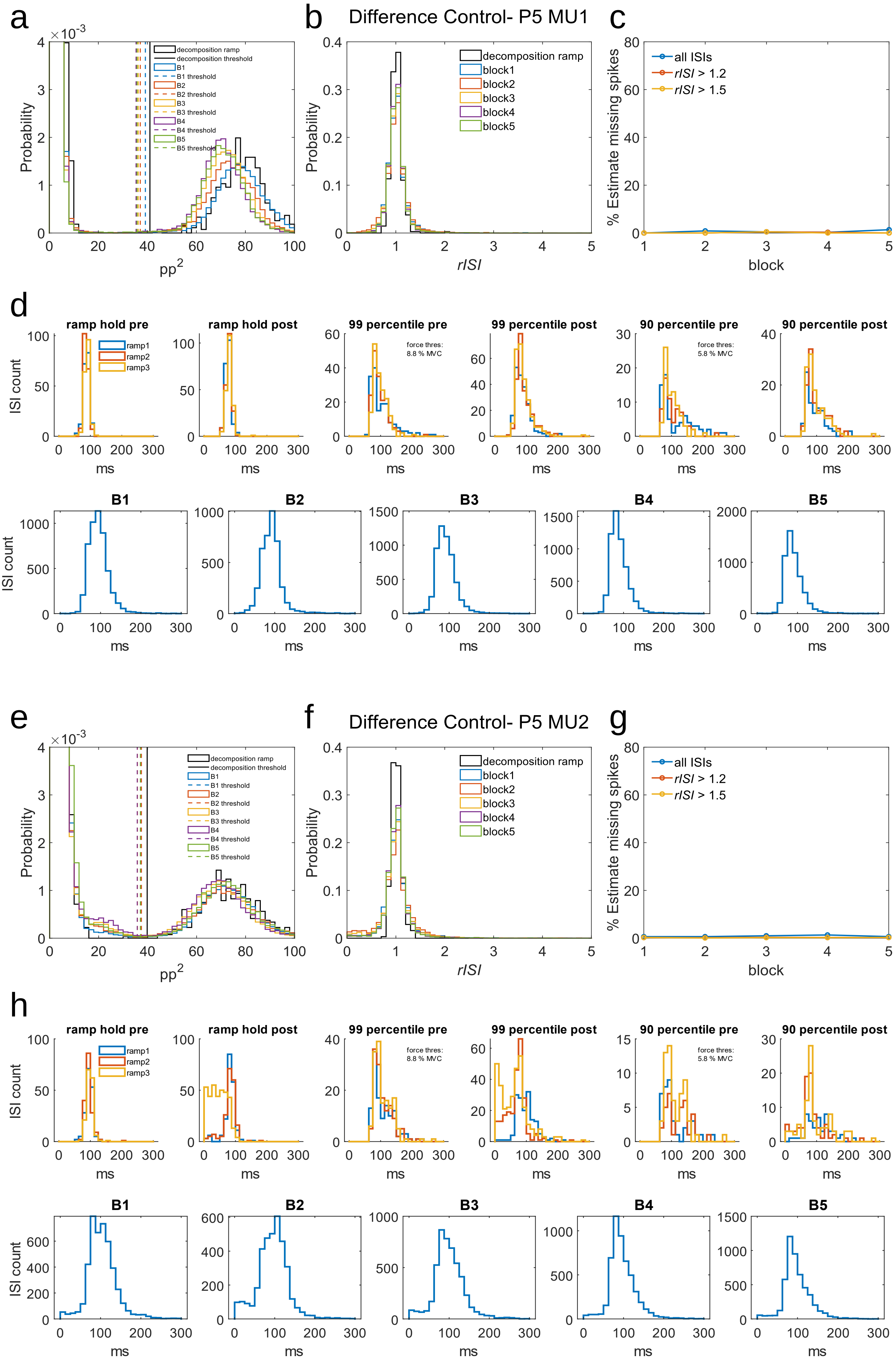


Supplementary Figure S31

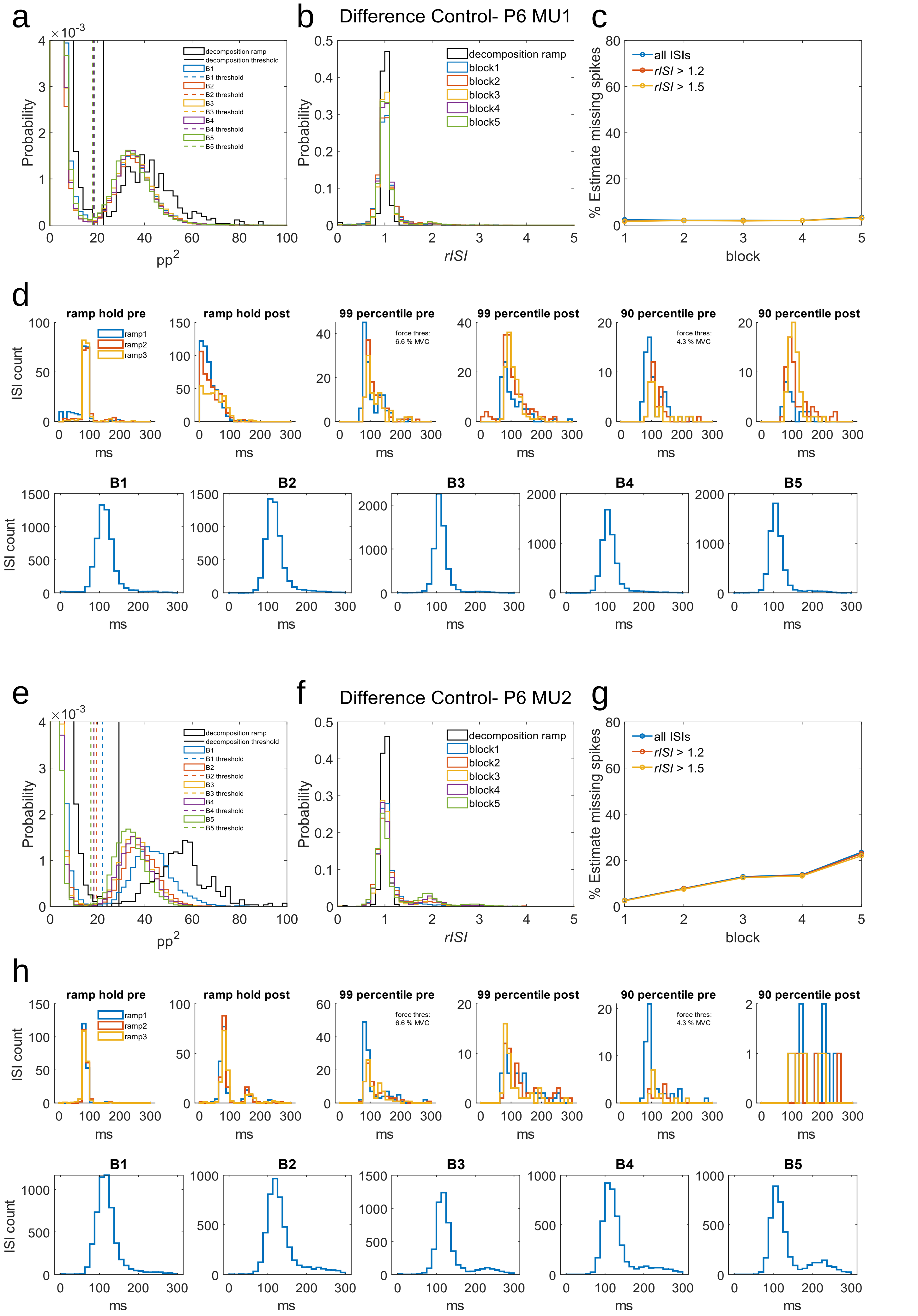


Supplementary Figure S32

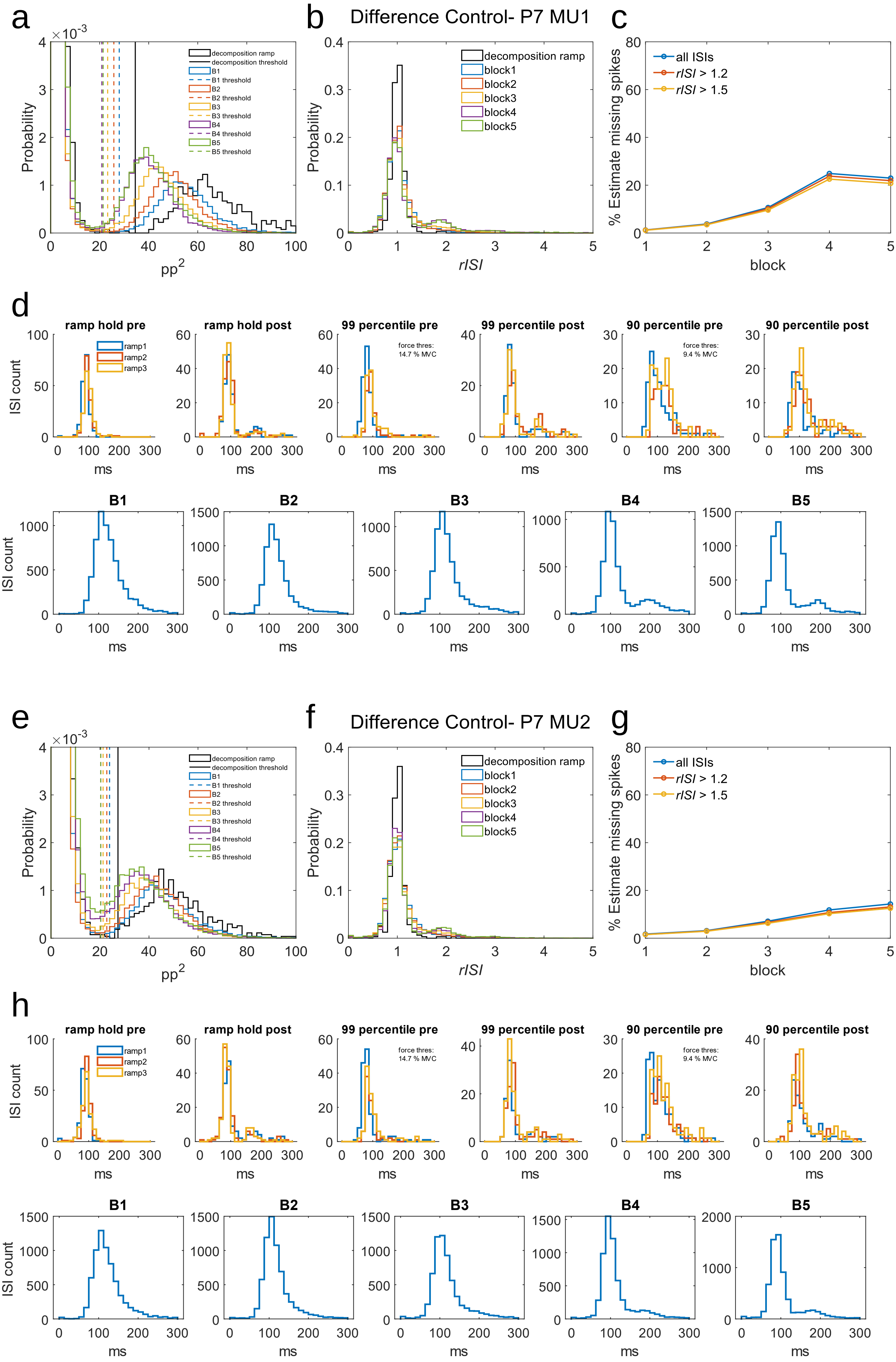


Supplementary Figure S33

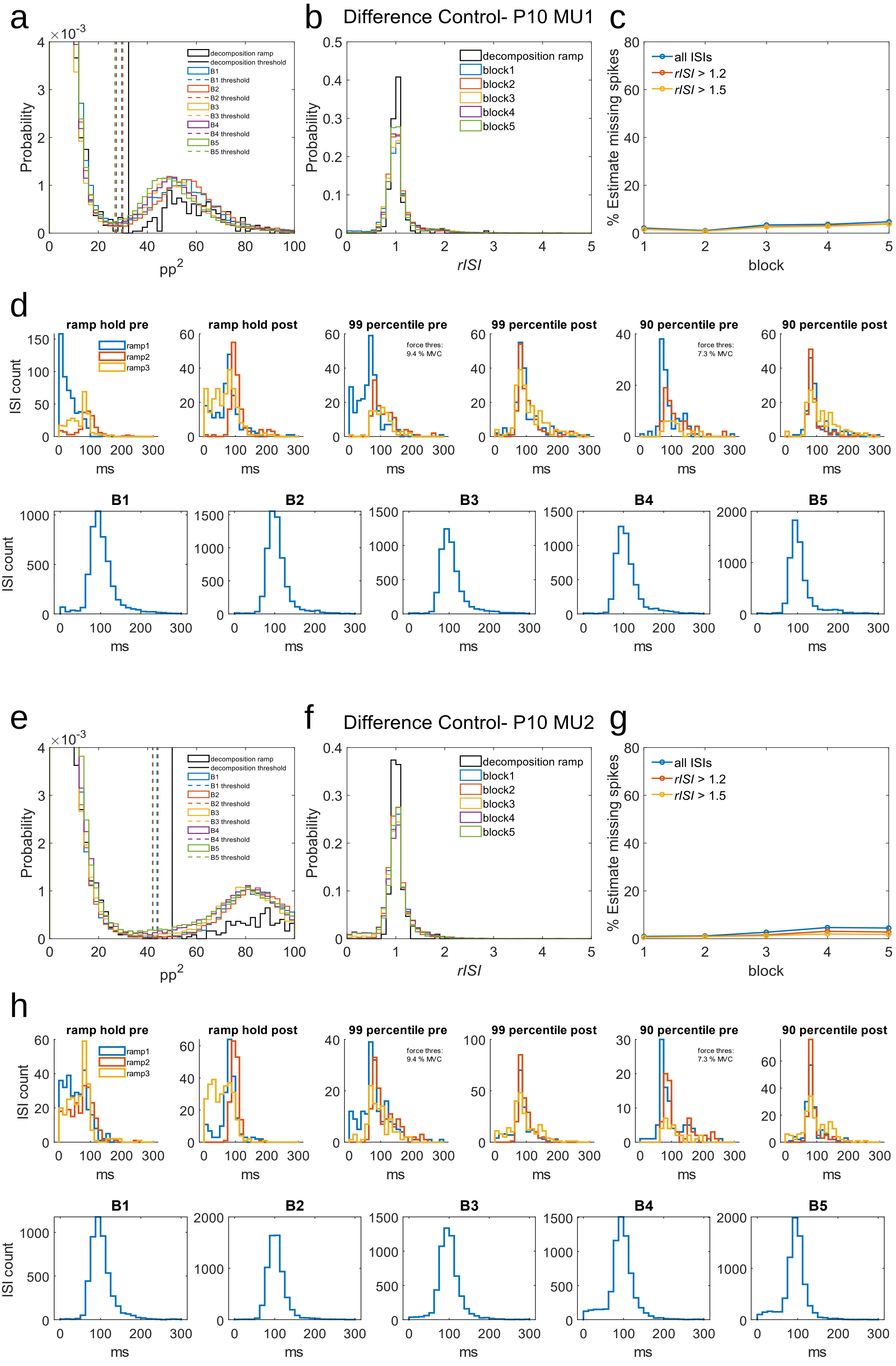


Supplementary Figure S34

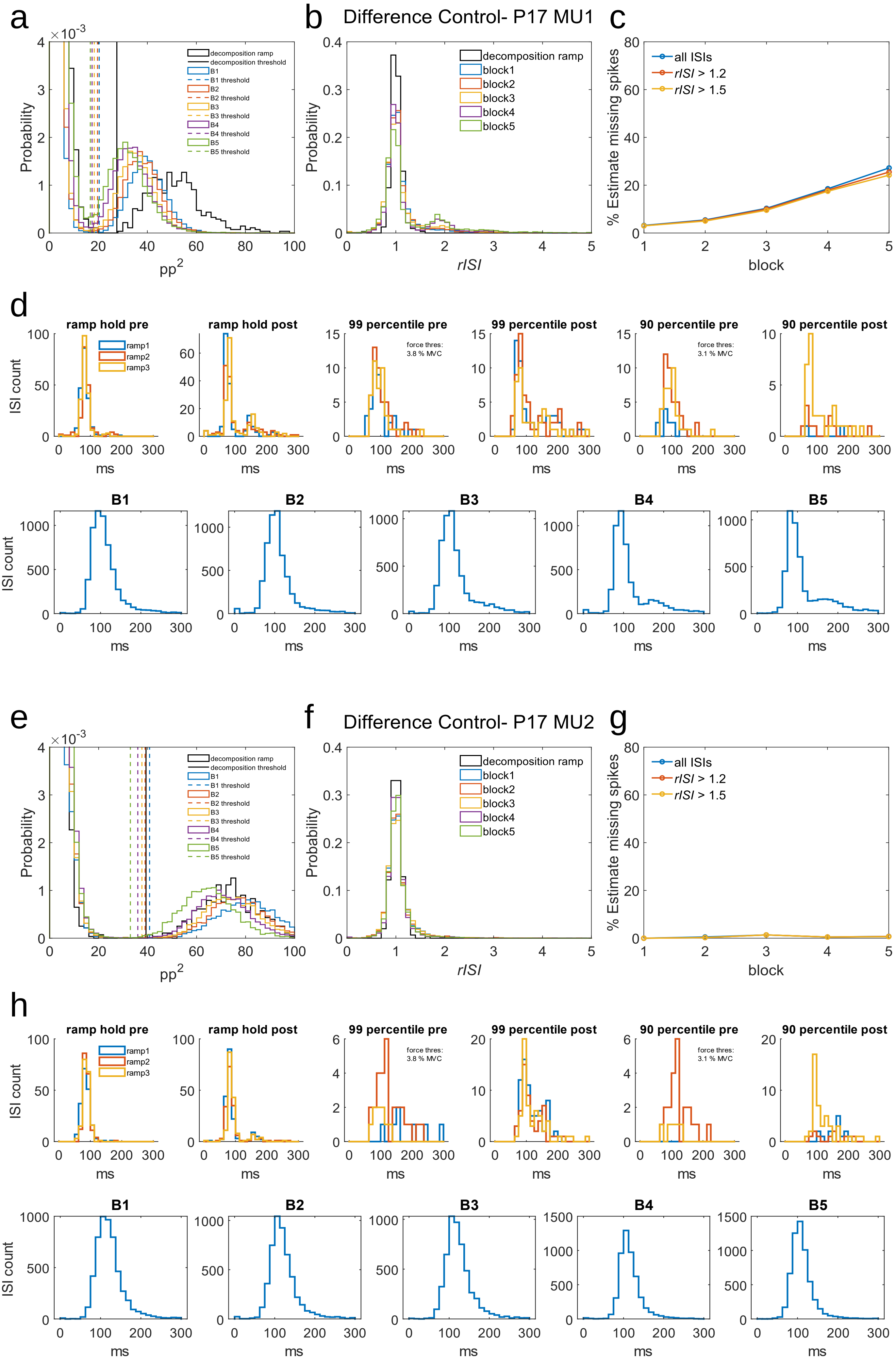


Supplementary Figure S35

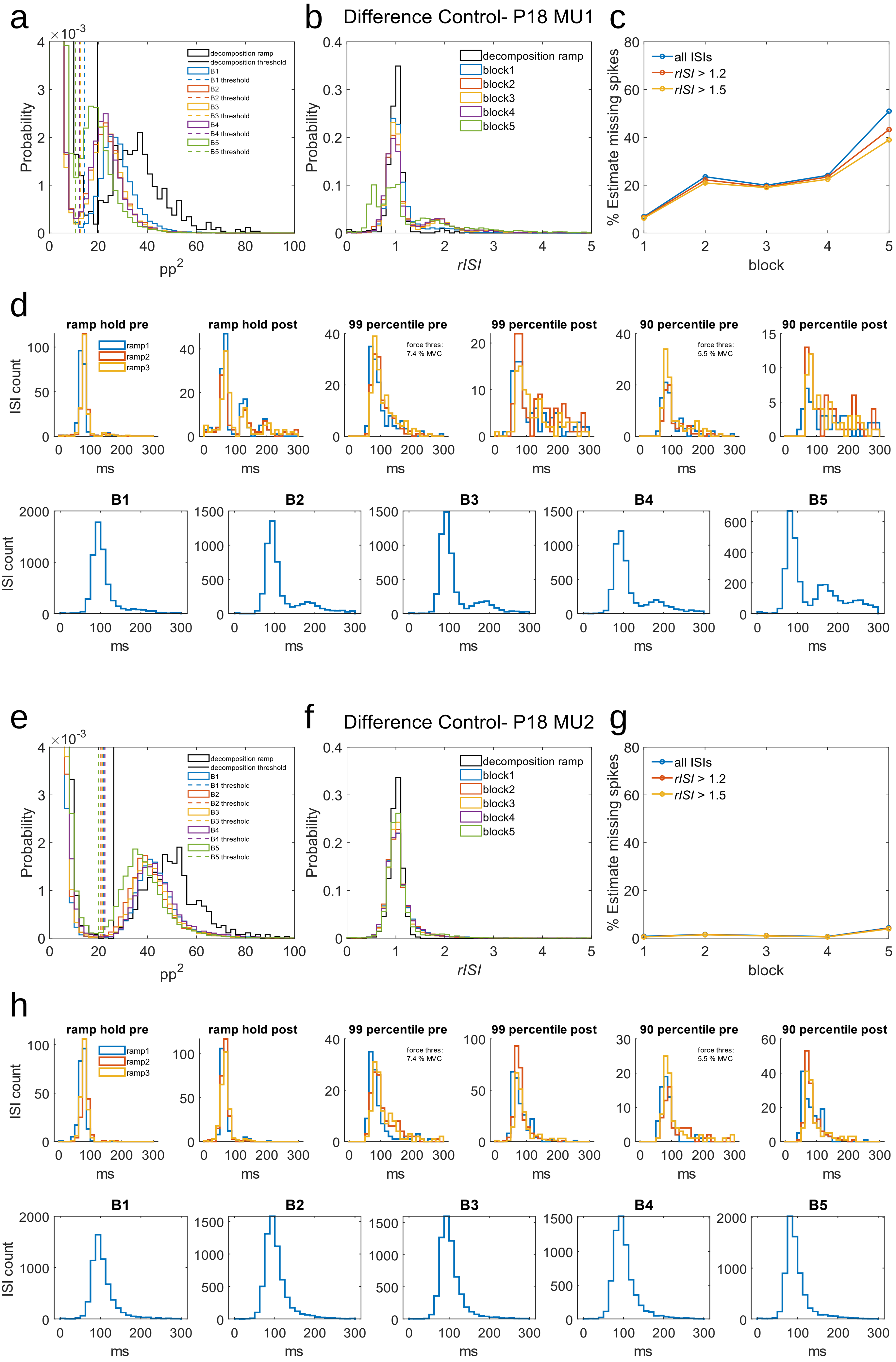


Supplementary Figure S36

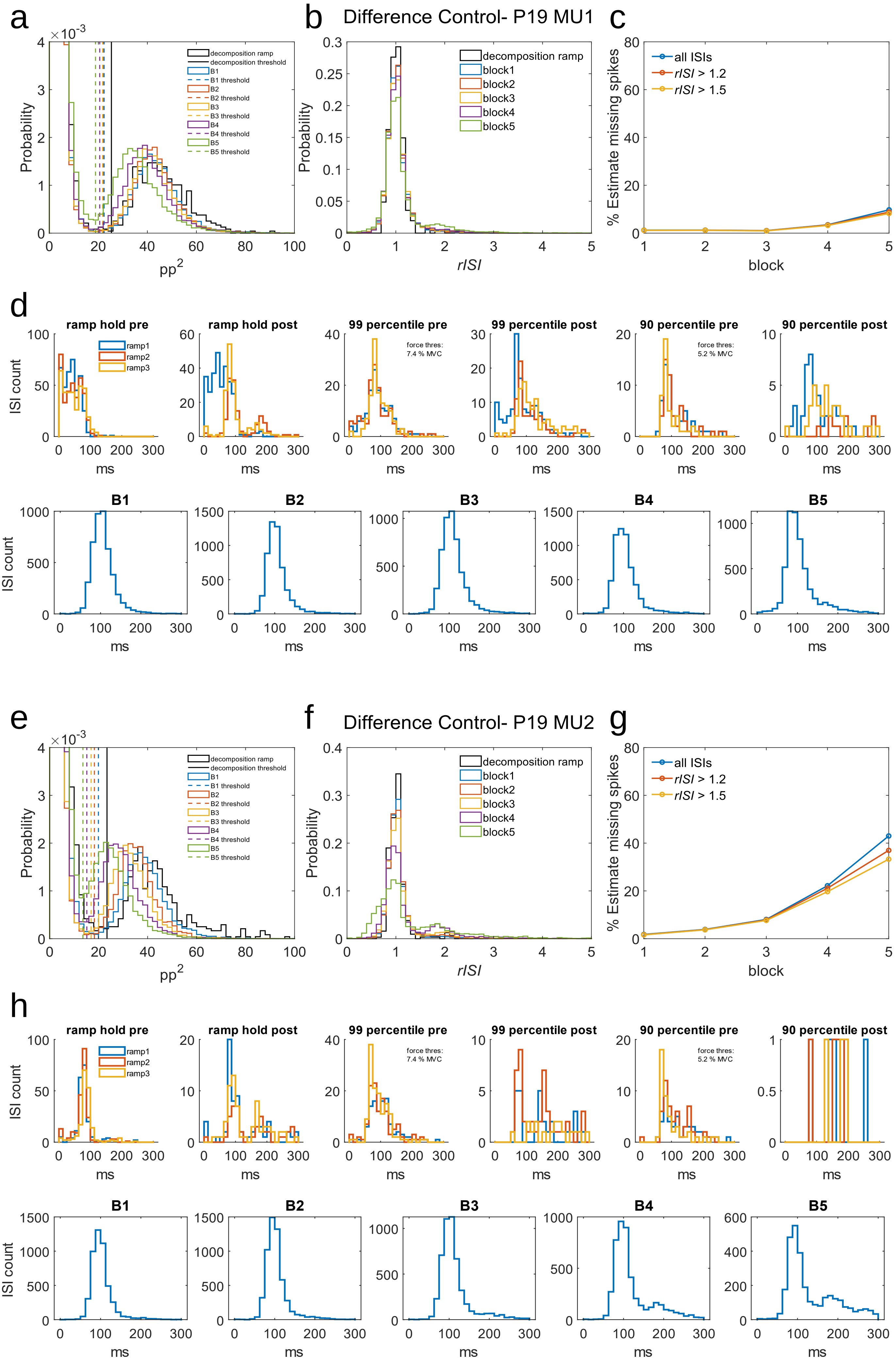


Supplementary Figure S37

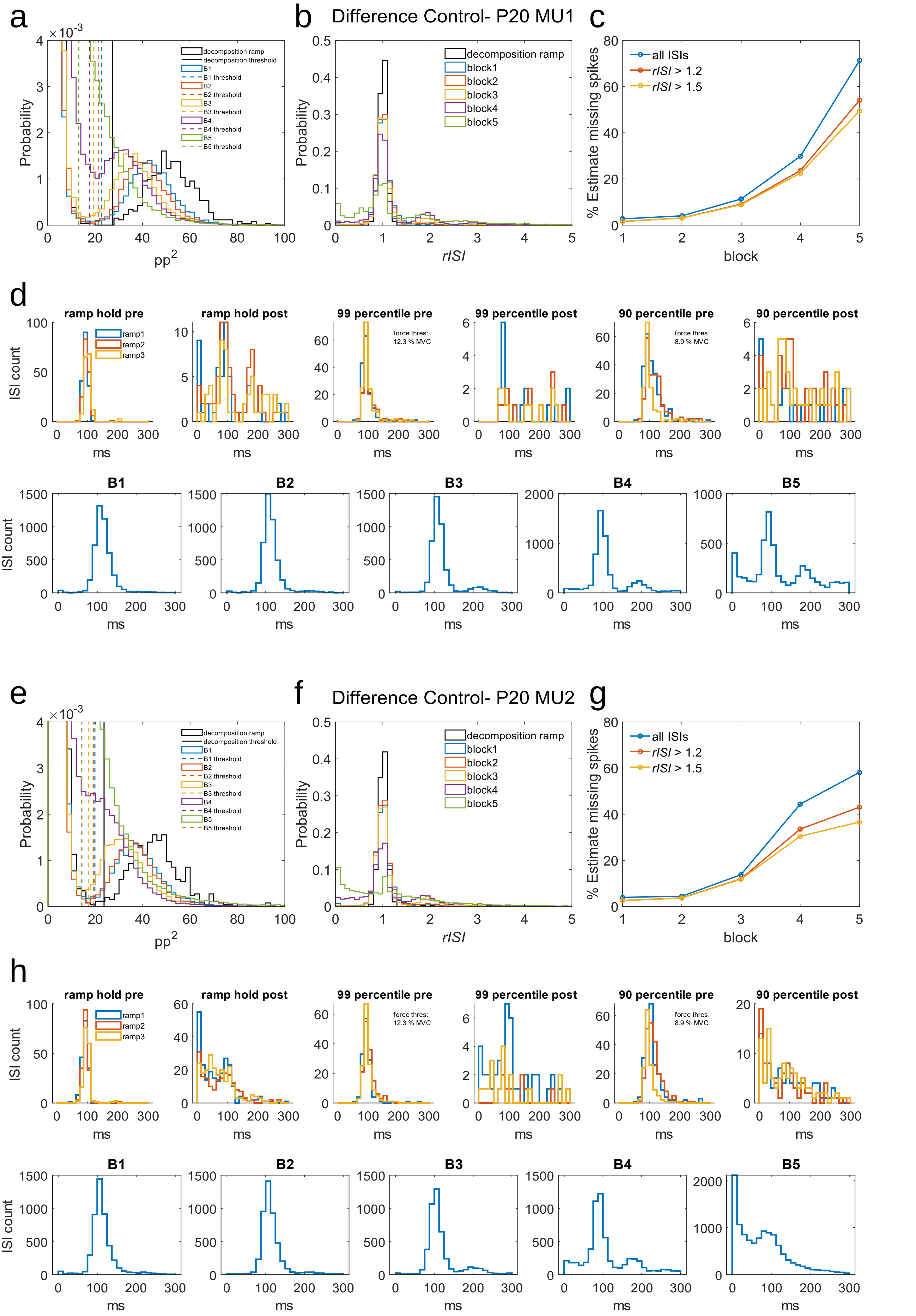


Supplementary Figure S38

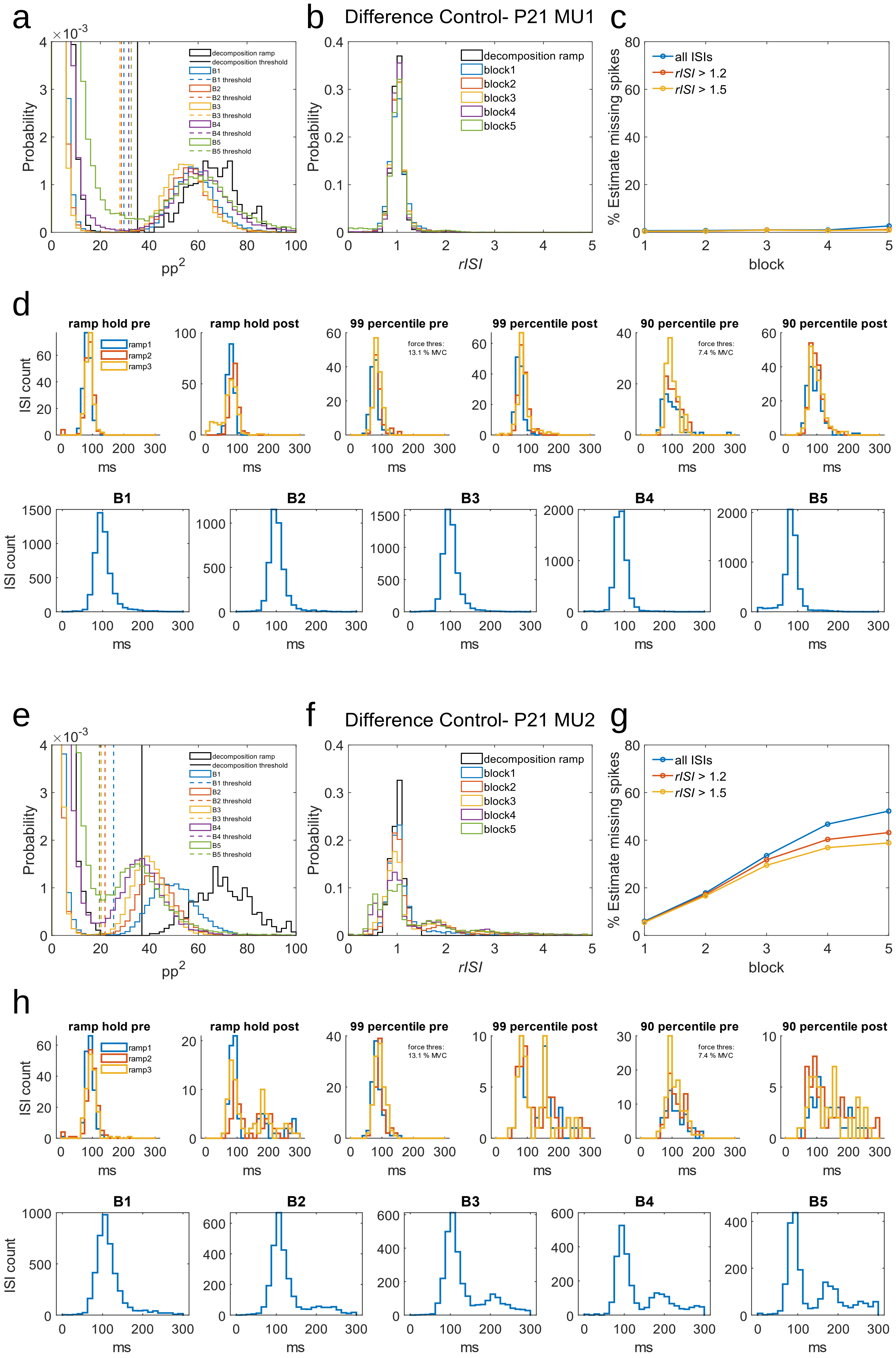


Supplementary Figure S39

### Excluded participants- displacement control

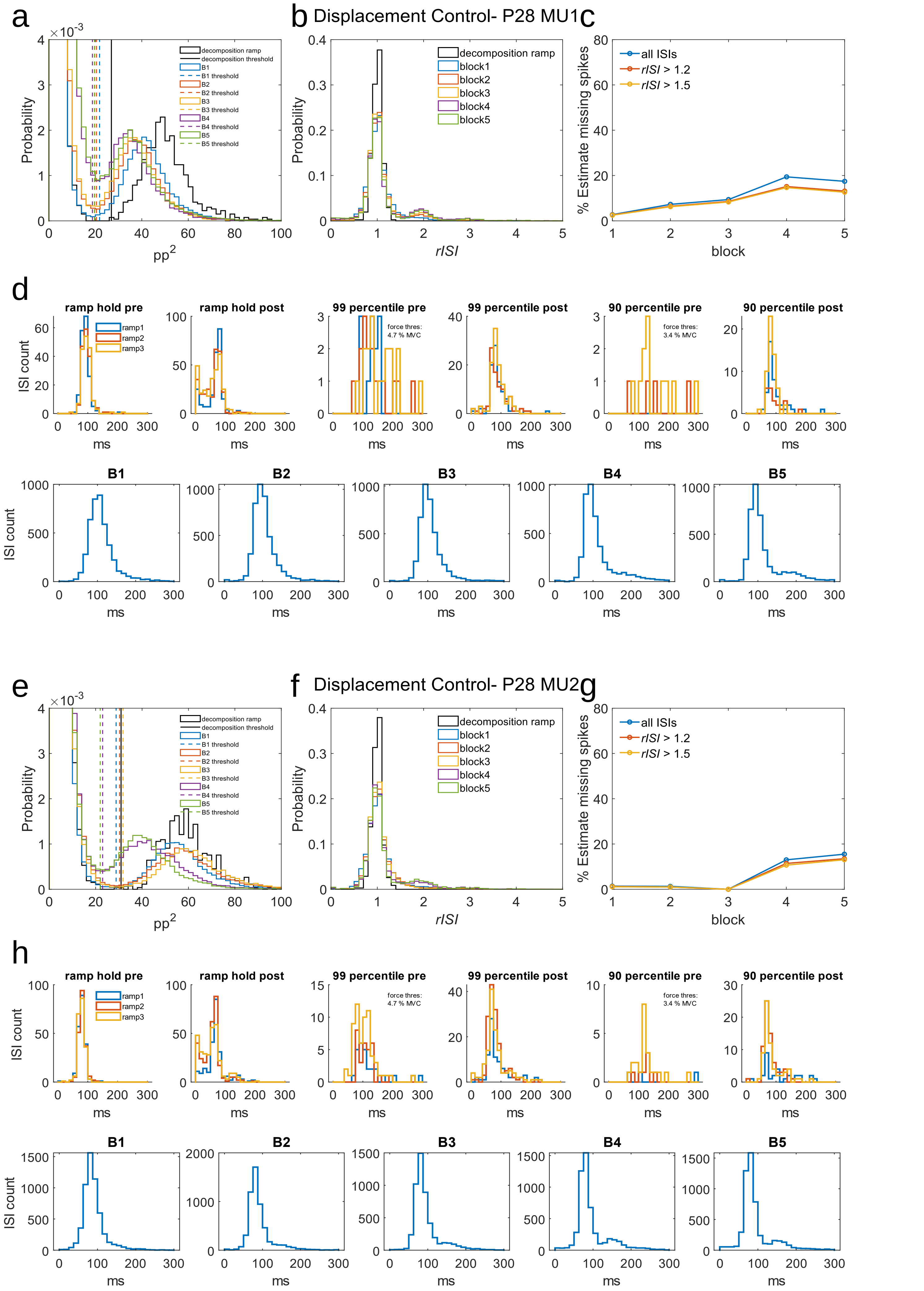


Supplementary Figure S40
